# Changes in regional white matter volumetry and microstructure during the post-adolescence period: a cross-sectional study of a cohort of 1,713 university students

**DOI:** 10.1101/2021.04.13.439695

**Authors:** Ami Tsuchida, Alexandre Laurent, Fabrice Crivello, Laurent Petit, Antonietta Pepe, Naka Beguedou, Stephanie Debette, Christophe Tzourio, Bernard Mazoyer

## Abstract

Human brain white matter undergoes a protracted maturation that continues well into adulthood. Recent advances in diffusion-weighted imaging (DWI) methods allow detailed characterizations of the microstructural architecture of white matter, and they are increasingly utilised to study white matter changes during development and ageing. However, relatively little is known about the late maturational changes in the microstructural architecture of white matter during post-adolescence. Here we report on regional changes in white matter volume and microstructure in young adults undergoing university-level education. As part of the MRi-Share multi-modal brain MRI database, multi-shell, high angular resolution DWI data were acquired in a unique sample of 1,713 university students aged 18 to 26. We assessed the age and sex dependence, as well as hemispheric asymmetry of diffusion metrics derived from diffusion tensor imaging (DTI) and neurite orientation dispersion and density imaging (NODDI), in the white matter regions as defined in the John Hopkins University (JHU) white matter labels atlas. We demonstrate that while regional white matter volume is relatively stable over the age range of our sample, the white matter microstructural properties show clear age-related variations. Globally, it is characterised by a robust increase in neurite density index (NDI), and to a lesser extent, orientation dispersion index (ODI). These changes are accompanied by a decrease in diffusivity. In contrast, there is minimal age-related variation in fractional anisotropy. There are regional variations in these microstructural changes: some tracts, most notably cingulum bundles, show a strong age-related increase in NDI coupled with decreases in radial and mean diffusivity, while others, mainly cortico-spinal projection tracts, primarily show an ODI increase and axial diffusivity decrease. These age-related variations are not different between males and females, but males show higher NDI and ODI and lower diffusivity than females across many tracts. We also report a robust hemispheric asymmetry in both the volume and microstructural properties in many regions. These findings emphasise the complexity of changes in white matter structure occurring in this critical period of late maturation in early adulthood.

## 1. Introduction

Early adulthood is characterised by significant changes in lifestyle and behaviour for many, when individuals explore their identity and various life possibilities to become fully independent. For some, it involves the attainment of higher education and training to acquire new skills and knowledge necessary for their planned vocation. Although the most dramatic development in the human brain takes place earlier in life, with the total brain volume reaching 90 % of the adult volume by the age of 5 years (Dekaban, 1978; Lenroot and Giedd, 2006), both global and regional changes in brain structure and function persist throughout childhood and adolescence, and some of the maturational changes continue well into adulthood (Dumontheil, 2016). In particular, the white matter (WM) of the brain shows a protracted course of development, with its total volume continuing to increase up to the fourth or fifth decade of life (Walhovd et al., 2011; Lebel et al., 2012). The development of WM microstructure is also sensitive to common life experiences in young adults, including exposure to alcohol and tobacco, and other recreational drugs (Bava et al., 2013; Gogliettino et al., 2016; Silveri et al., 2016), changes in sleep patterns (Telzer et al., 2015; Elvsåshagen et al., 2015), and intensive motor and cognitive training (Scholz et al., 2009; Lövdén et al., 2010; Mackey et al., 2012; Schlegel et al., 2012; Lakhani et al., 2016). Detailed characterization of the late maturational processes of the WM in young adults is crucial for elucidating how the learning and other life experiences may shape the structural and functional organization of the brain through their impact on the brain wiring. Understanding the normative development during this period may also shed light on the vulnerability of this particular period in life to various neuropsychiatric disorders, such as substance abuse, mood and anxiety disorders (Kessler et al., 2007).

What we know about the normative WM development primarily comes from noninvasive neuroimaging of typically developing individuals with magnetic resonance imaging (MRI). In addition to the macro-structural changes that can be measured with T1-weighted images, diffusion-weighted imaging (DWI) methods allow detailed characterizations of the WM microstructural properties. Over the past two decades, studies using DWI have provided much insight into the WM microstructural changes during development (reviewed in Tamnes et al., 2018; Lebel and Deoni, 2018; Lebel et al., 2019). The majority of these studies quantify DWI through a diffusion tensor imaging (DTI) model that characterises the direction and magnitude of diffusion of tissue water molecules as a single tensor in each voxel (Tournier et al., 2011). Most commonly, fractional anisotropy (FA), which measures the degree of diffusion directionality, is used to quantify maturational changes, with an increase in FA attributed to myelination and increased axonal size and/or packing. Other DTI measures include axial and radial diffusivity (AD/RD), which represent diffusion along the longest and shortest axis, respectively, of the tensor modelled in each voxel, and mean diffusivity (MD), representing the average magnitude of diffusion. Across studies, FA increases and overall decreases in diffusivity with increasing age are observed in most WM regions through childhood and adolescence (e.g. Bonekamp et al., 2007; Lebel et al., 2008; Giorgio et al., 2010; Tamnes et al., 2010; Lebel and Beaulieu, 2011; Brouwer et al., 2012; Simmonds et al., 2014; Pohl et al., 2016). In a large-scale, multi-cohort study, we have recently demonstrated that such changes continue up to early to mid-adulthood (Beaudet et al., 2020).

However, being a “signal” based model, the DTI model only describes the diffusion process in each voxel and does not attempt to delineate signals attributable to different biological tissue components (Ferizi et al., 2017). Thus, changes in DTI metrics only indicate alterations in magnitude or directionality of diffusivity, and different biological processes that affect diffusion properties of the tissue cannot be distinguished (Jones et al., 2013). More concretely, FA can be increased due to myelination and/or increased axonal packing but would decrease with increasing fibre population complexity. In contrast, “tissue” based models attempt to estimate the components of underlying tissue, typically using DWI acquisitions with multiple *b*-values, and likely provide more biologically specific insights (Alexander et al., 2019). One such model is neurite orientation dispersion and density imaging (NODDI), which models three tissue compartments (intra- and extra-cellular, and cerebrospinal fluid). It estimates separate indices for neurite density (neurite density index, NDI) and fibre orientation complexity (orientation dispersion index, ODI), together with the isotropic volume fraction (i.e. cerebrospinal fluid compartment, IsoVF) (Zhang et al., 2012). Several recent studies have used NODDI to examine developmental changes in the WM microstructural properties through infancy (Dean et al., 2017; Jelescu et al., 2015), childhood to adolescence (Genc et al., 2017; Mah et al., 2017; Dimond et al., 2020; Lynch et al., 2020). These studies have indicated an age-related increase in NDI, with very little change observed in ODI in the first two decades of life (Mah et al., 2017; Dimond et al., 2020; Lynch et al., 2020), although studies covering a wider age range indicate that ODI in many WM tracts starts to increase in early adulthood (Chang et al., 2015; Slater et al., 2019). Nevertheless, a large-scale study focusing on the period of early adulthood to detail the late maturational changes in regional WM properties is still lacking.

In the present study, we characterise variations in WM-related image-derived phenotypes (IDP), including regional volumes and microstructural properties measured using both DTI and NODDI, in a large cross-sectional cohort of young adults undergoing university-level education. As part of the MRi-Share multi-modal brain MRI database (Tsuchida et al., 2020), multi-shell DWI data were acquired, together with T1-weighted and T2-FLAIR structural images, in a unique sample of 1,713 university students, aged 18 to 26 (mean ± SD, 21.7 ± 1.8 years). This study’s primary goal is to document the age-related variations in the regional WM properties in this cohort and describe the interrelations among the age effects on different IDPs in an effort to better understand biophysical processes underlying the late maturational changes in the WM. We additionally characterise sex and hemisphere effects in this cohort since there is some evidence that both sex and hemisphere affect WM microstructural properties and possibly the rate of their maturation (e.g. Simmonds et al., 2014; Reynolds et al., 2019; Dean et al., 2017), although specific findings are largely mixed and inconsistent (reviewed in Lebel et al., 2019; Kaczkurkin et al., 2019; Honnedevasthana Arun et al., 2021).

## 2. Methods

### 2.1. Participants

The MRi-Share study protocol was approved by the local ethics committee (CPP2015-A00850-49). All participants were recruited through the larger i-Share cohort study (for internet-based Student Health Research enterprise; www.i-share.fr). Participants signed an informed written consent form and received compensation for their contribution. Out of 2,000 individuals who were enrolled between October 2015 and June 2017, 1,823 completed the MRI acquisition protocol for both structural (T1-weighted and FLAIR) and diffusion imaging. While the study protocol allowed enrolment of students up to 35 years of age, almost 95% of our sample was under 26 years old. Table 1 provides the demographic summary of this sub-sample. Age distribution was similar in males and females, with only a marginal difference in their mean (2 months difference in age, *p* = 0.066, Welch’s *t-*test). The higher proportion of females relative to males in MRi-Share is a feature observed among university students at the French national level that is amplified in the i-Share cohort due to an over-recruitment of students coming from faculties in which an even greater proportion of women are observed.

**Table 1.** Summary of the study sample composition and relevant acquisition parameters. Age (in years) is mean (S.D.) and range, and voxel size is in mm^3^. MB: multiband factor, AP/PA: anterior-posterior/posterior-anterior; b-values are in s/mm^2^ and the number of directions is given in parentheses.

| Study sample composition |  |  |  |  |
| --- | --- | --- | --- | --- |
|  | Females | Males | Total |  |
| N | 1,246 (73%) | 467 (27%) | 1,713 |  |
| Age | 21.7 (1.7)<br>[18.1, 26.0] | 21.9 (1.8)<br>[18.1, 26.0] | 21.7 (1.8)<br>[18.1, 26.0] |  |
| MRI acquisition parameters |  |  |  |  |
| Modality | Duration<br>(min:sec) | Voxel (matrix)<br>size | Sequence key parameters |  |
| T1 | 4:54 | 1.0x1.0x1.0<br>(192x256x256) | 3D MPRAGE, sagittal, R=2, TR/TE/TI=2000/2.0/880 ms |  |
| FLAIR | 5:50 | 1.0x1.0x1.0<br>(192x256x256) | 3D SPACE, sagittal, R=2, PF=7/8,<br>TR/TE/TI=5000/394.0/1800 ms |  |
| DWI | 9:45 | 1.75x1.75x1.75<br>(118x118x84) | 2D axial, MB=3, R=1, PF=6/8, TR/TE=3540/75.0 ms, fat sat,<br>b=0 $\text{s/mm}^2$ (8PA +8AP), b=300 $\text{s/mm}^2$ (8), b=1000 $\text{s/mm}^2$<br>(32), b=2000 $\text{s/mm}^2$ (60) | |

### 2.2. MRI acquisition

The complete MRi-Share brain imaging acquisition and analysis protocols of the MRi-Share study have been detailed in Tsuchida et al. (Tsuchida et al., 2020). Briefly, all MRI data were acquired on the same Siemens 3T Prisma scanner with a 64-channels head coil (gradients: 80 mT/m - 200 T/m/s) in the two years between November 2015 and November 2017. The MRi-Share acquisition protocol closely emulated that of the UKB MR brain imaging study (Alfaro-Almagro et al., 2018), in terms of both modalities and scanning parameters, with the exception of task-related functional MRI that was not acquired in MRi-Share participants. Here, we will focus on the MRi-Share structural (T1 and T2-FLAIR) and DWI brain imaging protocol.

Table 1 summarises the key acquisition parameters for the T1, FLAIR, and DWI modalities. There were some minor differences between the MRi-Share and UKB protocols for DWI acquisition: we acquired 8, 32, and 64 directions each for *b* values 300, 1000, and 2000 s/mm^2^, respectively, while the UKB did not acquire diffusion data for a *b* value of 300 s/mm^2^ and instead acquired 50 directions each for *b* values 1000, and 2000 s/mm^2^. We also acquired more sets of *b* = 0 images acquired in Anterior-Posterior (AP) and the reverse PA phase encoding (8 pairs of AP and PA) than in UKB (3 pairs).

### 2.3. Image processing

The acquired images were managed and processed with the Automated Brain Anatomy for Cohort Imaging platform (ABACI, IDDN.FR.001.410013.000.S.P.2016.000.31235; details in Tsuchida et al, 2020). Below we briefly describe the processing steps in each pipeline pertaining to the generation of the JHU atlas ROI IDPs presented in the current paper.

#### 2.3.1. T1 and T2-FLAIR structural pipeline

Our structural pipeline processed T1 and FLAIR images for multi-channel volume- and surface-based morphometry, primarily with SPM12 (https://www.fil.ion.ucl.ac.uk/spm/) and Freesurfer v6.0 (http://surfer.nmr.mgh.harvard.edu/). For generating the regional WM volumes based on JHU atlas, we used the Jacobian-modulated WM probability map (1mm isotropic) outputted by the ‘Unified Segmentation’ framework (Ashburner and Friston, 2005) in the SPM-based volume processing branch of our pipeline (for details, see Tsuchida et al., 2020). The same Jacobian-modulated WM map was also used to obtain the total WM volume (TWMV). We also obtained the estimate of total intracranial volume (eTIV) based on the Freesurfer-branch of our pipeline.

#### 2.3.2. Field map generation pipeline

As in the UKB (Alfaro-Almagro et al., 2018), we estimated the fieldmap images from the b=0 images with opposing AP-PA phase-encoding directions from DWI scans, rather than from “traditional” fieldmaps based on dual echo-time gradient-echo images. The pipeline was built primarily using tools available from the FMRIB Software Library (FSL, v5.0.10: https://fsl.fmrib.ox.ac.uk), and provided the parameters required for eddy-current and top-up distortion corrections in the DWI pipeline. It also generated the brain mask based on the average distortion-corrected b0 maps, used for the distortion corrections in the DWI pipeline.

#### 2.3.3. Diffusion MRI pipeline

The preprocessing steps of DWI are described in the Supplementary Material section, and Supplemental Figure 1 shows the schematic representation of the pipeline. Briefly, following the eddy current and top-up distortion correction and denoising of the DWI data, the resulting image was then used to fit 1) DTI (Diffusion-Tensor Imaging; Basser et al., 1994) modelling and 2) microstructural model fitting with NODDI (Neurite Orientation Dispersion and Density Imaging; Zhang et al., 2012). For fitting DTI, volumes with the highest *b* value (*b =* 2000 s/mm^2^) were stripped from the data, as the accuracy of the fit starts to decrease above *b =* 1000 s/mm^2^ (Jensen and Helpern, 2010). Note that it still used multi-shell data, using volumes with both *b =* 300 and 1000 s/mm^2^ in addition to *b =* 0 image. For NODDI, the full set of multi-shell data was used for the fitting. The preprocessing and DTI fitting were performed using tools from FSL and the *dipy* package (0.12.0, https://dipy.org; Garyfallidis et al., 2014), while the AMICO (Accelerated Microstructure Imaging via Convex Optimization) tool (Daducci et al., 2015) was used for NODDI fitting. For each participant, the DWI processing pipeline produced 7 images in native space: fractional anisotropy (FA), mean, axial, and radial diffusivity (MD, AD, and RD), based on DTI modelling, neurite density index (NDI), orientation dispersion index (ODI), and isotropic volume fraction (IsoVF), derived from NODDI.

**Figure 1.**
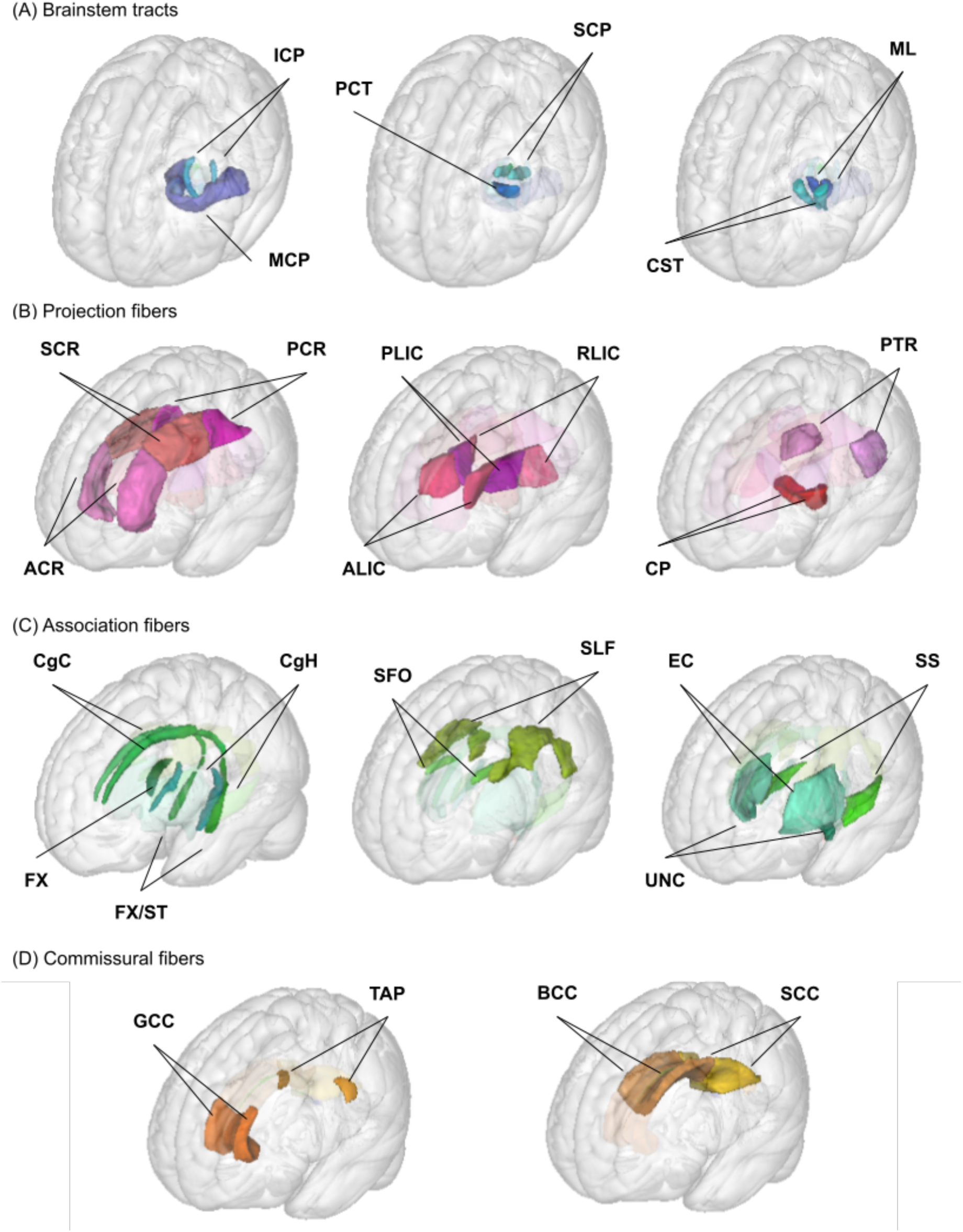
Illustration of JHU ROIs used in the analysis. Locations of the 27 ROIs (6 medially located plus 21 pairs of ROIs in each hemisphere) from the JHU ICBM-DTI-81 white matter labels atlas are shown in the glass brain for each broad group; (A) brainstem, (B) projection, (C) association, and (D) commissural fibres. See Table 1 for the full ROI name corresponding to the abbreviations in the figure.

#### 2.3.4. Generation of JHU atlas region IDPs

We used the JHU ICBM-DTI-81 white matter labels atlas (Mori et al., 2008; Oishi et al., 2008) to generate regional IDPs for each of the following metrics: regional WM volume and mean values for 4 DTI (FA, MD, AD, and RD) and 3 NODDI (NDI, ODI, and IsoVF) metrics. We used the atlas packaged with FSL v5.0.10, which does not have the orientation or labelling issues noted in other versions (Rohlfing, 2013) but is missing medial longitudinal fasciculus and inferior fronto-occipital fasciculus ROIs described by the authors of the atlas (Mori et al., 2008). We present the WM volume and mean DTI/NODDI values for 48 ROIs in this atlas. Table 2 provides the abbreviations of ROIs used in the figures and tables throughout the manuscript, and Figure 1 presents the locations of these ROIs. They are organised according to the broad classification used by the author of the atlas: 1) tracts in the brainstem, 2) projection fibres, 3) association fibres and 4) commissural fibres (Mori et al., 2008).

**Table 2.** Abbreviations of JHU atlas ROI names.

| ROI name | abbreviation | hemisphere side | ROI name | abbreviation | hemisphere side |
| --- | --- | --- | --- | --- | --- |
| Brainstem |  |  | Association |  |  |
| middle cerebellar peduncle | MCP | Both | fornix | FX | Both |
| pontine crossing tract | PCT | Both | fornix cres or stria terminalis | FX/ST | Right/Left |
| corticospinal tract | CST | Right/Left | cingulum cingulate gyrus | CgC | Right/Left |
| medial lemniscus | ML | Right/Left | cingulum hippocampus | CgH | Right/Left |
| superior cerebellar peduncle | SCP | Right/Left | superior fronto-occipital fasciculus | SFO | Right/Left |
| inferior cerebellar peduncle | ICP | Right/Left | superior longitudinal fasciculus | SLF | Right/Left |
| Projection |  |  | external capsule | EC | Right/Left |
| anterior corona radiata | ACR | Right/Left | uncinate fasciculus | UNC | Right/Left |
| superior corona radiata | SCR | Right/Left | sagittal stratum | SS | Right/Left |
| posterior corona radiata | PCR | Right/Left | Commissural |  |  |
| anterior limb of the internal capsule | ALIC | Right/Left | genu corpus callosum | GCC | Both |
| posterior limb of the internal | PLIC | Right/Left | body corpus callosum | BCC | Both |
| capsule |  |  |  |  |  |
| retrolenticular part of the internal capsule | RLIC | Right/Left | splenium corpus callosum | SCC | Both |
| posterior thalamic radiation | PTR | Right/Left | tapetum | TAP | Right/Left |
| cerebral peduncle | CP | Right/Left |  |  |  |

For extracting the DTI and NODDI metric IDPs, we first computed the rigid transform for aligning DTI and NODDI maps to the native T1 reference space (1mm isotropic) with the SPM12 ‘Coregister’ function. This transform was then aggregated with the deformation field generated in the structural pipeline to transform DTI/NODDI maps in the native DWI space to the standard template space in one step, using the SPM12 ‘Normalise’ function. When computing the mean values within each of the 48 ROIs, we used the subject-specific, spatially normalised WM probability map, thresholded at 0.5, as an inclusive mask. It ensured that the mean values were computed within regions that are primarily WM, and minimised the partial volume effects from the surrounding non-WM tissues.

### 2.4. Quality control

A detailed description of the quality control (QC) procedure for image analysis is provided in Tsuchida et al. (Tsuchida et al., 2020). Notably, visual inspection of QC images generated by the structural or DWI pipelines did not reveal any obvious problems with either the quality of acquired images or with any specific processing steps (e.g. registration of images to the reference T1 or the standard MNI space).

At the level of individual IDPs, group-level distributions were checked for any missing values as well as the extreme outliers using interquartile range (IQR), or Tukey fence method (Tukey, 1977), defining those with values below 3*IQR from the first quartile or above 3*IQR from the third quartile as the “far out” outliers. The distributions and the number of these extreme outliers for the WM volume and mean values for the 7 DTI/NODDI metrics within each of 48 JHU ROIs are reported in Supplementary Material. Four subjects did not have any volumetric or DTI/NODDI values for fornix, as the WM probability map did not overlap with this small ROI in the standard space. For the same reason, one subject was missing data for the left tapetum. In addition, for corticospinal tract (CST; *n* = 4 for both left and right) and inferior cerebellar peduncle (ICP; *n* = 5 and 6 for left and right, respectively) ROIs, mean DTI/NODDI values were not computed in the pipeline, since these ROIs extended beyond bounding box of the DWI-derived images in the standard space. Beyond these missing data, the extreme outliers were rare, and each IDP was roughly normally distributed (see Supplemental Figures 2-9). Exceptions were some ROIs, in particular those surrounded by cerebrospinal fluid and/or relatively small ROIs (e.g. fornix (FX), tapetum (TAP), brainstem ROIs), which had slightly skewed distributions, most likely caused by small misalignments in DWI-derived images and structural images in standard space. Given the large sample size, however, the effects of these outliers on the analyses were expected to be relatively minor. For simplicity, here we report the results without any outlier removal, with total sample size of 1,713 for all ROI-metric combinations, except for FX (*N* = 1,709), TAP (*N* = 1,712), CST (*N* = 1,709) and ICP (*N* = 1,707) ROIs. However, we report the comparison of results with and without outliers in Supplementary Material.

We also investigated the effects of subject motion during the DWI scan on the analyses of age, sex, and hemispheric asymmetry described below. A recent study has demonstrated that mean values of different WM metrics, including DTI and NODDI, are sensitive to in-scanner motion with varying degrees (Pines et al., 2020). However, as described in Supplementary Material, the effects of motion parameters on the estimates of age, sex, and hemispheric asymmetry were minimal in our dataset.

### 2.5. Statistical analysis

The primary goal of the present manuscript is to describe both the age and sex effects on regional WM volumes as well as microstructural properties in young adults. In addition, for 21 pairs of JHU ROIs present in each hemisphere, we also describe any asymmetry between the two hemispheres, as well as interactions between hemispheric asymmetry and age or sex. Given our sample’s narrow target age range, we expected most of the age-related variations in the volumetric and DW-IDPs to be captured by a linear age model. Indeed, a preliminary comparison of models that included quadratic age effect to capture any non-linear trend did not significantly improve model fit relative to linear age effect models in any of the IDPs examined, as judged by the Bayesian information criterion (BIC; see Supplemental Material). For both WM volumetry and microstructural property analyses, we report age and sex effects (and hemispheric asymmetry for lateralized ROIs) in a model that included eTIV to account for any differences that can be attributed to the overall head size. Even though this is not commonly done when analysing DTI/NODDI metrics, we have previously demonstrated that eTIV does impact mean DTI and/or NODDI values computed over the WM mask (Tsuchida et al., 2020) or WM skeleton (Beaudet et al., 2020). Thus, for all metrics, we tested the following models;

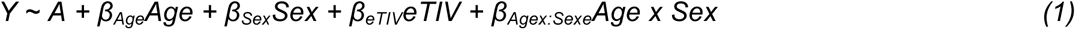

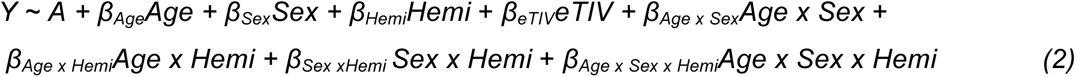

Model (1) was applied to each of the six medially located ROIs (genu, body, and splenium of the corpus callosum; fornix; middle cerebral peduncle; pontine crossing tract), while model (2) was used for the 21 pairs of left and right homotopic ROIs. Both included the sex and age interaction to test any sex differences in the age-related variations in the volumes or WM properties. Model (2) additionally tested the hemispheric asymmetry, as well as any age-related changes in asymmetry (*Age x Hemi*), sex differences in asymmetry (*Sex x Hemi*), and sex differences in age-related patterns in asymmetry (if present; *Age x Sex x Hemi*).

When analysing the volumetric asymmetry, we corrected the asymmetry due to the label size differences in the JHU ICBM-DTI-81 atlas itself by adjusting the left volume data based on the ratio of label volumes in the two hemispheres. For example, if a given ROI label had a size of 100 and 105 mm^3^ in the right and left hemispheres, respectively, the left volume for each subject was multiplied by (100/105). In effect, it computes the estimate of left volume when the two labels in each hemisphere have the same size. Therefore, any remaining asymmetry indicates the difference in the Jacobian-modulated WM maps within the ROIs in the two hemispheres.

While we mainly report the age and sex effects in a model that includes eTIV, we checked the consistency of the results with and without the correction for global volume for the regional WM volumes, and in the case of DTI/NODDI metrics, with or without global and/or regional volume corrections. The impact of global or local volume corrections is reported in Supplementary Material.

All model fits were performed in R, version 3.4.4 (R Core Team, 2018). For fitting model (1) we used the *lm* function as implemented in the *stats* library, while we used the *lmer* function from *lmer4* library (Bates et al., 2015) to fit model (2) in order to account for the repeated measures within participants (right and left hemisphere values) by including subject-specific intercepts as random effects in the linear mixed-effects model (with all other terms described above as fixed effects). The goodness of fit in the linear models (i.e. model (1)) was assessed with adjusted *R^2^*, while for the linear mixed-effects models (model (2)), we computed marginal *R^2^* using the *r.squaredGLMM* function implemented in the *MuMIn* package (Bartoń, 2020). The latter uses the method described by (Nakagawa et al., 2017) to quantify the variance explained by all the fixed effects in the linear mixed-effects models. Categorical variables in both types of models (*Sex* and *Hemi*) were deviation-coded using ‘contr.sum’ setting so that parameter estimate (*β) and t* statistics for non-categorical variables (i.e age in our case) represent those across sexes and hemispheres, and not for the specific reference sex or hemisphere (as would be in the case of treatment-coding, in the presence of interaction terms). Meanwhile, such contrast treatment results in *β* for sex and hemisphere that represents half the difference between the two sexes or the hemispheres. Continuous variables were mean-centred so that the intercept represented the value at group mean age, eTIV, etc. For all analyses, we report *p* values as significant when below 0.05 after Bonferroni correction for multiple tests (27 ROIs x 8 measures, nominal *p* threshold = 0.05/216 = 0.00023). The *p* values associated with *t* statistics in the linear mixed-effects models were obtained using Satterthwaite’s method for approximating degrees of freedom (Hrong-Tai Fai and Cornelius, 1996), as implemented in the *lmerTest* package (Kuznetsova et al., 2017). We also report generalized eta squared (*η^2^_G_*) as a measure of effect size (Olejnik and Algina, 2003), obtained using *aov_car* function in *afex* package (Singmann et al., 2021), including all terms in the model as the “observed” variables. The specification of the observed variables (as opposed to manipulated variables in other research designs) allows the correction of the effect size estimate, which makes this measure less dependent on specific research design features (Olejnik and Algina, 2003), and offers comparability across between- and within-subject designs (Bakeman, 2005; Lakens, 2013).

Visualisations of statistical summaries were created with *ggplot2 (Wickham, 2016)*, and tables were created with the *gt* package (Iannone et al., 2020) in R. Linear fitting of age effects for different sex and/or hemisphere was performed by predicting the given WM property in each sex and/or hemisphere at the mean eTIV using the *emmeans* package (Lenth, 2021). Similarly, line plots of sex and hemisphere effects were produced by predicting the mean values for each sex and hemisphere at mean age and eTIV. For evaluating the interrelations between the age, sex, or hemisphere-related variations in the regional WM volumes and DWI properties, we first computed the standardised parameter estimates (*β\**) for the respective terms in each metric using the robust standardization through refitting, implemented with the *effectsize* package (Ben-Shachar et al., 2020). Then, the *β\** values across 27 (or 21, in the case of hemisphere effects) ROIs were used to compute Pearson’s correlation between the pairwise metrics. The computation of correlation values and visualisation of the results was performed using the *Ggally* package (Schloerke et al., 2021).

## 3. Results

### 3.1. Age effects

Table 3 presents the age effects for each metric (WM volume, 4 DTI and 3 NODDI metrics) across the ROIs, and Figure 2 visually presents the summary by showing the *t* statistics as a heatmap, filtering out those that did not survive Bonferroni corrections. A similar heatmap for effect sizes (*η^2^_G_*) is presented in Supplementary Material (Supplemental Figure 10). Figure 3 provides selected scatter plots of age effects for each sex (and each hemisphere if applicable) to present the examples of such effects. Similar plots of age effects for the entire metrics and ROIs are also provided in Supplementary Material (Supplemental Figures 11-18). As evident in Table 3 and Figure 2, a number of WM ROIs showed robust age-related variations in one or more metrics we examined.

**Table 3.** Summary of age effects for each diffusion phenotype across JHU ROIs. Non-standardised parameter estimates (*β*) for age effects for each phenotype and ROI are shown (see Table 2 for the full names of abbreviated ROIs). The unit of the age effect is mm^3^/year for the volume, mm^2^/sec/year for the diffusivity measures (MD, AD and RD), and /year for FA and the NODDI phenotypes (NDI, ODI, IsoVF). Statistical significance symbols (uncorrected for multiple comparisons) *: 0.05 < *p* <0.001, **: 0.001 < *p* < 0.0001, ***: *p* < 0.0001. Bold symbols indicate Bonferroni-corrected significant p-values.

| | volume | FA ($\times 10^{-3}$) | MD ($\times 10^{-6}$) | AD ($\times 10^{-6}$) | RD ($\times 10^{-6}$) | NDI ($\times 10^{-3}$) | ODI ($\times 10^{-3}$) | IsoVF ($\times 10^{-3}$) |
| --- | --- | --- | --- | --- | --- | --- | --- | --- |
| Brainstem |  |  |  |  |  |  |  |  |
| MCP | -13.6 | 0.0 | -0.9* | -1.5* | -0.5 | 1.1* | 0.8* | -0.5* |
| PCT | -1.5 | -0.7 | -1.8 | -3.3* | -1.1 | 0.4 | 1.4* | -1.9* |
| CST | -0.7 | -1.3* | -2.6* | **-5.7\*\*\*** | -1.1 | 0.3 | 2.0* | -2.0* |
| ML | -0.7 | -1.0 | -2.5* | **-5.2\*\*\*** | -1.2 | 2.4* | 1.3 | -1.5* |
| SCP | -1.8* | -0.2 | **-2.1\*\*\*** | **-4.2\*\*\*** | -1.0* | **2.2\*\*\*** | 0.9* | **-1.3\*\*\*** |
| ICP | 0.2 | -0.3 | -2.1** | **-3.6\*\*\*** | -1.3* | **2.5\*\*\*** | 1.2* | -1.2* |
| Projection |  |  |  |  |  |  |  |  |
| ACR | 10.2* | 1.4** | **-1.9\*\*\*** | -1.3* | **-2.2\*\*\*** | **2.9\*\*\*** | -0.3 | 0.0 |
| SCR | **12.6\*\*\*** | -0.6 | **-1.2\*\*\*** | **-2.5\*\*\*** | -0.6 | **2.2\*\*\*** | **1.2\*\*\*** | 0.0 |
| PCR | 6.6** | 0.6 | -1.2** | -1.3* | -1.2* | **2.8\*\*\*** | 0.4 | 0.5* |
| ALIC | 4.0* | 0.6 | **-2.0\*\*\*** | **-2.8\*\*\*** | **-1.6\*\*\*** | **3.2\*\*\*** | 0.7* | -0.3 |
| PLIC | 2.8 | -0.7* | **-1.4\*\*\*** | **-3.3\*\*\*** | -0.5 | **2.0\*\*\*** | **1.3\*\*\*** | -0.6* |
| RLIC | 1.1 | 0.8* | **-1.6\*\*\*** | **-1.8\*\*** | **-1.5\*\*\*** | **3.5\*\*\*** | 0.5* | -0.1 |
| PTR | 2.1 | 0.3 | **-1.4\*\*\*** | **-2.1\*\*** | -1.0* | **2.1\*\*\*** | 0.5* | 0.1 |
| CP | -2.1 | -1.2* | -2.3* | **-5.9\*\*\*** | -0.5 | 1.7* | **2.4\*\*\*** | -1.6* |
| Association |  |  |  |  |  |  |  |  |
| FX | **2.8\*\*\*** | 0.7 | 0.3 | 1.1 | -0.1 | **3.7\*\*\*** | 1.5* | 2.1* |
| FX/ST | 0.4 | 1.0* | **-2.3\*\*\*** | **-3.1\*\*\*** | **-1.8\*\*\*** | **3.8\*\*\*** | 0.8* | -0.7* |
| CgC | **9.2\*\*\*** | **2.2\*\*\*** | **-2.0\*\*\*** | -0.6 | **-2.7\*\*\*** | **3.9\*\*\*** | -0.5 | -0.3 |
| CgH | **6.1\*\*\*** | 1.2* | **-3.1\*\*\*** | **-3.6\*\*\*** | **-2.8\*\*\*** | **6.9\*\*\*** | **1.6\*\*** | -0.1 |
| SFO | 0.9** | 0.1 | **-1.7\*\*\*** | **-2.7\*\*\*** | -1.3* | **3.5\*\*\*** | 0.6 | 0.0 |
| SLF | **12.4\*\*\*** | 1.0* | **-1.2\*\*\*** | -0.7 | **-1.5\*\*\*** | **2.6\*\*\*** | -0.1 | 0.1 |
| EC | 6.3* | **1.3\*\*\*** | **-1.7\*\*\*** | -1.3** | **-1.9\*\*\*** | **3.5\*\*\*** | 0.2 | 0.3 |
| UNC | -0.2 | 0.9 | **-1.8\*\*\*** | -2.2* | -1.6** | **4.1\*\*\*** | 0.8* | 0.8* |
| SS | 1.1 | **1.6\*\*\*** | **-1.9\*\*\*** | -1.1* | **-2.3\*\*\*** | **3.8\*\*\*** | -0.1 | 0.1 |
| Commissural |  |  |  |  |  |  |  |  |
| GCC | 12.5* | **1.7\*\*\*** | **-1.6\*\*\*** | -0.7 | **-2.1\*\*\*** | **2.4\*\*** | -0.5 | -0.3 |
| BCC | 32.7** | 0.6 | **-1.3\*\*\*** | -1.6* | -1.2* | **2.5\*\*\*** | 0.3 | -0.1 |
| SCC | **41.8\*\*** | **1.3\*\*\*** | **-1.5\*\*\*** | -1.1 | **-1.7\*\*\*** | **2.7\*\*\*** | 0.1 | -0.4* |
| TAP | -2.7* | 1.9* | -1.3 | 0.6 | -2.2* | 2.4* | -0.3 | 0.6 |

**Figure 2.**
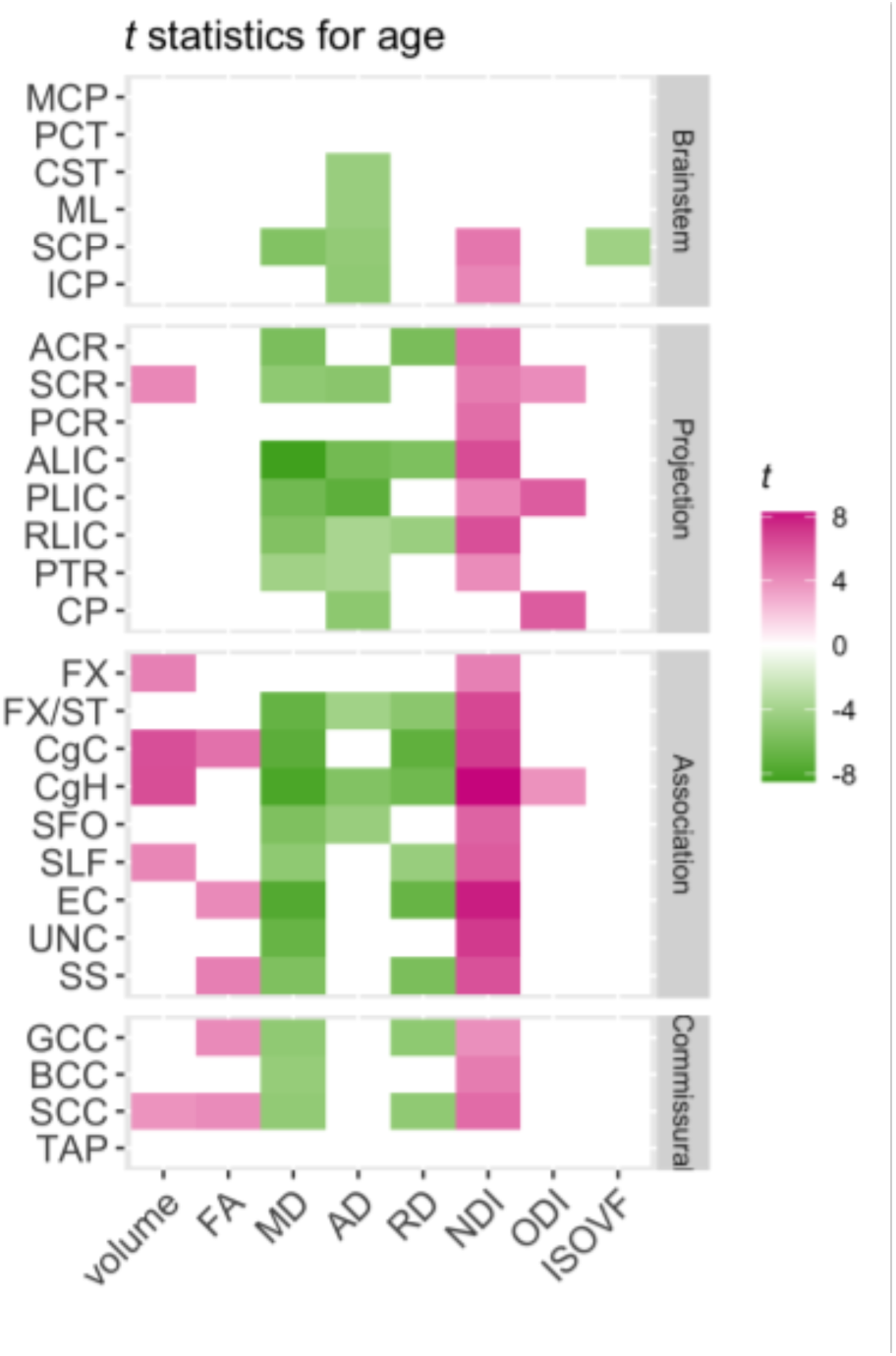
Patterns of significant age effects across WM volume and diffusion phenotypes and ROIs. Relative statistical strengths of age effects across diffusion phenotypes and ROIs are shown as heatmaps of *t* statistics (see Table 2 for the full names for the abbreviated ROIs). Those that did not survive Bonferroni corrections for multiple comparisons were filtered out (set to 0) to facilitate comparisons within significant results. Positive *t-*scores in pink indicate an age-related increase and negative values in green indicate an age-related decrease. A similar figure for effect sizes (*η^2^_G_*) is presented in the Supplementary Materials.

**Figure 3.**
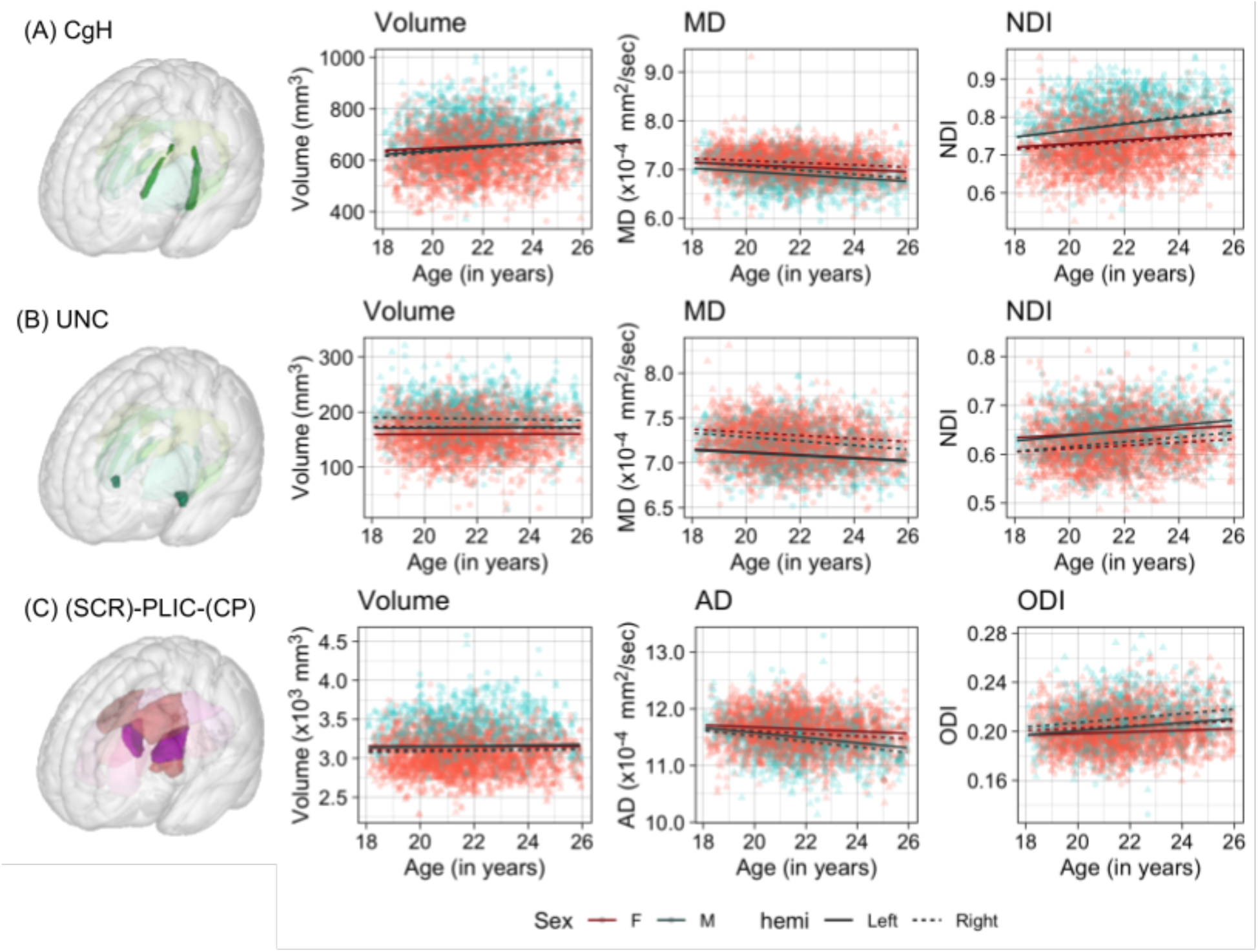
Scatter plots of individual age effects in (A) the cingulum-hippocampus (CgH), (B) uncinate fasciculus (UNC), and (C) posterior limb of the internal capsule (PLIC) ROIs. Predicted linear regression lines are superimposed for each sex (dark red: females, dark cyan: males) and hemisphere (solid: left, dashed: right) at the mean eTIV value.

The significant age-related variations in WM volumes were observed primarily in association and projection fibre ROIs, all of which increased with age. However, the apparent increase was mostly relative to eTIV, and only age-related volume increase in cingulum in the hippocampus (CgH) was consistent regardless of how, or if, the global volume was corrected (see Supplemental Material).

With or without significant volumetric increase, many of the association and projection fibre ROIs also showed robust age effects in DTI and NODDI metrics, most pronounced for MD and NDI (Figure 2). Those with significant age effects all showed an age-related increase in NDI, and decreases in diffusivity metrics. Many of these ROIs showed a tendency for the volumetric increase as well, but some showed a significant NDI increase and diffusivity decrease without any evidence for volumetric increase (see Figure 3 for examples in CgH, with the volumetric increase, and uncinate fasciculus (UNC), without). CgH additionally showed a significant age-related increase in ODI and a decrease in AD. The ODI increase and AD decrease was also observed across many ROIs in projection fibres and brainstem ROIs with varying degrees, but was particularly pronounced in the ROIs that represent a connected pathway of projection fibres: superior corona radiata (SCR), posterior limb of the internal capsule (PLIC), and cerebral peduncle (CP) (see Figure 3 for example in PLIC).

Figure 4 shows the correlation plots of the standardised parameter estimates (*β\**) for the age effect between pairs of metrics across the 27 ROIs. It shows that while the age-related variations in the regional volumes and FA values are themselves mostly non-significant, age-related increases in these metrics are related to the decrease in the MD/RD, and increase in the NDI. The age-related variations in FA are also highly and negatively related to those of ODI. Note that even though there are correlations between the age effects on the regional volume and the regional MD, RD, and NDI values, the age effects in these microstructural properties cannot be explained by the volumetric variations; the inclusion of the regional volume as a covariate in the model had a very little impact on the estimated age effects for the DTI/NODDI metrics (see Supplementary Material).

**Figure 4.**
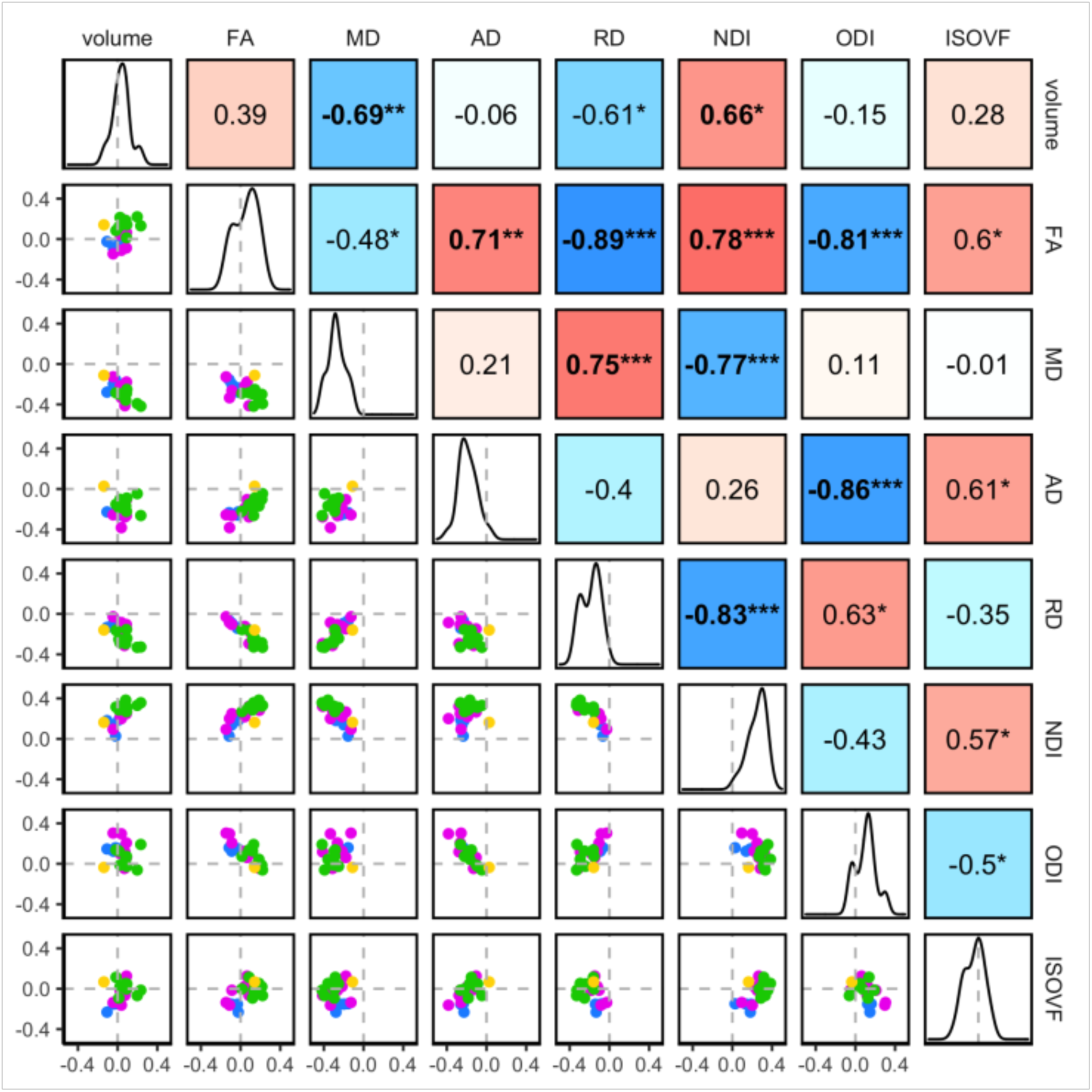
The inter-relations between the age-related variations in the regional WM properties. Pairwise correlations of the standardised parameter estimates (*β*\*) for age effects in the 27 ROIs are shown. The diagonal of the plot matrix shows the distributions of *β*\**_Age_* for the regional WM volume and DTI/NODDI values. The upper triangle shows Pearson’s correlation (*r*) values. The lower triangle shows the pairwise scatter plots of *β*\**_Age_* values in each ROI, with the colours indicating the ROI groups (*blue*: brainstem, *pink*: projection, *green*: association, *yellow*: commissural). Statistical significance symbols (uncorrected for multiple comparisons) *: 0.05 < *p* <0.001, **: 0.001 < *p* < 0.0001, ***: *p* < 0.0001. Bold-face indicates a significant correlation after Bonferroni correction for multiple comparisons.

### 3.2. Sex effects

Table 4 provides the summary of sex effects for each of the 8 metrics (volume and mean DTI/NODDI values) across the JHU ROIs, and Figure 5 presents a visual summary of statistical significance for the sex effects (see Supplemental Figure 10 for a similar figure for the effect size). Several ROI volumes were larger in males than in females even when overall head size was corrected by including eTIV in the model. Superior cerebral peduncle (SCP) showed reversed sex effects when correcting for the global volume (Table 4 and Figure 4; also see Supplementary Material for the effects of global volume correction), with larger relative volume in females than in males.

**Table 4.** Summary of sex effects for each metric across JHU ROIs. Non-standardised parameter estimates (*β*) for sex effects for each metric and ROI are shown. The units for the volume are mm^3^ and mm^2^/sec for diffusivity measures (MD, AD and RD). The *β* values represent half the difference between males and females, with positive values indicating the higher values in females than in males.

| | volume | FA ($\times 10^{-3}$) | MD ($\times 10^{-6}$) | AD ($\times 10^{-6}$) | RD ($\times 10^{-6}$) | NDI ($\times 10^{-3}$) | ODI ($\times 10^{-3}$) | IsoVF ($\times 10^{-3}$) |
| --- | --- | --- | --- | --- | --- | --- | --- | --- |
| Brainstem |  |  |  |  |  |  |  |  |
| MCP | -70.0* | **2.8\*\*** | **10.9\*\*\*** | **20.1\*\*\*** | **6.3\*\*\*** | **-12.9\*\*\*** | **-7.8\*\*\*** | **8.4\*\*\*** |
| PCT | **-14.0\*\*\*** | 3.3* | **24.6\*\*\*** | **38.2\*\*\*** | **17.9\*\*\*** | **-10.1\*\*\*** | **-17.7\*\*\*** | **25.9\*\*\*** |
| CST | **-10.8\*\*\*** | 0.3 | **23.4\*\*\*** | **35.9\*\*\*** | **17.2\*\*\*** | **-15.5\*\*\*** | **-11.9\*\*\*** | **22.6\*\*\*** |
| ML | -2.8* | 0.9 | **15.7\*\*\*** | **29.1\*\*\*** | **9.0\*\*** | **-27.3\*\*\*** | **-10.7\*\*\*** | **7.3\*\*\*** |
| SCP | **7.9\*\*\*** | **7.5\*\*\*** | **9.0\*\*\*** | **29.2\*\*\*** | -1.0 | **-16.1\*\*\*** | **-9.2\*\*\*** | **3.8\*\*\*** |
| ICP | 1.9 | 1.9 | **11.2\*\*\*** | **20.8\*\*\*** | **6.4\*\*\*** | **-20.6\*\*\*** | **-7.6\*\*\*** | 4.0** |
| Projection |  |  |  |  |  |  |  |  |
| ACR | 11.0 | 0.3 | **4.9\*\*\*** | **8.0\*\*\*** | **3.3\*\*** | **-5.9\*\*\*** | **-3.4\*\*\*** | 1.4* |
| SCR | 11.4 | 2.2* | 0.5 | 3.3* | -0.8 | **-5.7\*\*\*** | **-3.1\*\*\*** | **-2.2\*\*\*** |
| PCR | **-25.5\*\*\*** | **4.8\*\*\*** | -1.9* | 2.7* | **-4.2\*\*\*** | **-5.1\*\*\*** | **-4.4\*\*\*** | **-4.6\*\*\*** |
| ALIC | -6.4 | -2.7** | **4.2\*\*\*** | 3.8** | **4.3\*\*\*** | -1.9 | -0.8 | **3.9\*\*\*** |
| PLIC | 3.4 | -1.0 | **4.6\*\*\*** | **7.0\*\*\*** | **3.4\*\*\*** | **-9.0\*\*\*** | **-2.5\*\*\*** | **2.2\*\*\*** |
| RLIC | -7.9** | **4.3\*\*\*** | **4.5\*\*\*** | **14.0\*\*\*** | -0.3 | **-14.0\*\*\*** | **-7.0\*\*\*** | -1.5** |
| PTR | -4.8 | 1.6 | 1.7* | **5.6\*\*\*** | -0.2 | **-5.9\*\*\*** | **-2.9\*\*\*** | **-1.7\*\*\*** |
| CP | 1.4 | -1.3 | **13.9\*\*\*** | **22.4\*\*\*** | **9.6\*\*\*** | **-11.6\*\*\*** | **-3.8\*\*\*** | **9.5\*\*\*** |
| Association |  |  |  |  |  |  |  |  |
| FX | -3.7* | **4.8\*\*\*** | 3.5 | **17.0\*\*\*** | -3.2 | -4.7* | **-5.3\*\*\*** | 2.8 |
| FX/S<br>T | -0.7 | **7.2\*\*\*** | 0.8 | **11.0\*\*\*** | **-4.3\*\*\*** | **-15.5\*\*\*** | **-8.1\*\*\*** | **-4.3\*\*\*** |
| CgC | -10.4* | -0.4 | 1.6* | 2.0 | 1.4 | **-11.8\*\*\*** | **-3.1\*\*\*** | **-3.0\*\*\*** |
| CgH | 2.5 | **6.9\*\*\*** | **7.6\*\*\*** | **20.3\*\*\*** | 1.2 | **-22.4\*\*\*** | **-13.5\*\*\*** | -2.0* |
| SFO | 2.0** | 4.0* | **2.9\*\*\*** | **8.9\*\*\*** | 0.0 | -4.9** | **-5.8\*\*\*** | 0.5 |
| SLF | **-29.8\*\*\*** | -0.1 | -0.3 | -0.9 | -0.1 | **-4.3\*\*\*** | 0.0 | **-2.2\*\*\*** |
| EC | **-48.0\*\*\*** | 2.3* | **3.2\*\*\*** | **8.8\*\*\*** | 0.4 | -3.4** | **-4.6\*\*\*** | **1.8\*\*** |
| UNC | **-6.9\*\*\*** | 3.4* | 1.8* | **7.3\*\*\*** | -0.9 | -2.3 | **-3.4\*\*\*** | 0.3 |
| SS | -11.3*** | 5.6*** | 2.5* | 11.8*** | -2.1* | -11.0*** | -6.5*** | -3.2*** |
| Commissural |  |  |  |  |  |  |  |  |
| GCC | 25.1 | 0.5 | 4.5*** | 10.0*** | 1.8 | -5.4*** | -3.4*** | 2.7*** |
| BCC | 1.8 | -0.1 | 0.4 | 1.4 | -0.1 | -9.0*** | -1.5* | -2.7*** |
| SCC | -23.1 | 1.7* | 3.7*** | 10.2*** | 0.5 | -10.0*** | -3.6*** | 1.1* |
| TAP | -4.0 | -0.4 | 3.7* | 5.5* | 2.8 | -9.0*** | 2.6* | 0.5 |

**Figure 5.**
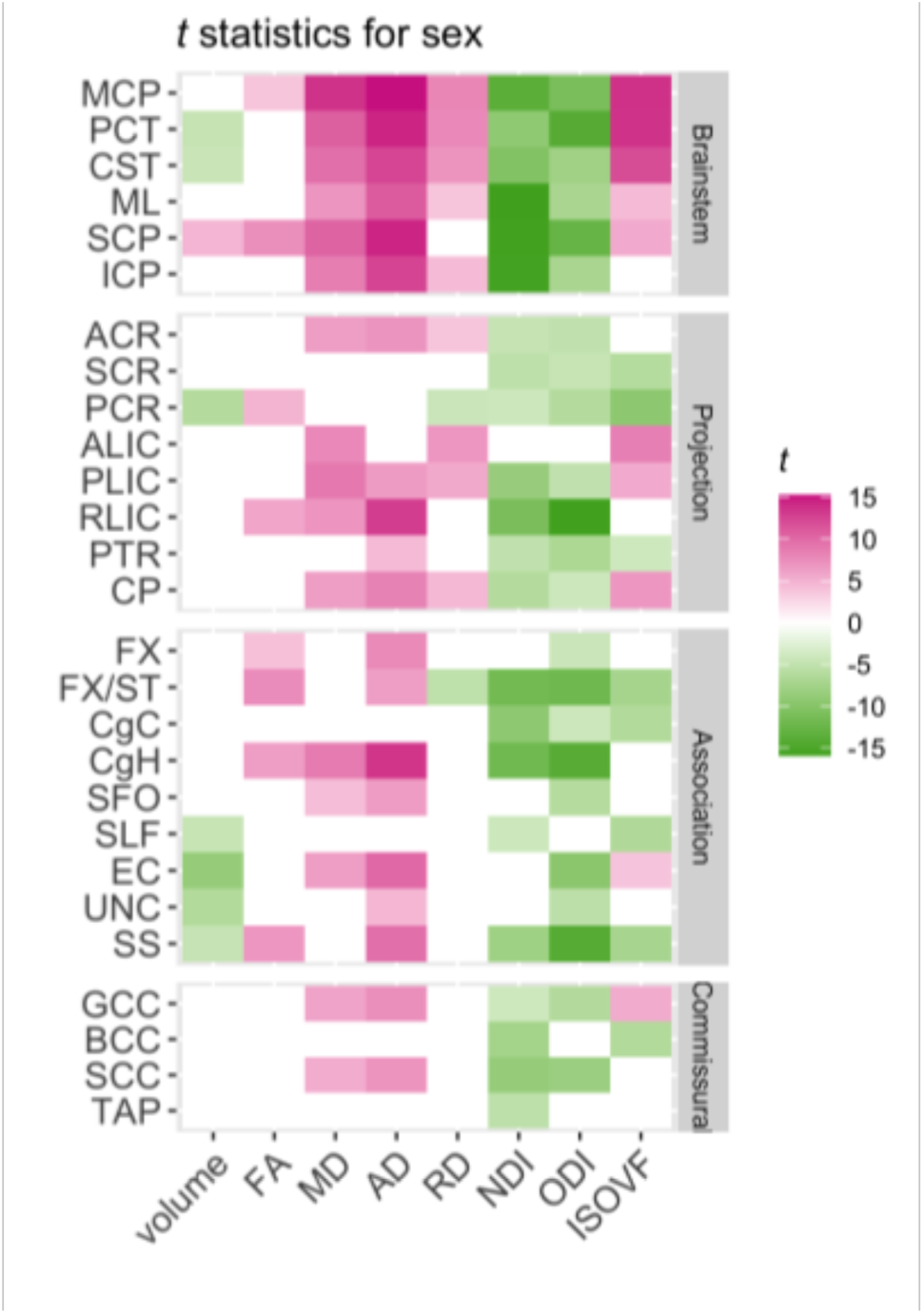
Patterns of sex effects across metrics and JHU ROIs. Relative statistical strengths of sex effects across metrics and ROIs are shown as a heatmap of *t* statistics (see Table 2 for the full names for the abbreviated ROIs). Those that did not survive Bonferroni corrections for multiple comparisons were filtered out (set to 0) to facilitate comparisons within significant results. Positive *t* scores in pink indicate higher values in females than in males, and negative values in green indicate higher values in males than in females.

For the DTI and NODDI metrics, the most substantial sex effects were present in brainstem ROIs, especially for AD and NDI. However, it was not limited to the brainstem, and AD, NDI, and in some cases ODI differences between the sexes were present across many ROIs. Overall, the sex effects were much stronger than age effects, as indicated by the magnitude of *η^2^_G_* (Supplemental Figure 10). When there were significant sex differences, diffusivity metrics were almost always higher in females than males (with the exception of RD in posterior corona radiata (PCR) and fornix/stria terminalis (FX/ST), which were higher in males), while NDI and ODI were consistently higher in males than females. Volume differences did not always accompany these sex differences in DTI and NODDI metrics (see Figure 6 for an example in middle cerebellar peduncle (MCP)), and the inclusion of ROI volumes in the model did not affect the patterns of sex differences in DTI/NODDI (see Supplementary Material).

**Figure 6.**
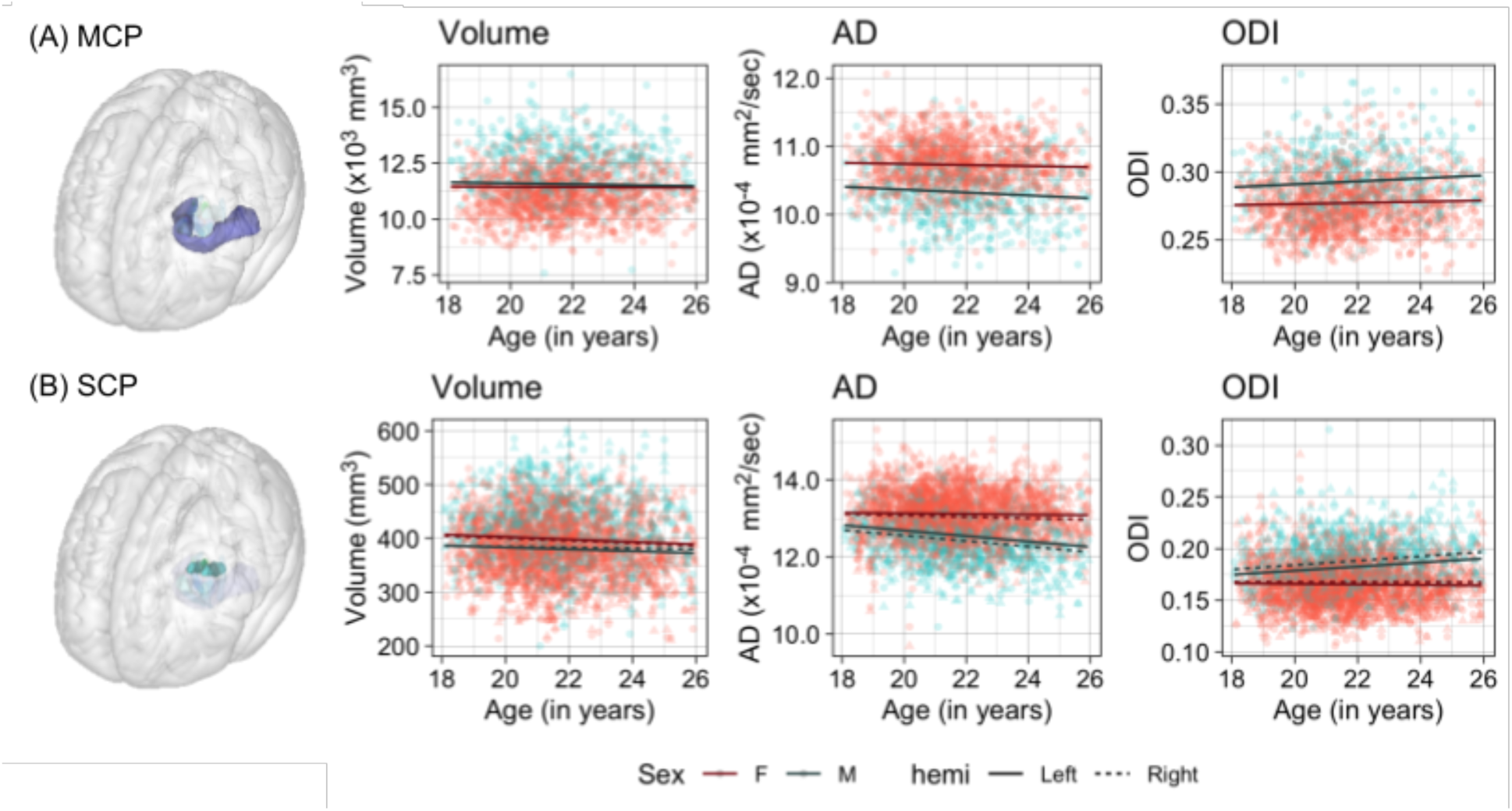
Scatter plots of individual age effects in (A) the middle cerebellar peduncle (MCP) and (B) superior cerebellar peduncle (SCP). Predicted linear regression lines are superimposed for each sex (dark red: females, dark cyan: males) and hemisphere (solid: left, dashed: right) at the mean eTIV value.

While a few ROIs showed a trend for the sex difference in the age-related changes, none of the interactions survived the Bonferonni corrections for multiple comparisons (the lowest uncorrected *p* = 0.0006). Overall, any non-significant sex differences in the age-related trajectory tend to show a steeper slope in males than in females, in particular for AD and ODI (see for example in the SCP, Figure 6 (B), and Supplemental Figures 11-18).

Figure 7 shows the pairwise correlation plots for the sex effects for the eight metrics across the 27 ROIs. Unlike the age effects, the sex difference in the regional volume or FA had a minimal relationship with the sex difference in NDI, and was primarily related to ODI difference between males and females. Also unlike the age effects, the higher NDI across ROIs in males was moderately related to the higher ODI, and both were associated with lower AD in males. The sex difference in diffusivity was associated with IsoVF differences, although the direction of sex differences in IsoVF was more mixed than diffusivity, which was greater in females across ROIs.

**Figure 7.**
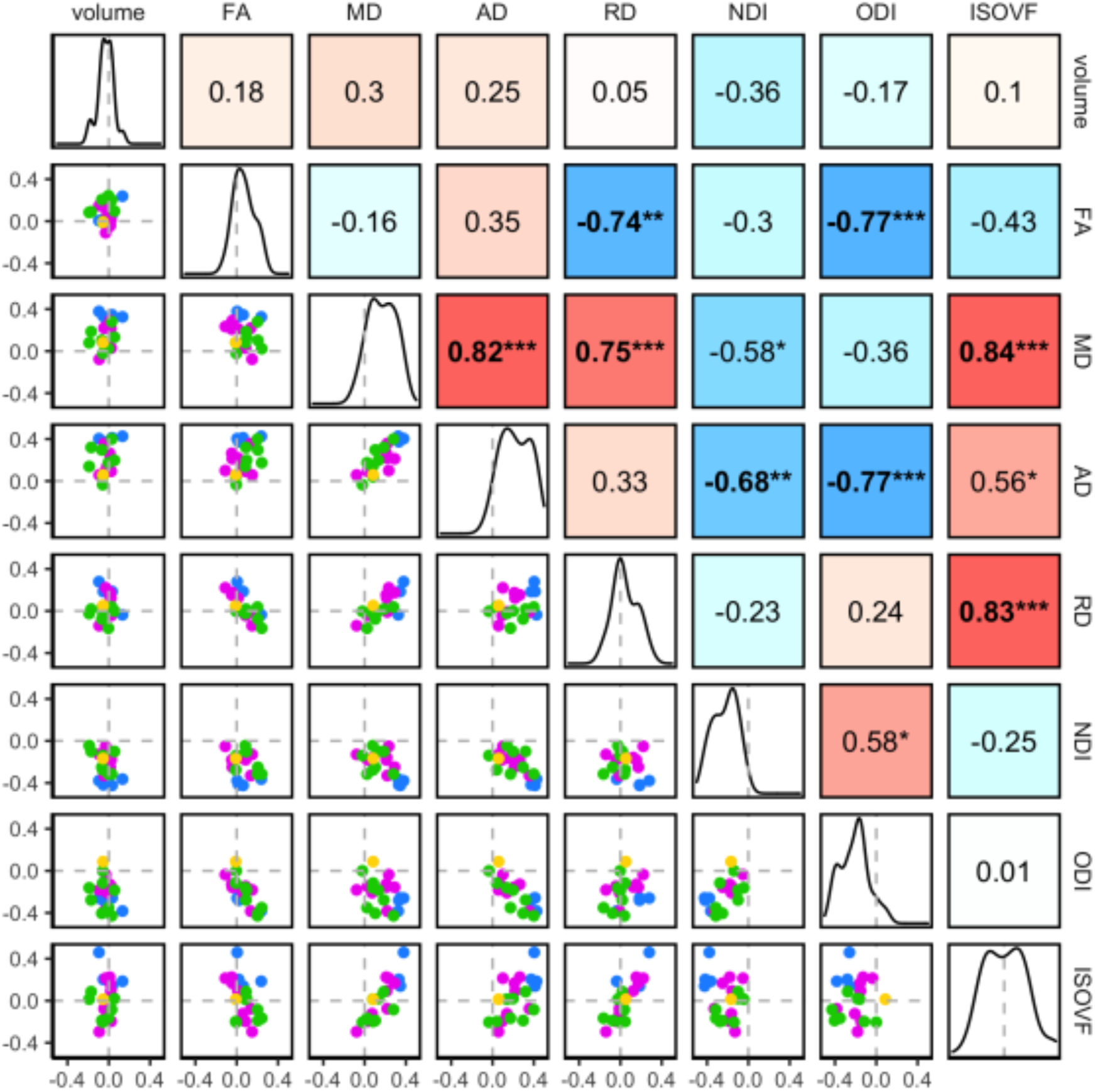
The interrelations between the sex-related variations in the regional WM properties. Pairwise correlations of the standardised parameter estimates (*β*\*) for sex effects in the 27 ROIs are shown. The diagonal of the plot matrix shows the distributions of *β*\**_Sex_* for the regional WM volume and DTI/NODDI values. The upper triangle shows Pearson’s correlation (*r*) values. The lower triangle shows the pairwise scatter plots of *β*\**_Sex_* values in each ROI, with the colours indicating the ROI groups (*blue*: brainstem, *pink*: projection, *green*: association, *yellow*: commissural). Statistical significance symbols (uncorrected for multiple comparisons) *: 0.05 < *p* <0.001, **: 0.001 < *p* < 0.0001, ***: *p* < 0.0001. Bold-face indicates a significant correlation after Bonferroni correction for multiple comparisons.

### 3.3. Hemispheric asymmetry

Table 5 summarises the hemispheric asymmetry (*β* for *Hemisphere* term) in the 21 pairs of left/right ROIs across the eight metrics. Figure 8 provides the corresponding *t* statistics for hemisphere effects that were significant after corrections (see Supplemental Figure 10 for the *η^2^_G_* heatmap).

**Table 5.** Summary of hemisphere effects for each metric across JHU ROIs. Non-standardised parameter estimates (*β*) for hemisphere effects for each metric and ROI are shown. The units for the volume are mm^3^ and mm^2^/sec for diffusivity measures (MD, AD and RD), and represent half the difference between the left and right hemisphere ROI IDPs, with positive values indicating higher values on the left than on the right side.

| | volume | FA ($\times 10^{-3}$) | MD ($\times 10^{-6}$) | AD ($\times 10^{-6}$) | RD ($\times 10^{-6}$) | NDI ($\times 10^{-3}$) | ODI ($\times 10^{-3}$) | IsoVF ($\times 10^{-3}$) |
| --- | --- | --- | --- | --- | --- | --- | --- | --- |
| Brainstem |  |  |  |  |  |  |  |  |
| CST | -0.8*** | 2.2*** | -0.8 | 3.8*** | -3.0*** | 4.2*** | -5.1*** | 0.4 |
| ML | 5.2*** | -2.5*** | 6.5*** | 8.0*** | 5.8*** | -7.4*** | -0.2 | 3.4*** |
| SCP | -0.1 | 1.2*** | 1.5*** | 5.1*** | -0.4 | -6.1*** | -2.0*** | 0.3 |
| ICP | 9.2*** | 2.9*** | 1.2* | 4.9*** | -0.6 | -11.6*** | -4.0*** | -1.9*** |
| Projection |  |  |  |  |  |  |  |  |
| ACR | 20.1*** | 0.4 | -2.0*** | -3.1*** | -1.4*** | 1.1*** | 2.4*** | -1.9*** |
| SCR | 21.6*** | 5.4*** | 0.2* | 4.2*** | -1.7*** | 1.6*** | -2.1*** | 0.0 |
| PCR | -1.1 | -8.2*** | -4.8*** | -15.1*** | 0.4 | 9.5*** | 8.6*** | 1.4*** |
| ALIC | **-30.9\*\*\*** | **-9.7\*\*\*** | **-5.1\*\*\*** | **-17.7\*\*\*** | **1.2\*\*\*** | **-5.5\*\*\*** | **11.4\*\*\*** | **-8.3\*\*\*** |
| PLIC | **26.5\*\*\*** | **9.0\*\*\*** | **-3.0\*\*\*** | **3.5\*\*\*** | **-6.3\*\*\*** | **1.2\*\*\*** | **-3.0\*\*\*** | **-3.8\*\*\*** |
| RLIC | **-12.2\*\*\*** | **18.5\*\*\*** | **-5.4\*\*\*** | **16.2\*\*\*** | **-16.2\*\*\*** | **-7.0\*\*\*** | **-8.8\*\*\*** | **-12.0\*\*\*** |
| PTR | **-30.9\*\*\*** | **7.4\*\*\*** | **-5.6\*\*\*** | -0.8* | **-8.1\*\*\*** | **5.1\*\*\*** | **-1.2\*\*\*** | **-3.9\*\*\*** |
| CP | **13.6\*\*\*** | **1.9\*\*\*** | **2.6\*\*\*** | **6.0\*\*\*** | 0.9 | **-4.4\*\*\*** | 0.9* | **1.6\*\*\*** |
| Association |  |  |  |  |  |  |  |  |
| FX/ST | **90.0\*\*\*** | **-1.7\*\*\*** | **-15.5\*\*\*** | **-29.7\*\*\*** | **-8.4\*\*\*** | **-8.7\*\*\*** | **5.0\*\*\*** | **-17.1\*\*\*** |
| CgC | 2.9 | **18.0\*\*\*** | 0.3* | **25.8\*\*\*** | **-12.5\*\*\*** | **1.7\*\*\*** | **-13.1\*\*\*** | **-2.3\*\*\*** |
| CgH | 2.4** | **4.2\*\*\*** | **-4.7\*\*\*** | **-2.7\*\*\*** | **-5.7\*\*\*** | 0.8 | **-3.0\*\*\*** | **-4.6\*\*\*** |
| SFO | **3.4\*\*\*** | **-2.2\*\*\*** | -0.6* | **-3.7\*\*\*** | 0.9* | **4.8\*\*\*** | **5.4\*\*\*** | -0.6 |
| SLF | **14.0\*\*\*** | -0.4* | **-4.5\*\*\*** | **-8.3\*\*\*** | **-2.7\*\*\*** | **6.1\*\*\*** | **4.0\*\*\*** | **-1.4\*\*\*** |
| EC | **105.8\*\*\*** | **10.5\*\*\*** | **-6.8\*\*\*** | **0.9\*\*\*** | **-10.6\*\*\*** | **-5.0\*\*\*** | **-5.5\*\*\*** | **-12.0\*\*\*** |
| UNC | **-7.2\*\*\*** | **4.5\*\*\*** | **-9.5\*\*\*** | **-11.0\*\*\*** | **-8.8\*\*\*** | **12.9\*\*\*** | 0.7* | **-2.0\*\*\*** |
| SS | **-30.9\*\*\*** | **6.8\*\*\*** | **-6.2\*\*\*** | -0.2 | **-9.2\*\*\*** | 0.8 | **-1.0\*\*\*** | **-8.4\*\*\*** |
| Commissural |  |  |  |  |  |  |  |  |
| TAP | **-23.4\*\*\*** | **-2.6\*\*\*** | **3.7\*\*\*** | 1.6 | **4.8\*\*\*** | **7.0\*\*\*** | **6.8\*\*\*** | **10.2\*\*\*** |

**Figure 8.**
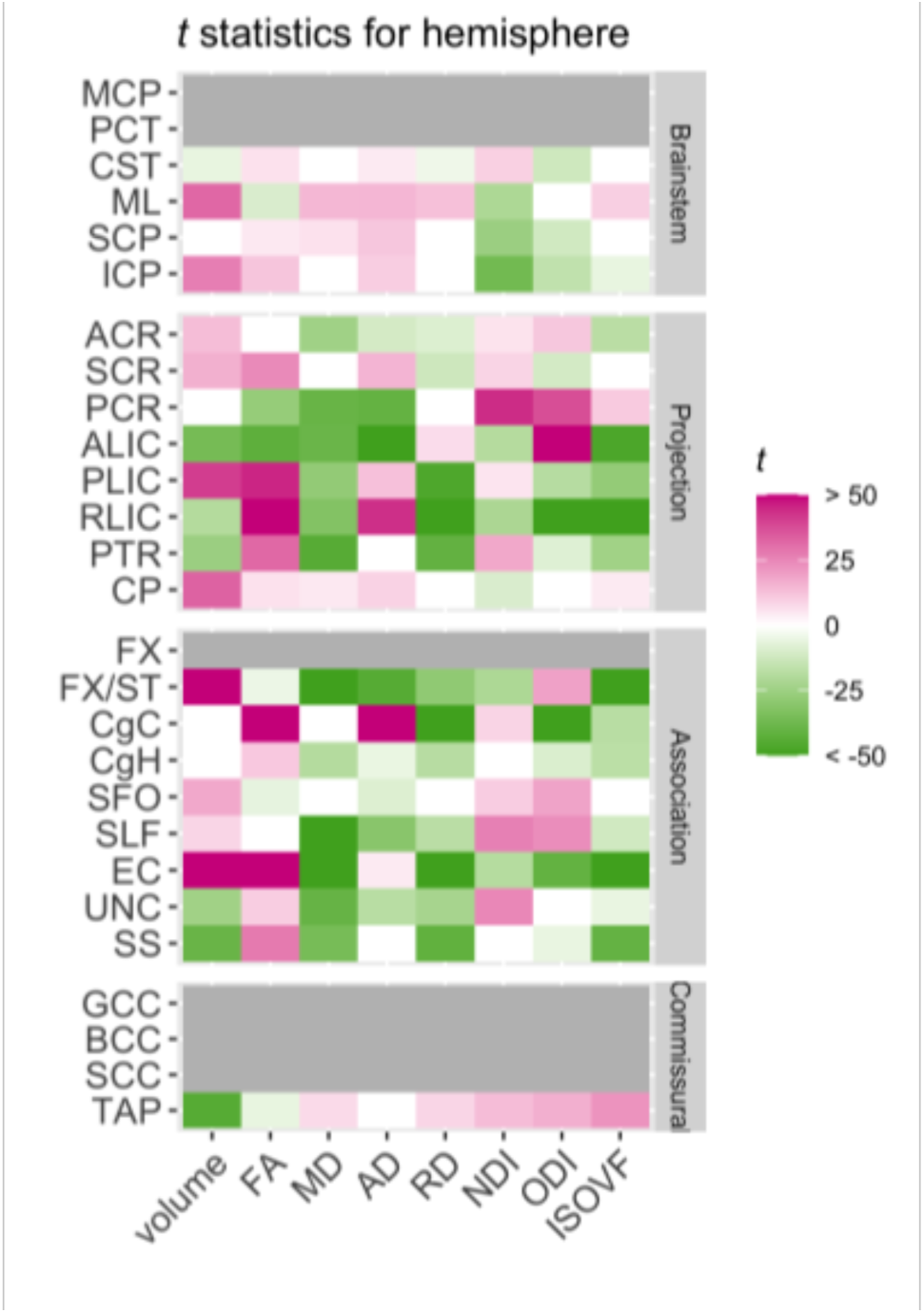
Patterns of hemispheric asymmetry across metrics and ROIs. Relative statistical strengths of hemisphere asymmetry across metrics and ROIs are shown as heatmaps of *t* statistics (see Table 2 for the full names for the abbreviated ROIs). Those that did not survive Bonferroni corrections for multiple comparisons were filtered out (set to 0) to facilitate comparisons within significant results. Positive *t* scores in pink indicate higher values in the left than in the right hemisphere, and negative values in green indicate higher values in the right than in the left.

There was a widespread hemispheric asymmetry across many ROIs, with the effect sizes that dwarfed those of age or sex effects (see Supplemental Figures 19 - 26 for plots comparing values in each hemisphere for each metric and ROI). For the regional WM volumes, the asymmetry was significant even though we corrected for label size asymmetry in the atlas itself. The direction of asymmetry was mixed, with some ROIs showing larger volume in the left than the right side (e.g. fornix and stria terminalis (FX/ST) and external capsule (EC)) while others larger in the right than left (e.g. sagittal stratum (SS), uncinate fasciculus (UNC), and tapetum (TAP). There was mixed volumetric asymmetry even within the connected ROIs of the projection fibres, from corona radiata (anterior (ACR), superior (SCR), and posterior (PCR)), internal capsule (anterior (ALIC) and posterior limb (PLIC), and retrolenticular part (RLIC)), then to cerebral peduncle (CP).

A number of ROIs also showed asymmetry in DTI/NODDI metrics. The hemispheric asymmetries in mean DTI/NODDI had very little relationship with the volumetric asymmetries (Figure 9), and were not affected by the inclusion of left/right ROI volumes in the model (see Supplementary Material). The asymmetries in DTI metrics were primarily associated with the asymmetry in ODI, such that the hemisphere showing higher ODI typically had lower FA and AD, and higher RD than the other hemisphere (Figure 9). In contrast, the asymmetry in NDI was more or less independent of diffusion metric asymmetry. Those with a strong leftward asymmetry for ODI (i.e. higher ODI values on the left compared to the right side) included the anterior part of the internal capsule (ALIC), posterior corona radiata (PCR), superior fronto-occipital fasciculus (SFO), superior longitudinal fasciculus (SLF), and fornix and stria terminalis (FX/ST) ROIs. Among them, the PCR, SFO and SLF ROIs also showed a leftward asymmetry for NDI, while ALIC and FX/ST had higher NDI on the right side. In contrast, the posterior limb and retrolenticular part of the internal capsule (PLIC and RLIC), cingulum in the cingulate gyrus (CgC) and external capsule (EC) ROIs were among those showing a rightward asymmetry for ODI. The RLIC and EC ROIs showed a rightward asymmetry for NDI as well. The uncinate fasciculus (UNC) ROI showed a leftward asymmetry for NDI without any significant asymmetry in ODI.

**Figure 9.**
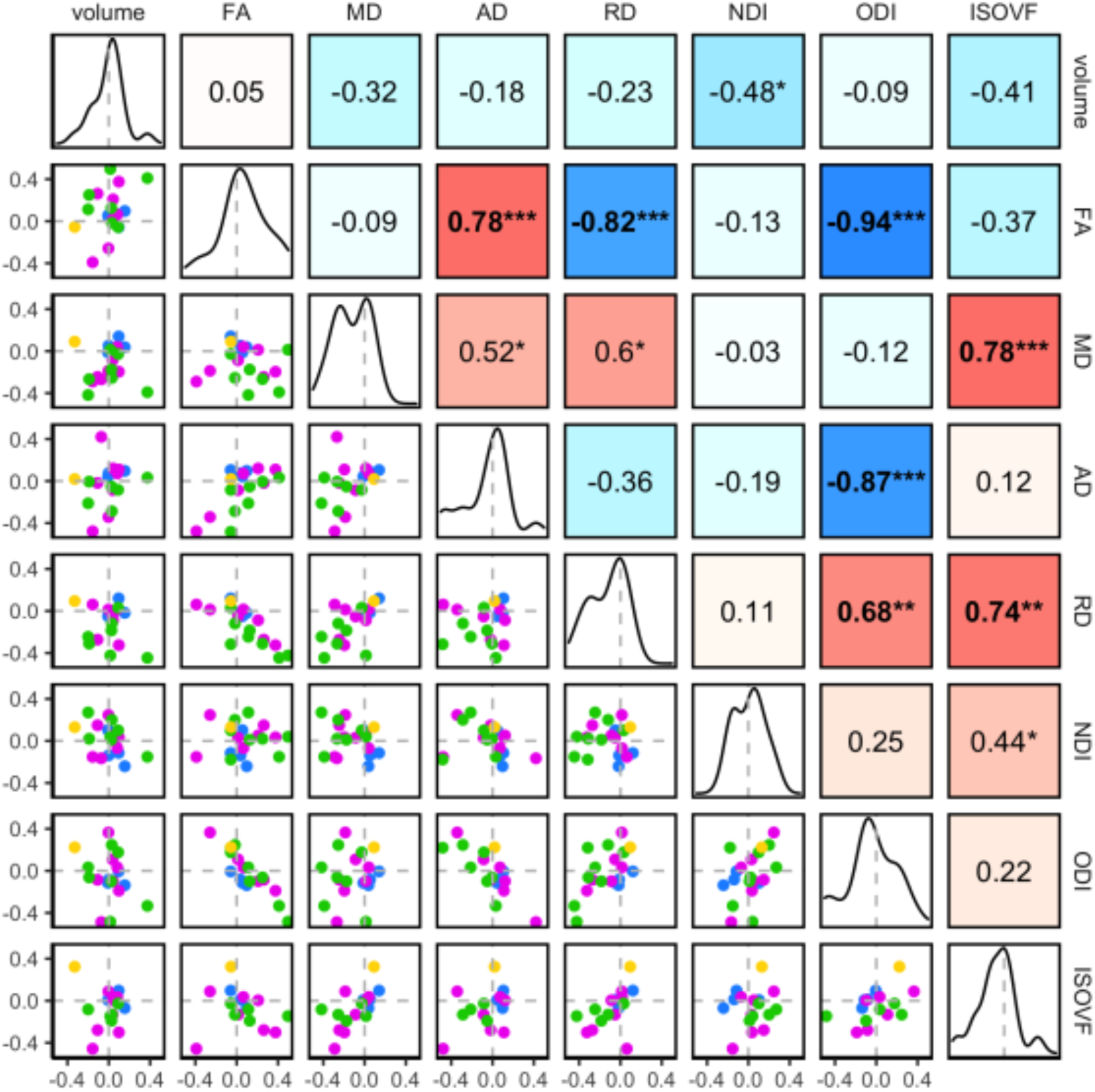
The interrelations between the hemisphere-related variations in the regional WM properties. Pairwise correlations of the standardised parameter estimates (*β*\*) for hemisphere effects in the 21 ROIs are shown. The diagonal of the plot matrix shows the distributions of *β*\**_Hemi_* for the regional WM volume and DTI/NODDI values. The upper triangle shows Pearson’s correlation (*r*) values. The lower triangle shows the pairwise scatter plots of *β*\**_Hemi_* values in each ROI, with the colours indicating the ROI groups (*blue*: brainstem, *pink*: projection, *green*: association, *yellow*: commissural). Statistical significance symbols (uncorrected for multiple comparisons) *: 0.05 < *p* <0.001, **: 0.001 < *p* < 0.0001, ***: *p* < 0.0001. Bold-face indicates a significant correlation after Bonferroni correction for multiple comparisons.

**Figure 10.**
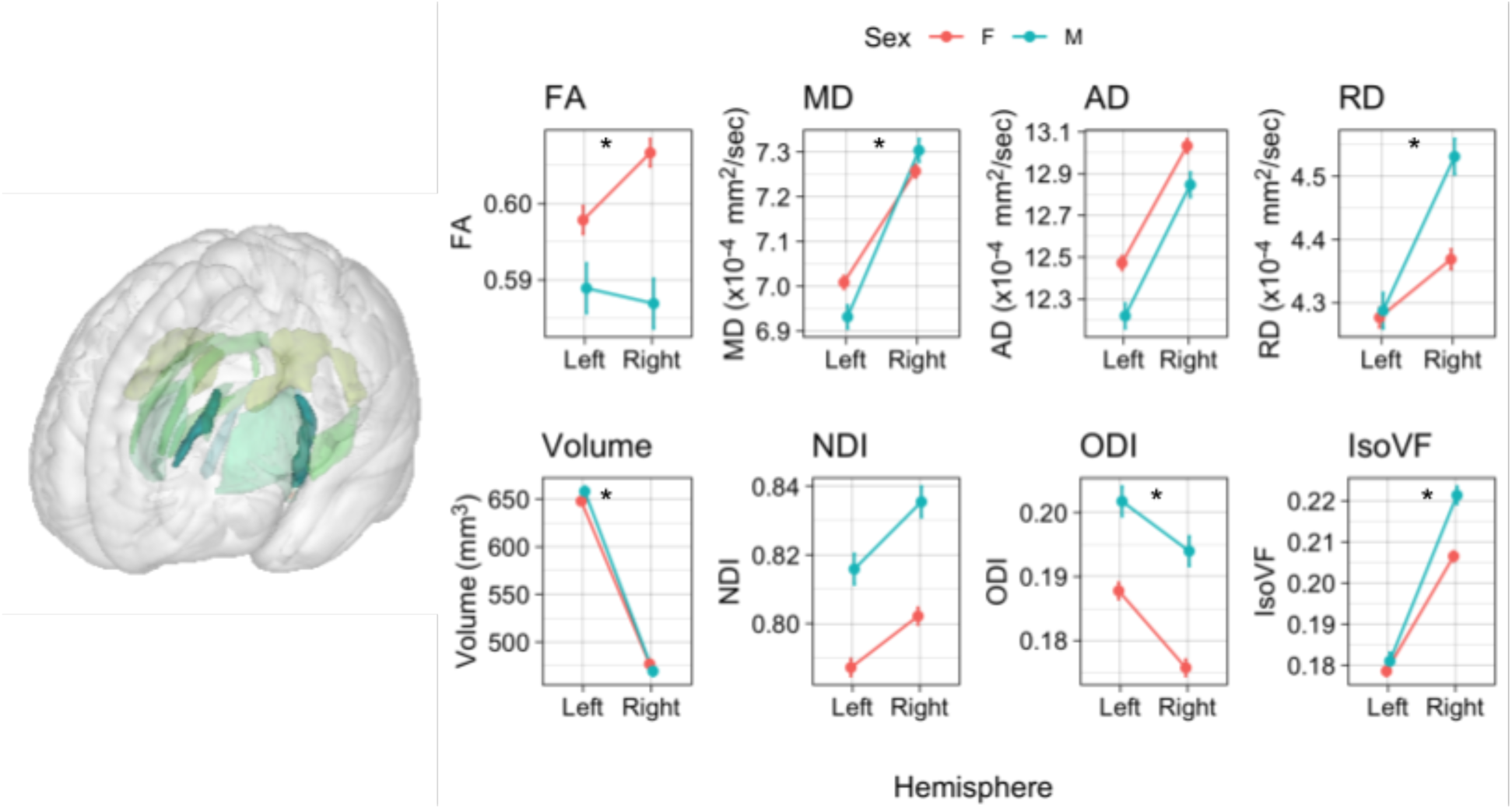
Hemispheric asymmetry in fornix and stria terminalis (FX/ST) ROI WM properties. Predicted values of each metric in the FX/ST are shown for each sex and hemisphere, at mean age and eTIV. The error bars indicate 95% confidence intervals for the prediction. * denotes significant sex and hemisphere interactions. F: female, M: male.

Hemispheric asymmetry was significantly different between males and females in several ROIs (Supplemental Tables 2-9), most notably in the FX/ST, which showed significant sex and hemisphere interactions for all metrics except for AD and NDI. However, the effect sizes for the interactions were very small, accounting for less than 1 % of the total variance in all cases (highest in FA in the FX/ST, with *η^2^_G_* = 0.007). In the FX/ST ROI, males showed greater hemispheric asymmetry than females for volume, MD, RDm and IsoVF (leftward for volume, rightward for MD, RD and IsoVF). For FA, males and females showed reversed asymmetry patterns, with females showing rightward asymmetry while males showing slightly leftward asymmetry.

A few ROIs also showed significant interactions between age and hemisphere, indicating age-related variations in the strengths of hemispheric asymmetry; the IsoVF asymmetry in the FX/ST and PTR was smaller in older subjects, while it was larger in the PCR (Supplemental Table 9 and Supplemental Figure 18). The FX/ST additionally showed a slightly smaller MD asymmetry in older subjects (Supplemental Table 4 and Supplemental Figure 13). However, the effect sizes for these interactions were minimal, with the highest *η^2^_G_* of 0.002 for IsoVF in PTR. There were no significant three-way interactions between age, sex, and hemisphere after multiple comparison corrections (the lowest uncorrected *p* = 0.0059).

### 3.4. Summary of the results

In a sample of students aged between 18 and 26 years, we observed significant age-associated increases in NDI and decreases in all diffusivity metrics (most pronounced for MD) across many JHU ROIs. Many ROIs in projection and brainstem fibre groups showed primarily significant decreases in AD, and association and commissural fibre groups were more characterised by decreases in RD. There were strong main effects of sex across many ROIs without interactions with the age effects. Globally, males had higher NDI and ODI than females, while diffusivity was higher in females than males. We also observed robust hemispheric asymmetries across many ROIs. Several ROIs showed sex differences in the degree of, and a few cases in the direction of, hemispheric asymmetry. For the most part, there were very few age-related variations in the degree of hemispheric asymmetry.

The patterns of interrelations among the estimated effects on different WM properties were distinct for each effect. The age-related variations in volume and FA were both correlated with the age-related increase in NDI, but FA was also affected by the age-related variation in ODI. Although some regions showed a significant age-related increase in ODI, this increase had no association with the degree of age-related increase in NDI. In contrast, the sex effects on NDI and ODI were moderately correlated, indicating that the greater NDI in males is associated with the greater ODI. The degree of hemispheric asymmetry in NDI was not related to volumetric asymmetry or asymmetries in diffusion properties. The left-right asymmetries in diffusion properties were highly correlated with ODI asymmetry, indicating that the widespread asymmetries in diffusion properties primarily reflect differences in the degree of fibre orientation complexity/coherence in the corresponding ROIs in each hemisphere.

## 4. Discussion

The primary objective of the present study was to characterise the late maturational changes in the regional WM properties during post-adolescence in the large and unique sample from the MRi-Share database. We also examined sex differences and hemispheric asymmetry in the WM of this sample and assessed whether the age-related changes differed between two sexes or across hemispheres. Below we discuss our main findings in relation to the existing literature, comment on the specific features of our dataset, and methodological strengths and limitations of the present study.

### 4.1. Age-related variations in regional WM properties

We observed wide-spread age-related increases in the NDI, a measure of neurite density, as well as decreases in diffusivity (MD, AD, and RD) across many of the JHU ROIs in our sample of young adults aged between 18 and 26 years. Although more regionally variable and statistically weaker, FA and volumes tended to show an age-related increase overall. The age-related increases in both the regional volume and FA values were correlated with the changes in the NDI and diffusivity. We also observed more regionally specific age-related increases in ODI, mostly in projection fibre ROIs. These global patterns are consistent with a wealth of literature showing a relatively protracted maturation of human brain WM: both developmental and life-span studies of WM volume and DTI metrics have indicated continued increases in global WM volume and FA into young adulthood, together with decreases in diffusivity that only peaks sometime in young to mid-adulthood (Hasan et al., 2007, 2010; Westlye et al., 2010; Lebel et al., 2012; Slater et al., 2019; Beaudet et al., 2020; Tsuchida et al., 2020). More recent studies using NODDI have also shown the continuous increase of NDI through development (Mah et al., 2017; Genc et al., 2017; Dimond et al., 2020; Lynch et al., 2020; Pines et al., 2020) and adulthood (Chang et al., 2015; Billiet et al., 2015; but see Kodiweera et al., 2016), peaking around the fourth and fifth decade of life (Slater et al., 2019; Qian et al., 2020). ODI, on the other hand, has not been reported to change noticeably during development (Dimond et al., 2020) or show a slight decrease in some tracts (Lynch et al., 2020), but starts to increase during young adulthood (Chang et al., 2015; Slater et al., 2019) that continues through ageing (Beck et al., 2021; Billiet et al., 2015).

More robust and wide-spread increase in NDI than FA observed in our data likely results from the fact that we sampled the FA values from the entire WM regions within each ROI, rather than a limited ‘core’ region with high FA values, a common approach in studies using the same JHU atlas, as discussed in the section on Methodological considerations below. When sampling over regions with more complex fibre organizations, it is likely that NODDI better approximates axonal packing than a simple tensor model, in which FA can be influenced by both the fibre density and myelination as well as by the composition of fibre orientations (among other things) in the sampled voxel (Zhang et al., 2012; Jones et al., 2013). This interpretation is corroborated by the negative correlations observed between the age effects on the regional FA and ODI; it suggests that even though the age-related increase in neurite density drives an FA increase, concomitant increases in fibre orientation complexity in some regions attenuates such increase.

Regionally, we observed that cingulum WM showed a prominent age-related increase in NDI as well as MD and RD decreases. The NDI increase, with concurrent RD reduction, is suggestive of increased axonal packing and/or myelination (Song et al., 2005; Jelescu and Budde, 2017). Cingulum WM in both cingulate gyrus (CgC) and hippocampal region (CgH) also showed a robust volumetric increase as well, both in terms of raw volume and relative to eTIV or TWMV. However, in these and other ROIs, the regional volume had little impact on the observed age effects on other WM properties, suggesting the distinct biological processes governing the age-related changes in WM volumes and other metrics related to microstructural properties (Lebel et al., 2019). Previous studies have indicated cingulum to be one of the last major tracts to mature during development, reaching peak values in FA or minimum values in MD later than other tracts (Westlye et al., 2010; Lebel et al., 2012; Tamnes et al., 2010). Similarly, a higher rate of NDI growth in limbic tracts that include CgC and CgH has been reported in a sample of 66 healthy subjects with a mean age of 25 years (Chang et al., 2015). A more recent and larger-scale lifespan study on regional DTI and NODDI metrics in 801 individuals aged 7 to 84 years has also indicated a relatively late peak age for NDI in CgC and CgH (Slater et al., 2019). The cingulum bundle primarily contains fibres that link cingulate gyrus and hippocampus (Mori et al., 2008), but also consists of many short association fibres that interconnect medial parts of the frontal, parietal, and temporal regions (Heilbronner and Haber, 2014). With the diverse fibre populations that make up this bundle, neuroimaging studies in healthy subjects as well as in clinical populations have implicated this region for a wide range of cognitive functions: these include executive control, motivation, and pain in anterior/dorsal cingulate and memory in hippocampal region (reviewed in Bubb et al., 2018). Several studies have also shown the link between the microstructural integrity of the cingulum bundle and cognitive performance in children (Bathelt et al., 2019) and older adults (Kantarci et al., 2011; Bettcher et al., 2016). In this context, robust age-related changes observed in the cingulum ROIs in our sample of young adults undergoing higher-level education are particularly interesting. Future studies should investigate the relevance of volumetric and microstructural differences across subjects in cingulum to cognitive and academic performance and emotional and behavioural development in these subjects.

Beyond the cingulum bundle, all association ROIs tended to show a higher increase in NDI (average annual percentage increase, computed from the base value at age 18) of 0.56 %/yr, ranging from 0.37 %/yr in SLF to 0.94 %/yr in CgH) than commissural (average of 0.35 %/yr, ranging from 0.33 %/yr in GCC and 0.41 %/yr in TAP) and projection ROIs (average of 0.35 %/yr, ranging from 0.19 %/yr in CP and 0.46 %/yr in RLIC). The NDI increase was smallest in the brainstem ROIs (average of 0.17 %/yr) and least statistically significant. Of note, the brainstem ROIs also had the highest estimated NDI at age 18 (mean [range] = 0.89 [0.83 - 0.96]), while association fibre ROIs had the lowest estimated NDI at the same age (mean [range] = 0.71 [0.62 - 0.80]). It suggests a possibility that most of the NDI growth in brainstem ROIs takes place earlier than the age-range of our sample. The observed pattern is broadly consistent with previous DTI studies suggesting earlier maturation in the commissural and projection fibres, followed by association fibres, especially in fronto-temporal regions (Westlye et al., 2010; Tamnes et al., 2010; Lebel et al., 2012). More recent studies with NODDI also support similar regional patterns of the developmental trajectory (Dean et al., 2017; Slater et al., 2019; Lynch et al., 2020). For instance, in a recent study examining the maturational timing of regional NODDI parameters in a cross-sectional sample of 104 subjects aged between 0 to 18 years, the NDI growth in callosal fibres reached a plateau the earliest, followed by projection and association fibres (Lynch et al., 2020).

While relatively modest in terms of NDI growth, we found that the connected ROIs of projection fibres, from superior corona radiata (SCR), through the posterior limb of the internal capsule (PLIC), then to cerebral peduncle (CP), showed the age-related increase in ODI and decrease in AD. It suggests the increasing fibre complexity in this large WM bundle that contains the pyramidal and cortico-pontine tracts. This observation is novel, and has not been reported in previous studies examining age-related variations in regional NODDI values in subjects with age-range that overlaps with our study (Chang et al., 2015; Slater et al., 2019; Pines et al., 2020; Billiet et al., 2015): None of these studies reported notable age-related ODI increase in this projection fibre pathway that stood out from other regions (e.g. non-brainstem projection fibre ROI in Chang et al., 2015 and tractography-based corticospinal tract in Slater et.al., 2019). However, different methodology in defining the tract ROI as well as modelling strategies makes the direct comparison difficult: Future studies are needed to confirm the validity of our observation and investigate the functional relevance of such age-related changes.

### 4.2. Sex differences

In many ROIs, we detected significant sex differences in the regional WM properties, but found very little evidence for sex differences in the age-related variations in the WM properties. For the main effects of sex on the regional WM volumes, larger volumes in males than in females in many ROIs became non-significant with the inclusion of eTIV in the model, with a reversed sex effects in a few ROIs. For the DTI and NODDI metrics, we found higher diffusivity, in particular AD, in females than males, and higher NDI and ODI in males than in females across many JHU ROIs. These overall trends are consistent with our previous study that found higher diffusivity in females than males and higher NDI and ODI in males than females in the cerebral WM of the same MRiShare sample (Tsuchida et al., 2020). The present study additionally revealed regional variations in the overall pattern, with the most substantial differences observed in the brainstem ROIs. The observed differences in DTI and NODDI metrics were not related to the sex differences in the global or regional volumes. Examination of the inter-correlations among the sex effects in each metric revealed moderate correlations between NDI, ODI, and AD sex differences, suggesting that both higher axonal density and lower fibre coherence (or higher fibre complexity) in males in many ROIs are driving the overall sex differences.

The overall lack of sex differences in the age-related variations of regional WM properties in our study is in accordance with prior studies that report no or minimal interaction between sex and age in DTI (Hsu et al., 2010; Hasan et al., 2010; Tamnes et al., 2010; Inano et al., 2011; Kochunov et al., 2011; Lebel et al., 2012; Pohl et al., 2016) or NODDI (Kodiweera et al., 2016; Cox et al., 2016; Slater et al., 2019; Lynch et al., 2020), except earlier in development, when steeper age-related changes in boys than girls have been reported (Simmonds et al., 2014; Reynolds et al., 2019). Similarly, while not surviving multiple comparison corrections, any trend for the sex differences in age-related variations in our data (e.g. AD and ODI in the superior cerebellar peduncle (SCP)) indicated steeper age-related changes in males than in females.

For the main effects of sex, findings from the prior work on DTI in youth have been mixed, with many studies reporting no significant difference (e.g. Lebel et al., 2008; Tamnes et al., 2010). However, those reporting positive findings often show higher FA and lower MD in males (e.g. Lebel and Beaulieu, 2011; Herting et al., 2012). While generally lower diffusivity in males in our study is consistent with such findings, we found higher FA values in females in many ROIs. A partial explanation for the discrepancy comes from one large-scale, multi-site study of WM microstructure in youth, which showed that the higher FA values in males were partly reversed when controlling for the supratentorial volume difference (Pohl et al., 2016). Although we still observed higher FA in females in many ROIs, we did find more ROIs where FA values are higher in males when not controlling for the eTIV difference by including it as a covariate in the model. In contrast, the statistical inference for diffusivity differences (lower in males overall) was generally not affected by the inclusion of eTIV or TWMV in the model. In our recent multi-cohort study, we investigated mean DTI metrics in the WM skeleton in a large sample (total N >20,000) that included the MRi-Share and 9 other cohorts that collectively spanned the entire adult lifespan, and found significantly greater AD and MD in females than males in the pooled dataset (this study also included eTIV as a covariate in the model). In the same study, the sex effects on FA varied across the cohorts, and there was no significant sex difference in the pooled dataset (Beaudet et al., 2020).

With regard to NODDI, most studies do not report any sex differences in childhood and adolescence (Genc et al., 2017; Mah et al., 2017; Dimond et al., 2020; Lynch et al., 2020). However, one study with an adult sample (age range of 18 to 55 years) reported a robustly higher NDI and ODI in males than in females (Kodiweera et al., 2016), similar to our findings. The higher values for both NDI and ODI in males than females across many ROIs in our study indicate both higher axonal packing and the complexity of fibre organizations in males, which would have an opposing impact on FA (the former should increase, while the latter should decrease FA). Together with non-significant trends for steeper ODI increase (and AD decrease) in males than females in several ROIs, it suggests a possibility that the FA is higher in males at an earlier age, but the faster increase in fibre complexity eventually cancels and even reverses the FA difference. However, such an explanation is not consistent with a finding of higher FA and lower ODI in males in mid- to late-adulthood (Ritchie et al., 2018). Further studies are needed to determine factors that may influence apparent sex differences in the WM properties in specific cohorts, such as body mass index, physical and intellectual activities, and other sex-related behavioural differences.

Regionally, we found the strongest sex differences in the brainstem ROIs. In addition to the overall patterns of higher diffusivity and lower NDI/ODI in females, females showed higher IsoVF than males in these ROIs. It suggests that the higher diffusivity in these ROIs in females can be partly attributed to the larger volume fraction for cerebrospinal fluid in the brainstem region of the female participants. Although this raises the question about the registration quality of the brainstem region, it is not clear why it would affect data from female subjects more than those from male subjects. Also, removing the outliers from data, thereby reducing the noise from the (most likely) poorly registered data, actually slightly increases the observed sex effects in the brainstem, rather than decreasing it (see Supplemental Material), arguing against the spurious sex differences in IsoVF caused by poor registration. Yet, if the differences are physiologically driven, their biological significance is not immediately clear. There is a paucity of comparable data from prior studies since the cerebellar and brainstem regions are often not targeted, and may not even be in the field of view of acquisition in many studies. Further studies are needed to replicate our findings and investigate the functional or biological significance of such sex differences.

### 4.3. Hemispheric asymmetries

While not our study’s primary focus, we observed surprisingly strong hemispheric asymmetry effects across many ROIs and metrics in our data. The direction of the asymmetry was mixed across ROIs for both WM regional volumes and microstructural properties. Even after correcting the ROI label size differences in the JHU ICBM-DTI-81 atlas, volumetric asymmetries were found across many ROIs. These volumetric asymmetries appeared largely independent from the asymmetries in the WM microstructural properties: The direct comparison of the standardised estimates for the hemisphere effects for volume and DTI/NODDI metrics across the 21 ROIs showed very little correlations, and controlling for the ROI volume difference in each hemisphere by including it as a covariate in the model did not affect the estimates for hemispheric effects for DTI/NODDI metrics. The relative independence of the macro- and microstructural asymmetries is consistent with previous work demonstrating the FA asymmetry to be largely independent of volumetric asymmetry (Powell et al., 2012; Takao et al., 2013). In contrast, within the hemispheric asymmetries of microstructural properties, the standardised parameter estimates of hemisphere for ODI showed a strong negative correlation with FA and AD, and a moderate positive correlation with RD. The degree of NDI asymmetry was not strongly related to the asymmetry in any other metrics.

Previous literature on WM asymmetry has focused mainly on the regional volumetric and FA asymmetry (e.g. Büchel et al., 2004; Hasan et al., 2010; Takao et al., 2011b; Thiebaut de Schotten et al., 2011; Powell et al., 2012; Takao et al., 2013). While these studies have revealed the hemispheric asymmetry in several WM regions, many of the findings are mixed and inconsistent (for the summary of key findings, see Table 1 in Honnedevasthana Arun et al., 2021). For example, even on the well-organised and circumscribed WM structure such as the posterior limb of the internal capsule, and using the same voxel-based approach in healthy adult samples, some studies have reported a rightward asymmetry for FA (Park et al., 2004), while others demonstrated a leftward FA asymmetry (Takao et al., 2011a; Honnedevasthana Arun et al., 2021). In addition, the limitations in the single tensor-based model make the inference of microstructural asymmetry based solely on FA values difficult; as noted earlier, a difference in FA can result from the asymmetry in a wide range of WM bundle properties, such as the neurite density, composition of fibre populations with different orientations, the degree of myelinations, etc.

The hemispheric asymmetry in NODDI metrics has not been investigated as much, and the few studies that examined the asymmetry are also inconsistent. One study in 104 infants under three months of age reported widespread hemispheric asymmetries for both NDI and ODI, with the varying directions of asymmetry across regions they examined (Dean et al., 2017), but another study in older children (between 4 - 8 years of age) found only rightward asymmetries for both NDI and ODI (Dimond et al., 2020). In two large-scale studies that examined both DTI and NODDI metrics in samples that include adult populations (3,513 subjects aged 44 - 77 years, Cox et al., 2016; 801 subjects aged 7 - 84 years, Slater et al., 2019) found the most robust hemispheric effects in FA and ODI, consistent with our findings. However, the effects appeared to be much more modest than what we observed in our data, although the difference in the definitions of ROIs and modelling strategies make the direct comparison difficult. In terms of the direction of the asymmetry, Cox et al. (2016) reported mostly rightward asymmetry for ODI (and correspondingly leftward asymmetry for FA), while Slater et al. (2019) reported more mixed asymmetries. Neither studies found very strong asymmetries for NDI, except for cingulum in the cingulate gyrus in Cox et al. study (but not in Slater et al. study), which showed higher NDI on the left than on the right. Notably, a leftward asymmetry for FA in the cingulum is relatively consistently found across many studies (Gong et al., 2005; Takao et al., 2011a; Cox et al., 2016; Slater et al., 2019; Bonekamp et al., 2007). A recent study using a fixel-based analytical approach found a leftward asymmetry for the “fibre density” parameter in the cingulum bundle as well, suggesting higher intra-axonal fibre volumes on the left side than the right (Honnedevasthana Arun et al., 2021). In our data, the cingulum in cingulate gyrus (CgC) ROI shows both higher NDI and lower ODI on the left compared to the right side. Since the lower degree of fibre complexity and the higher neurite density would both increase FA values, it points to the possible contribution of asymmetries in both of these microstructural properties to the FA asymmetry in this ROI observed in this and other studies.

Except for the IsoVF and MD asymmetry in a few ROIs, we found very little evidence of age-related variations in the hemispheric asymmetry effects overall. This is consistent with prior work that reports no or very little age-related changes in the hemispheric asymmetry patterns (Takao et al., 2011a; Lebel and Beaulieu, 2011; Slater et al., 2019; Dimond et al., 2020; Takao et al., 2013). Although it may indicate the emergence and establishment of hemispheric asymmetry very early in life (Song et al., 2015; Dean et al., 2017), clearly more work is needed, given the inconsistencies in the regional asymmetry patterns in WM properties in the literature. For example, the leftward asymmetry for FA in the cingulum bundle reported in many studies in adults has not been found earlier in development (Song et al., 2015; Dean et al., 2017; Dimond et al., 2020).

In addition to the main effects of the hemisphere, we found several ROIs with significant sex differences in the hemispheric asymmetries, most notably the fornix and stria terminalis ROI, which showed the sex and hemispheric interactions in several metrics, with non-negligible effect sizes. There have been few well-powered studies that examined the sex differences in the hemispheric asymmetry of the WM properties, and those reporting significant differences in either WM volumetric (Pujol et al., 2002; Savic, 2014) or FA asymmetry (Hasan et al., 2010; Powell et al., 2012; Takao et al., 2013) have conflicting findings. For example, Pujol et al. (2002) reported a stronger leftward asymmetry in males than in females for frontal WM volume in 100 healthy adults aged between 20 to 50 years, while Savic et al. (2014) reported a greater leftward asymmetry in males for WM volume around middle frontal gyrus in 86 adults with the similar age range. In 119 subjects from 7 to 68 years of age, Hasan et al. (2010) found a stronger leftward FA asymmetry in the inferior longitudinal fasciculus in males than females. In contrast, Powell et al. (2012) reported a stronger rightward FA asymmetry in males than females in the WM regions around the middle occipital lobe, middle temporal gyrus, and the supramarginal gyrus, in 82 young adults aged between 18 to 31 years. In a voxelwise analysis of WM skeleton in a larger sample of 857 healthy adults (age range of 25 to 85 years), Takao et al. (2013) reported a larger FA asymmetry in females in males in only 0.1% on the skeleton, and an even smaller proportion (0.04%) of WM skeleton with a larger FA asymmetry in males in females, and noted that the effect sizes for such differences were very small. Although we have higher power than most of these studies, given the finding in a relatively small ROI, it is important to confirm the robustness of the observed sex differences in the asymmetry to the different methodological approaches (discussed below).

### 4.4. Potential limitations

As we describe more in detail in Tsuchida *et al*. (2020), our sample from the MRi-Share database is drawn from students undergoing university-level education in Bordeaux, and as such, not necessarily a representative sample of healthy young adults. As a consequence, our sample is dominated by female participants, for example, and likely have different socio-demographic backgrounds and levels of education than the rest of the population of the same age range. They are also not guaranteed to be perfectly ‘healthy’, as the i-Share study, from which the MRi-Share participants were drawn, was designed to investigate the physical and mental health of students, and did not exclude those with past or current history of mental illness, alcohol intake, smoking habits, and/or use of any recreational drugs and psychotropic medications. While this undoubtedly increases the variance unaccounted for in our data, it also makes our data more representative of the sampled population.

The MRi-Share database is also currently cross-sectional, limiting our inference of maturational trajectory from the age-related variations in the data. The analysis of age effects based on cross-sectional data has been shown to lead to spurious findings unsupported from longitudinal analysis, especially when using quadratic models to describe non-linear patterns of age-related changes (Fjell et al., 2010; Pfefferbaum and Sullivan, 2015). In our sample with a relatively limited age range of 18 to 26 years, we found that linear age trends were sufficient for characterizing age-related variations in the data, thus avoiding some of the pitfalls associated with fitting quadratic age models. While we still need to exert caution when interpreting the apparent age-related variations in our data, our findings were found to be broadly consistent with the known age-related trajectories in WM properties.

In terms of the specific methodology for characterizing the regional WM properties, we used the ROIs based on the JHU ICBM-DTI-81 white matter labels atlas, computing the mean DTI/NODDI values within regions with high WM probability based on the multi-channel tissue segmentation with T1 and FLAIR scans. The ROIs in this atlas represent the WM regions with relatively well-organised structures that are clearly visible in the colour-coded map of the tensor fields, and should not be conflated with tracts obtained through tractography-based methods: The naming of these ROIs is based on the primary WM fibre population passing through the region, but these ROIs often represent a limited portion of a given tract, with arbitrary boundaries, and also may contain different fibre populations. For example, the corticospinal tract (CST) ROI in this atlas represents a portion of the CST at the level of medulla and pons, whereas the CST in the tractography-based methods usually refers to the fibre population that spans from corona radiata, passing through the internal capsule, then to the midbrain (Thiebaut de Schotten et al., 2011; Chenot et al., 2019). Another example is the sagittal stratum (SS) ROI, which, according to the authors of the atlas, includes both the inferior longitudinal fasciculus and the projection fibres from the internal capsule, therefore including both projection and association fibres (Mori et al., 2008). We also note that recent anatomical studies have seriously questioned the presence of superior fronto-occipital fasciculus (SFO) in humans (Türe et al., 1997; Forkel et al., 2014; Meola et al., 2015; Liu et al., 2020). Thus, this ROI most likely represents anterior thalamic radiation, as has been noted by the authors (Mori et al., 2008).

The anatomical definitions aside, all the WM microstructural properties we examined are sensitive to how the values are sampled. Most studies using the same JHU ICBM-DTI-81 atlas use it within the framework of Tract-Based Spatial Statistics (TBSS, Smith et al., 2006), included with the FSL package. With this approach, the analysis is limited to the one-voxel thick skeleton with maximal FA values, and the vast majority of WM regions surrounding it are ignored. Thus, by design, it biases the characterization of the WM microstructural properties to the small portion of WM, the “core” of the tracts with relatively simple fibre orientations (Lebel et al., 2019). The relative lack of age or sex effects for FA values in our study may be explained by the fact that we did not limit our analysis to the regions of high FA values, but sampled the mean DTI/NODDI values within the high WM probability region, based on the subject-specific tissue segmentation from the structural scans. In our view, it allows for a more complete characterization of the regional microstructural properties. Note that this approach is problematic mainly when making inferences about “WM integrity” based solely on the FA derived from a simple tensor model; the inclusion of voxels outside of the “core” of WM tracts increases the contribution of fibre geometry to the FA values, making it impossible to dissociate variations due to “WM integrity”, such as the axonal density and the degree of myelination, from those resulting from the complexity of orientations in the underlying fibre populations. Using less ambiguous metrics based on multi-component tissue models such as the NODDI allows for more specific inference about the variations or differences in the microstructural properties, as we demonstrated in this study. Our approach, in addition, allows for the direct comparison of the variations in the regional volume based on the Jacobian-modulated WM probability map and the variations in the microstructural properties in the same region.

Another critical difference between the TBSS-based approach and the current study is the method of spatial normalization: the TBSS projects the highest FA values onto a template FA skeleton in the standard space. Although it is meant to improve the alignment of the core of WM tracts, concerns have been raised with regard to the anatomical inaccuracies introduced by such a method (Bach et al., 2014). In the present study, we used the ‘Unified Segmentation’ framework (Ashburner and Friston, 2005) to perform spatial normalization based on tissue segmentation of the structural scans, a common approach in voxel-based morphometry studies (e.g. Takao et al., 2011a; Powell et al., 2012; Shiino et al., 2017). The non-linear deformation field obtained from the spatial normalization of the structural scans was then applied to DTI and NODDI maps, together with affine transformations that co-register these maps to the reference T1 scan of each subject. Although this is not necessarily the best available method to nonlinearly align images (Klein et al., 2009), we believe that the sampling and averaging of values within the regions comprising hundreds of voxels (or thousands, in many ROIs), defined based on both the template atlas label and subject-specific WM probability map, would limit the effects of small misalignments, especially with the large sample size in our study. Having said that, the robustness of the findings should be confirmed using different approaches, especially for the unexpected findings of hemispheric asymmetries. It is possible, for example, that some of the observed asymmetries in the ROIs surrounding the subcortical nuclei are due to the asymmetric partial volume effects caused by the asymmetry in these structures (Guadalupe et al., 2017).

### 4.5. Conclusions

In a large cohort of university students, we found a wide-spread increase in NDI, indicating a continuing increase in axonal/dendritic packing at this age range, with a more regionally-specific increase in ODI. Additionally, we observed substantial sex differences and hemispheric asymmetries across many WM regions in this cohort. We also demonstrated the distinct patterns of interrelations among the estimated effects on different WM properties for each effect. These findings highlight the complexity of the patterns of regional WM properties and individual variations in such patterns. Although we focused on the basic characterization of age and sex effects in the present study, they represent a small portion of the variance in data, and there are large individual differences in the regional WM volumes and microstructure. Future studies should investigate how the maturational processes in the WM influence, or are influenced by, cognitive, social, and behavioural factors, and how they are altered in neuropsychiatric conditions that manifest in early adulthood.

## Author Contributions

BM, CT, and SD contributed to conception and design of the study. AT and AL organised and processed imaging data to obtain IDPs described in the paper. AP and NB contributed to the QC of the IDPs. AT and BM performed the statistical analysis. AT and BM wrote the first draft of the manuscript. All authors contributed to manuscript revision, read, and approved the submitted version.

## Acknowledgements

The authors are indebted to Maxime Descoteaux (Sherbrooke University, Canada) for his help in implementing the DWI processing and QC pipelines.

## Data Availability

To access i-Share and MRi-Share de-identified data, a request can be submitted to the i-Share Scientific Collaborations Coordinator with a letter of intent (explaining the rationale and objectives of the research proposal), and a brief summary of the planned means and options for funding. The i-Share Steering Committee will assess this request, and provide a response (principle agreement, request to reformulate the application or for further information, refusal with reasons). If positive, applicants will have to complete and return an application package which will be reviewed by the principal investigator, the Steering Committee, and the operational staff. Reviews will be based on criteria such as the regulatory framework and adherence to regulations (access to data, confidentiality), the scientific and methodological quality of the project, the relevance of the project in relation to the overall consistency of the cohort in the long term, the complementarity/competition with projects planned or currently underway, ethical aspects. De-identified data (and data dictionaries) will be shared after (i) final approval of the application, and (ii) formalisation of the specifics of the collaboration.

