## Supplementary Material for "Changes in regional white matter volumetry and microstructure during the post-adolescence period: a cross-sectional study of a cohort of 1,713 university students"

### Table of Contents

**Table of Contents** 1

**Diffusion MRI Pipeline** 3

Supplemental Figure 1. Schematic of DWI pipeline 5

**Descriptive statistics of JHU IDPs** 5

Supplemental Table 1. Summary of descriptive statistics of JHU IDPs. 6

Supplemental Figure 2. Distribution of WM volumes in JHU ROIs. 9

Supplemental Figure 3. Distribution of WM FA in JHU ROIs. 10

Supplemental Figure 4. Distribution of WM MD in JHU ROIs. 11

Supplemental Figure 5. Distribution of WM AD in JHU ROIs. 12

Supplemental Figure 6. Distribution of WM RD in JHU ROIs. 13

Supplemental Figure 7. Distribution of WM NDI in JHU ROIs. 14

Supplemental Figure 8. Distribution of WM ODI in JHU ROIs. 15

Supplemental Figure 9. Distribution of WM IsoVF in JHU ROIs. 16

**Complete tables of model results for each metric** 17

Supplemental Table 2 Model results for WM JHU ROI volumes. 17

Supplemental Table 3. Model results for mean FA in JHU ROIs. 19

Supplemental Table 4. Model results for mean MD in JHU ROIs. 22

Supplemental Table 5. Model results for mean AD in JHU ROIs. 24

Supplemental Table 6. Model results for mean RD in JHU ROIs. 27

Supplemental Table 7. Model results for mean NDI in JHU ROIs. 30

Supplemental Table 8. Model results for mean ODI in JHU ROIs. 32

Supplemental Table 9. Model results for mean IsoVF in JHU ROIs. 35

**Summary figures for age, sex, and hemispheric asymmetry effects** 37

Supplemental Figure 10. Effect size heatmaps for age, sex, and hemispheric asymmetry. 38

Supplemental Figure 11. Scatter plots of individual age effects on WM volumes in each ROI. 39

Supplemental Figure 12. Scatter plots of individual age effects on WM FA values in each ROI. 40

Supplemental Figure 13. Scatter plots of individual age effects on WM MD values in each ROI. 41

Supplemental Figure 14. Scatter plots of individual age effects on WM AD values in each ROI. 42

Supplemental Figure 15. Scatter plots of individual age effects on WM RD values in each ROI. 43

Supplemental Figure 16. Scatter plots of individual age effects on WM NDI values in each ROI. 44

Supplemental Figure 17. Scatter plots of individual age effects on WM ODI values in each ROI. 45
Supplemental Figure 18. Scatter plots of individual age effects on WM IsoVF values in each ROI. 46
Supplemental Figure 19. Line plots of individual hemisphere effects on WM volumes in each ROI. 47
Supplemental Figure 20. Line plots of individual hemisphere effects on WM FA values in each ROI. 48
Supplemental Figure 21. Line plots of individual hemisphere effects on WM MD values in each ROI. 49
Supplemental Figure 22. Line plots of individual hemisphere effects on WM AD values in each ROI. 50
Supplemental Figure 23. Line plots of individual hemisphere effects on WM RD values in each ROI. 51
Supplemental Figure 24. Line plots of individual hemisphere effects on WM NDI values in each ROI. 52
Supplemental Figure 25. Line plots of individual hemisphere effects on WM ODI values in each ROI. 53
Supplemental Figure 26. Line plots of individual hemisphere effects on WM IsoVF values in each ROI. 54

**Effects of outliers on the estimates of age, sex, and hemispheric asymmetry** 55
Supplemental Figure 27. Impact of outlier removal on the standardized estimates of age, sex, and hemispheric asymmetry effects on the different WM IDPs in the JHU ROIs. 56

**Effects of motion parameters during the DWI scan** 57
Supplemental Figure 28. Comparison of model quality between models with and without RMS for the DTI/NODDI IDPs and JHU ROIs. 58
Supplemental Figure 29. In-scanner motion effects and their impact on the estimates of age and sex effects for the different DTI/NODDI IDPs and JHU ROIs. 58

**Comparison of quadratic versus linear age effect models** 59
Supplemental Figure 30. Comparison of linear and quadratic age model quality for the different WM IDPs and JHU ROIs. 59

**Effects of volume corrections** 60
Effects of global volume correction on regional WM volumetry 60
Supplemental Figure 31. Effects of global volume correction on the model quality and the standardized estimates of age and sex effects in the WM JHU ROI volumes. 61
Effects of global and regional volume correction on DTI/NODDI metrics 62
Supplemental Figure 32. Comparisons of model quality for models with or without global and/or local volume correction for the mean DTI/NODDI values in JHU ROIs. 63

Supplemental Figure 33. Effects of global or ROI volumes on the mean DTI/NODDI values and their impact on the estimated age, sex, and hemisphere effects. 64

**References** 65

### Diffusion MRI Pipeline

Supplemental Figure 1 summarizes the DWI processing pipeline. In this pipeline, individual raw DWI data are first corrected for eddy current (EC) and top-up distortion using the FSL Eddy tool, with replacement of outlier slices (eddy_openmp as implemented in FSL v5.0.10 patch; (Andersson et al., 2016; Andersson and Sotiropoulos, 2016). After cropping the data to reduce non-brain tissue volumes, we apply non-local means filter to denoise and boost the SNR, using the 'nlmeans' denoising tool (Coupe et al., 2008, 2011) as implemented in the Dipy package (0.12.0; Garyfallidis et al., 2014). The resulting image is then used to estimate 1) DTI (Diffusion-Tensor Imaging; Basser et al., 1994) model parameters and 2) microstructural NODDI (Neurite Orientation Dispersion and Density Imaging; Zhang et al., 2012) model parameters. For DTI modelling, the volumes with high b-value (b=2000 s/mm$^2$) are removed from the denoised data before fitting the data with the dipy tools (Jensen and Helpern, 2010) to compute DTI maps, namely the maps of fractional anisotropy (FA), mean, axial, and radial diffusivity (MD, AD, and RD, respectively). The diffusivity maps were further cleaned by removing diffusivity value outliers using Random Sample Consensus (RANSAC) approach (Choi et al., 2009), as implemented in the scikit-learn package (0.19.1; https://scikit-learn.org/stable/index.html). The denoising, DTI computation, and the RANSAC outlier removal were performed by wrapping Scipy scripts, developed by Sherbrooke Connectivity Imaging Lab (https://scilpy.readthedocs.io/en/latest/).

Before fitting denoised data for NODDI, we computed empirical values of cohort-specific isotropic and parallel diffusivity by computing the mean MD within lateral ventricles and mean AD within the corpus callosum in individual T1 space for each subject. The mean of these values across subjects was then used as dPar (1.5 x $10^{-3}$) and dIso (2.4 x $10^{-3}$) parameters in the AMICO (Accelerated Microstructure Imaging via Convex Optimization) tool (Daducci et al., 2015) when fitting the denoised DWI data to obtain the following NODDI metric maps: isotropic volume fraction (IsoVF), which indicates the proportion of free water volume of each voxel, neurite density index (NDI), which represents the proportion of intracellular volume in the remaining fraction, and orientation dispersion index (ODI), a measure of within-voxel fibre dispersion.

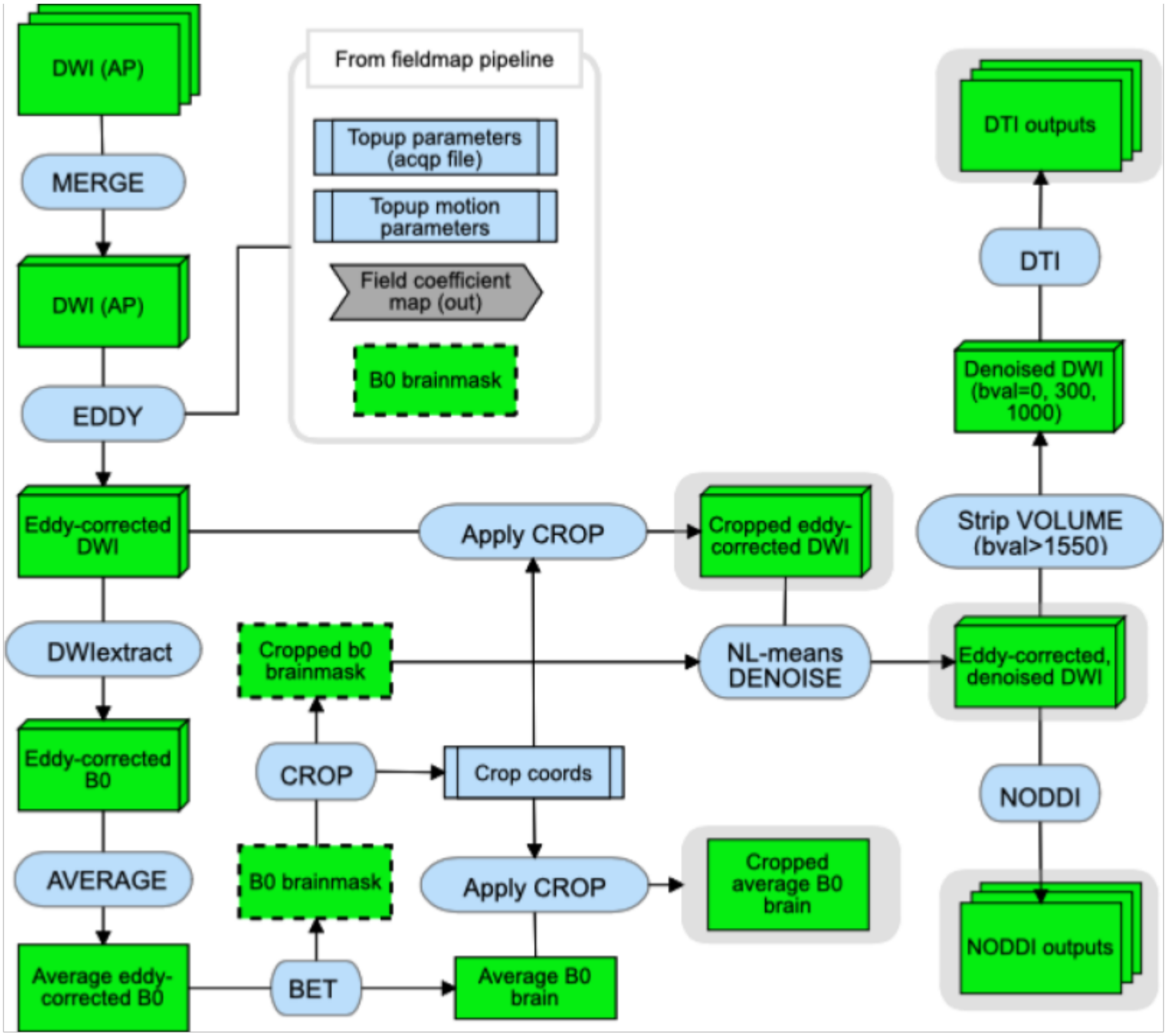

**Supplemental Figure 1. Schematic of DWI pipeline**

### Descriptive statistics of JHU IDPs

Supplemental Table 1 summarizes the descriptive statistics of the WM volume and mean DTI/NODDI values within each of the 48 ROIs in JHU ICBM-DTI-81 atlas. The distributions of each metric in each ROI are also visualized in the Supplemental Figures 2 - 9. With the exception of WM volume distributions (Figure S2) that are organized by ROI size, JHU ROIs in these figures are grouped according to the type of main fibers they contain, namely those in brainstem, those that are primarily projection, association, or commissural fibers. Red data points in the distribution figures are extreme outliers, defined as those below or above

3*interquartile range (IQR) from the first or third quartile, respectively. It can be seen that there are many (>10) extreme outliers for mean values of diffusivity (MD, AD, and RD) and NODDI (NDI, ODI, IsoVF) metrics in very small ROIs (e.g. fornix (FX) and tapetum (TAP L/R) as well as brainstem ROIs. It underscores the susceptibility of these ROIs to the slight misalignments between the DTI/NODDI maps and the T1w image, which can increase partial volume effects (PVEs) from non-WM tissue, in particular cerebrospinal fluid.

**Supplemental Table 1. Summary of descriptive statistics of JHU IDPs.**

Values are sample mean, standard deviation (in parentheses), and range (in square brackets). The first columns give the abbreviated JHU ROIs (see Table 2 of the main text for their full names). L: left, R: right.

| | Volume ($x10^3$ $mm^3$) | FA | MD ($x10^{-4}$ $mm^2$/sec) | AD ($x10^{-4}$ $mm^2$/sec) | RD ($x10^{-4}$ $mm^2$/sec) | NDI | ODI | IsoVF |
|---|---|---|---|---|---|---|---|---|
| Brainstem | | | | | | | | |
| MCP | 11.5 (1.2) [ 7.6, 16.5] | 0.51 (0.02) [ 0.44, 0.57] | 6.5 (0.3) [ 5.7, 7.5] | 10.6 (0.4) [ 9.1, 12.1] | 4.5 (0.2) [ 3.8, 5.5] | 0.89 (0.03) [ 0.76, 0.98] | 0.28 (0.02) [ 0.23, 0.37] | 0.19 (0.02) [ 0.11, 0.27] |
| PCT | 1.3 (0.1) [ 0.9, 1.8] | 0.41 (0.04) [ 0.29, 0.53] | 6.7 (0.7) [ 5.1, 10.8] | 9.7 (0.9) [ 7.5, 14.7] | 5.1 (0.7) [ 3.8, 8.9] | 0.96 (0.03) [ 0.70, 0.99] | 0.36 (0.04) [ 0.26, 0.52] | 0.22 (0.06) [ 0.06, 0.50] |
| CST L | 1.1 (0.1) [ 0.8, 1.5] | 0.48 (0.04) [ 0.33, 0.62] | 7.1 (0.7) [ 5.6, 10.9] | 11.0 (0.9) [ 8.5, 15.2] | 5.1 (0.7) [ 3.8, 9.0] | 0.92 (0.05) [ 0.66, 0.99] | 0.31 (0.05) [ 0.20, 0.57] | 0.23 (0.06) [ 0.10, 0.55] |
| CST R | 1.1 (0.1) [ 0.8, 1.5] | 0.47 (0.04) [ 0.30, 0.61] | 7.1 (0.8) [ 5.4, 11.4] | 10.9 (1.0) [ 8.2, 16.3] | 5.2 (0.8) [ 3.5, 9.4] | 0.91 (0.05) [ 0.61, 0.99] | 0.32 (0.05) [ 0.21, 0.61] | 0.23 (0.06) [ 0.07, 0.57] |
| ML L | 0.5 (0.1) [ 0.4, 0.7] | 0.54 (0.05) [ 0.28, 0.64] | 7.2 (0.7) [ 5.6, 15.1] | 11.9 (0.9) [ 9.2, 18.8] | 4.8 (0.8) [ 3.5, 13.2] | 0.82 (0.07) [ 0.38, 0.98] | 0.25 (0.05) [ 0.14, 0.71] | 0.21 (0.05) [ 0.10, 0.71] |
| ML R | 0.5 (0.1) [ 0.4, 0.7] | 0.54 (0.05) [ 0.27, 0.65] | 7.0 (0.7) [ 5.5, 13.7] | 11.7 (0.8) [ 9.3, 17.0] | 4.7 (0.7) [ 3.3, 12.2] | 0.84 (0.06) [ 0.48, 0.98] | 0.25 (0.05) [ 0.15, 0.66] | 0.21 (0.05) [ 0.09, 0.64] |
| SCP L | 0.4 (0.1) [ 0.2, 0.6] | 0.65 (0.03) [ 0.45, 0.75] | 6.8 (0.3) [ 5.8, 8.5] | 13.0 (0.7) [10.2, 15.3] | 3.7 (0.3) [ 2.7, 5.3] | 0.88 (0.04) [ 0.73, 1.00] | 0.17 (0.02) [ 0.11, 0.32] | 0.18 (0.02) [ 0.09, 0.31] |
| SCP R | 0.4 (0.1) [ 0.2, 0.6] | 0.65 (0.03) [ 0.54, 0.74] | 6.7 (0.3) [ 5.5, 8.2] | 12.9 (0.7) [ 9.7, 14.9] | 3.7 (0.3) [ 2.8, 5.1] | 0.89 (0.04) [ 0.74, 1.00] | 0.17 (0.02) [ 0.11, 0.29] | 0.18 (0.02) [ 0.08, 0.31] |
| ICP L | 0.6 (0.1) [ 0.3, 0.9] | 0.51 (0.03) [ 0.31, 0.61] | 7.1 (0.5) [ 6.2, 12.1] | 11.6 (0.6) [10.0, 16.1] | 4.9 (0.5) [ 3.9, 10.2] | 0.83 (0.05) [ 0.54, 0.98] | 0.25 (0.03) [ 0.17, 0.54] | 0.22 (0.04) [ 0.11, 0.56] |
| ICP R | 0.6 (0.1) [ 0.4, 0.8] | 0.50 (0.03) [ 0.31, 0.60] | 7.1 (0.4) [ 5.9, 9.3] | 11.5 (0.5) [ 9.7, 13.8] | 4.9 (0.4) [ 3.9, 7.1] | 0.85 (0.05) [ 0.68, 0.99] | 0.25 (0.03) [ 0.17, 0.48] | 0.22 (0.03) [ 0.13, 0.38] |
| Projection | | | | | | | | |
| ACR L | 5.4 (0.6) [ 3.8, 7.8] | 0.40 (0.03) [ 0.32, 0.56] | 7.2 (0.2) [ 6.5, 7.9] | 10.5 (0.4) [ 9.6, 12.7] | 5.5 (0.3) [ 4.7, 6.4] | 0.66 (0.04) [ 0.52, 0.80] | 0.28 (0.02) [ 0.19, 0.35] | 0.17 (0.01) [ 0.12, 0.22] |

| | | | | | | | | |
|---|---|---|---|---|---|---|---|---|
| ACR R | 5.4 (0.5)<br>[ 3.9, 7.6] | 0.40 (0.03)<br>[ 0.30, 0.50] | 7.2 (0.2)<br>[ 6.6, 7.9] | 10.5 (0.4)<br>[ 9.5, 11.9] | 5.5 (0.3)<br>[ 4.7, 6.4] | 0.66 (0.04)<br>[ 0.53, 0.80] | 0.28 (0.02)<br>[ 0.21, 0.36] | 0.17 (0.01)<br>[ 0.12, 0.23] |
| SCR L | 5.8 (0.6)<br>[ 4.3, 8.3] | 0.45 (0.03)<br>[ 0.36, 0.63] | 6.7 (0.2)<br>[ 6.2, 7.4] | 10.2 (0.3)<br>[ 9.2, 12.8] | 5.0 (0.2)<br>[ 3.9, 5.8] | 0.74 (0.03)<br>[ 0.63, 0.85] | 0.27 (0.02)<br>[ 0.14, 0.35] | 0.18 (0.01)<br>[ 0.12, 0.22] |
| SCR R | 5.8 (0.5)<br>[ 4.3, 8.1] | 0.44 (0.03)<br>[ 0.35, 0.58] | 6.7 (0.2)<br>[ 6.1, 7.4] | 10.1 (0.3)<br>[ 9.0, 12.2] | 5.0 (0.2)<br>[ 4.2, 5.8] | 0.74 (0.03)<br>[ 0.62, 0.86] | 0.27 (0.02)<br>[ 0.16, 0.34] | 0.18 (0.01)<br>[ 0.11, 0.22] |
| PCR L | 2.8 (0.3)<br>[ 2.1, 4.1] | 0.44 (0.03)<br>[ 0.33, 0.61] | 7.3 (0.2)<br>[ 6.5, 8.2] | 11.1 (0.4)<br>[ 9.8, 14.0] | 5.5 (0.3)<br>[ 4.5, 6.5] | 0.67 (0.04)<br>[ 0.54, 0.79] | 0.24 (0.02)<br>[ 0.13, 0.35] | 0.19 (0.02)<br>[ 0.13, 0.25] |
| PCR R | 2.8 (0.3)<br>[ 2.1, 4.0] | 0.46 (0.03)<br>[ 0.36, 0.59] | 7.4 (0.3)<br>[ 6.7, 8.4] | 11.4 (0.4)<br>[10.1, 13.9] | 5.4 (0.3)<br>[ 4.5, 6.6] | 0.65 (0.04)<br>[ 0.53, 0.83] | 0.22 (0.02)<br>[ 0.14, 0.32] | 0.19 (0.02)<br>[ 0.12, 0.27] |
| ALIC L | 1.9 (0.2)<br>[ 1.3, 2.7] | 0.52 (0.02)<br>[ 0.43, 0.59] | 6.7 (0.2)<br>[ 6.1, 7.5] | 11.1 (0.3)<br>[10.0, 12.3] | 4.4 (0.2)<br>[ 3.7, 5.3] | 0.75 (0.04)<br>[ 0.62, 0.87] | 0.24 (0.02)<br>[ 0.18, 0.32] | 0.14 (0.02)<br>[ 0.10, 0.21] |
| ALIC R | 2.0 (0.2)<br>[ 1.5, 3.0] | 0.54 (0.02)<br>[ 0.46, 0.62] | 6.8 (0.2)<br>[ 6.1, 7.3] | 11.4 (0.3)<br>[10.0, 12.8] | 4.4 (0.2)<br>[ 3.8, 5.2] | 0.76 (0.04)<br>[ 0.64, 0.90] | 0.21 (0.02)<br>[ 0.16, 0.28] | 0.16 (0.02)<br>[ 0.11, 0.22] |
| PLIC L | 3.2 (0.3)<br>[ 2.3, 4.6] | 0.59 (0.02)<br>[ 0.52, 0.70] | 6.5 (0.2)<br>[ 5.9, 7.0] | 11.6 (0.3)<br>[10.4, 13.3] | 4.0 (0.2)<br>[ 3.3, 4.7] | 0.84 (0.04)<br>[ 0.71, 0.97] | 0.20 (0.01)<br>[ 0.13, 0.26] | 0.17 (0.01)<br>[ 0.11, 0.22] |
| PLIC R | 3.1 (0.3)<br>[ 2.3, 4.5] | 0.58 (0.02)<br>[ 0.50, 0.66] | 6.6 (0.2)<br>[ 6.0, 7.1] | 11.5 (0.3)<br>[10.1, 12.8] | 4.1 (0.2)<br>[ 3.5, 4.8] | 0.84 (0.04)<br>[ 0.68, 0.98] | 0.21 (0.02)<br>[ 0.15, 0.28] | 0.18 (0.01)<br>[ 0.11, 0.23] |
| RLIC L | 1.9 (0.2)<br>[ 1.4, 2.8] | 0.61 (0.02)<br>[ 0.53, 0.70] | 7.0 (0.2)<br>[ 6.3, 7.6] | 12.5 (0.4)<br>[11.1, 13.5] | 4.2 (0.2)<br>[ 3.3, 5.0] | 0.76 (0.04)<br>[ 0.63, 0.90] | 0.17 (0.01)<br>[ 0.13, 0.23] | 0.17 (0.02)<br>[ 0.10, 0.23] |
| RLIC R | 2.0 (0.2)<br>[ 1.4, 2.7] | 0.57 (0.02)<br>[ 0.49, 0.66] | 7.1 (0.2)<br>[ 6.2, 7.9] | 12.2 (0.4)<br>[10.8, 13.3] | 4.5 (0.2)<br>[ 3.7, 5.3] | 0.78 (0.04)<br>[ 0.63, 0.95] | 0.19 (0.02)<br>[ 0.14, 0.25] | 0.20 (0.02)<br>[ 0.11, 0.26] |
| PTR L | 2.9 (0.3)<br>[ 1.9, 4.1] | 0.61 (0.03)<br>[ 0.51, 0.70] | 7.5 (0.2)<br>[ 6.9, 8.4] | 13.5 (0.4)<br>[12.0, 14.8] | 4.5 (0.3)<br>[ 3.5, 5.7] | 0.65 (0.03)<br>[ 0.53, 0.78] | 0.14 (0.01)<br>[ 0.10, 0.19] | 0.17 (0.01)<br>[ 0.10, 0.24] |
| PTR R | 3.0 (0.3)<br>[ 2.2, 4.2] | 0.59 (0.03)<br>[ 0.49, 0.67] | 7.6 (0.2)<br>[ 6.8, 8.4] | 13.6 (0.4)<br>[12.1, 14.9] | 4.7 (0.3)<br>[ 3.9, 5.7] | 0.64 (0.04)<br>[ 0.53, 0.91] | 0.14 (0.01)<br>[ 0.10, 0.20] | 0.18 (0.01)<br>[ 0.11, 0.26] |
| CP L | 1.7 (0.2)<br>[ 1.3, 2.4] | 0.61 (0.03)<br>[ 0.49, 0.71] | 7.4 (0.7)<br>[ 5.5, 9.9] | 13.0 (0.9)<br>[10.2, 15.9] | 4.5 (0.7)<br>[ 2.7, 7.1] | 0.85 (0.06)<br>[ 0.62, 0.99] | 0.23 (0.03)<br>[ 0.14, 0.36] | 0.22 (0.04)<br>[ 0.09, 0.40] |
| CP R | 1.7 (0.2)<br>[ 1.2, 2.4] | 0.60 (0.03)<br>[ 0.47, 0.70] | 7.3 (0.7)<br>[ 5.4, 10.8] | 12.9 (0.9)<br>[10.1, 16.9] | 4.5 (0.7)<br>[ 2.9, 7.8] | 0.86 (0.06)<br>[ 0.55, 0.98] | 0.23 (0.03)<br>[ 0.14, 0.39] | 0.22 (0.04)<br>[ 0.08, 0.43] |
| Association | | | | | | | | |
| FX | 0.2 (0.0)<br>[ 0.0, 0.3] | 0.74 (0.03)<br>[ 0.43, 0.83] | 8.0 (0.5)<br>[ 6.9, 15.9] | 16.7 (0.7)<br>[14.6, 23.4] | 3.7 (0.6)<br>[ 2.3, 12.1] | 0.71 (0.06)<br>[ 0.36, 0.87] | 0.06 (0.04)<br>[-0.02, 0.60] | 0.20 (0.05)<br>[ 0.09, 0.77] |
| FX/ST L | 0.7 (0.1)<br>[ 0.4, 1.0] | 0.60 (0.03)<br>[ 0.48, 0.70] | 7.0 (0.2)<br>[ 6.2, 7.7] | 12.4 (0.6)<br>[10.2, 14.4] | 4.3 (0.3)<br>[ 3.4, 5.3] | 0.80 (0.05)<br>[ 0.65, 0.96] | 0.19 (0.02)<br>[ 0.12, 0.29] | 0.18 (0.02)<br>[ 0.11, 0.25] |
| FX/ST R | 0.5 (0.1)<br>[ 0.3, 0.7] | 0.60 (0.03)<br>[ 0.49, 0.70] | 7.3 (0.3)<br>[ 6.2, 8.1] | 13.0 (0.6)<br>[10.7, 15.0] | 4.4 (0.3)<br>[ 3.4, 5.5] | 0.81 (0.05)<br>[ 0.65, 0.96] | 0.18 (0.02)<br>[ 0.12, 0.27] | 0.21 (0.02)<br>[ 0.11, 0.29] |
| CgC L | 1.6 (0.2)<br>[ 0.9, 2.5] | 0.55 (0.03)<br>[ 0.45, 0.64] | 7.0 (0.2)<br>[ 6.3, 7.6] | 11.8 (0.4)<br>[10.2, 13.3] | 4.5 (0.3)<br>[ 3.7, 5.4] | 0.74 (0.04)<br>[ 0.61, 0.86] | 0.21 (0.02)<br>[ 0.14, 0.30] | 0.16 (0.02)<br>[ 0.11, 0.21] |
| CgC R | 1.3 (0.2)<br>[ 0.8, 2.1] | 0.52 (0.03)<br>[ 0.41, 0.60] | 7.0 (0.2)<br>[ 6.3, 7.6] | 11.3 (0.4)<br>[ 9.8, 12.8] | 4.8 (0.3)<br>[ 4.0, 5.6] | 0.73 (0.04)<br>[ 0.60, 0.88] | 0.24 (0.02)<br>[ 0.16, 0.33] | 0.17 (0.02)<br>[ 0.12, 0.22] |

| | | | | | | | | |
|---|---|---|---|---|---|---|---|---|
| CgH L | 0.6 (0.1)<br>[ 0.3, 0.9] | 0.49 (0.04)<br>[ 0.35, 0.60] | 7.0 (0.3)<br>[ 5.9, 7.9] | 11.3 (0.5)<br>[ 9.2, 12.8] | 4.9 (0.3)<br>[ 3.9, 6.0] | 0.75 (0.07)<br>[ 0.56, 0.96] | 0.25 (0.03)<br>[ 0.16, 0.43] | 0.17 (0.02)<br>[ 0.09, 0.26] |
| CgH R | 0.6 (0.1)<br>[ 0.4, 1.0] | 0.48 (0.03)<br>[ 0.27, 0.61] | 7.1 (0.3)<br>[ 6.0, 9.3] | 11.3 (0.5)<br>[ 9.3, 12.8] | 5.0 (0.3)<br>[ 3.9, 8.0] | 0.75 (0.07)<br>[ 0.53, 0.96] | 0.26 (0.03)<br>[ 0.17, 0.49] | 0.18 (0.03)<br>[ 0.07, 0.31] |
| SFO L | 0.3 (0.0)<br>[ 0.2, 0.5] | 0.43 (0.04)<br>[ 0.31, 0.56] | 6.9 (0.2)<br>[ 6.0, 7.7] | 10.5 (0.4)<br>[ 8.8, 12.1] | 5.1 (0.3)<br>[ 4.0, 6.3] | 0.76 (0.05)<br>[ 0.60, 0.89] | 0.29 (0.03)<br>[ 0.19, 0.40] | 0.18 (0.02)<br>[ 0.11, 0.29] |
| SFO R | 0.3 (0.0)<br>[ 0.2, 0.5] | 0.44 (0.04)<br>[ 0.32, 0.59] | 6.9 (0.2)<br>[ 6.1, 7.9] | 10.6 (0.4)<br>[ 9.0, 12.2] | 5.1 (0.3)<br>[ 3.9, 6.1] | 0.75 (0.05)<br>[ 0.63, 0.91] | 0.28 (0.03)<br>[ 0.19, 0.39] | 0.19 (0.02)<br>[ 0.12, 0.28] |
| SLF L | 5.0 (0.5)<br>[ 3.6, 7.0] | 0.49 (0.02)<br>[ 0.40, 0.57] | 6.7 (0.2)<br>[ 6.3, 7.5] | 10.6 (0.3)<br>[ 9.6, 11.6] | 4.8 (0.2)<br>[ 4.1, 5.7] | 0.73 (0.03)<br>[ 0.62, 0.84] | 0.24 (0.02)<br>[ 0.20, 0.30] | 0.17 (0.01)<br>[ 0.13, 0.21] |
| SLF R | 5.0 (0.5)<br>[ 3.5, 6.9] | 0.49 (0.03)<br>[ 0.40, 0.58] | 6.8 (0.2)<br>[ 6.3, 7.6] | 10.8 (0.3)<br>[ 9.7, 11.8] | 4.9 (0.2)<br>[ 4.1, 5.7] | 0.72 (0.03)<br>[ 0.61, 0.86] | 0.24 (0.02)<br>[ 0.18, 0.30] | 0.17 (0.01)<br>[ 0.13, 0.22] |
| EC L | 2.4 (0.3)<br>[ 1.6, 3.6] | 0.51 (0.02)<br>[ 0.44, 0.59] | 7.0 (0.2)<br>[ 6.4, 7.7] | 11.3 (0.3)<br>[10.4, 12.2] | 4.8 (0.2)<br>[ 4.1, 5.6] | 0.67 (0.03)<br>[ 0.55, 0.79] | 0.22 (0.02)<br>[ 0.17, 0.28] | 0.14 (0.02)<br>[ 0.08, 0.21] |
| EC R | 2.2 (0.3)<br>[ 1.4, 3.2] | 0.49 (0.02)<br>[ 0.43, 0.58] | 7.1 (0.2)<br>[ 6.6, 7.7] | 11.3 (0.3)<br>[10.3, 12.2] | 5.0 (0.2)<br>[ 4.3, 5.6] | 0.68 (0.03)<br>[ 0.58, 0.84] | 0.23 (0.02)<br>[ 0.18, 0.28] | 0.16 (0.02)<br>[ 0.09, 0.26] |
| UNC L | 0.2 (0.0)<br>[ 0.0, 0.3] | 0.60 (0.04)<br>[ 0.36, 0.73] | 7.1 (0.2)<br>[ 6.5, 8.0] | 12.7 (0.6)<br>[10.1, 14.5] | 4.3 (0.4)<br>[ 3.0, 6.1] | 0.65 (0.05)<br>[ 0.50, 0.82] | 0.15 (0.02)<br>[ 0.09, 0.34] | 0.12 (0.02)<br>[ 0.05, 0.23] |
| UNC R | 0.2 (0.0)<br>[ 0.0, 0.3] | 0.59 (0.04)<br>[ 0.44, 0.69] | 7.3 (0.2)<br>[ 6.5, 8.3] | 12.9 (0.4)<br>[11.5, 14.4] | 4.5 (0.3)<br>[ 3.4, 5.7] | 0.62 (0.04)<br>[ 0.49, 0.82] | 0.15 (0.02)<br>[ 0.10, 0.22] | 0.13 (0.03)<br>[ 0.03, 0.27] |
| SS L | 1.6 (0.2)<br>[ 1.1, 2.3] | 0.58 (0.03)<br>[ 0.48, 0.69] | 7.4 (0.2)<br>[ 6.6, 8.4] | 12.8 (0.4)<br>[11.4, 14.2] | 4.7 (0.3)<br>[ 3.6, 6.0] | 0.70 (0.04)<br>[ 0.57, 0.85] | 0.18 (0.02)<br>[ 0.13, 0.24] | 0.19 (0.01)<br>[ 0.12, 0.27] |
| SS R | 1.7 (0.2)<br>[ 1.2, 2.3] | 0.56 (0.03)<br>[ 0.47, 0.66] | 7.5 (0.2)<br>[ 6.7, 8.4] | 12.8 (0.4)<br>[11.7, 14.2] | 4.9 (0.3)<br>[ 4.0, 6.1] | 0.69 (0.05)<br>[ 0.55, 0.92] | 0.18 (0.02)<br>[ 0.14, 0.23] | 0.20 (0.02)<br>[ 0.12, 0.31] |
| Commissural | | | | | | | | |
| GCC | 5.5 (0.7)<br>[ 2.4, 8.2] | 0.64 (0.03)<br>[ 0.45, 0.72] | 7.2 (0.2)<br>[ 6.4, 7.9] | 13.6 (0.4)<br>[11.3, 15.0] | 4.0 (0.3)<br>[ 3.2, 5.3] | 0.73 (0.04)<br>[ 0.61, 0.88] | 0.15 (0.02)<br>[ 0.10, 0.29] | 0.17 (0.01)<br>[ 0.12, 0.22] |
| BCC | 8.6 (1.0)<br>[ 5.8, 12.4] | 0.65 (0.02)<br>[ 0.55, 0.73] | 7.1 (0.2)<br>[ 6.5, 7.8] | 13.7 (0.3)<br>[12.2, 14.8] | 3.8 (0.2)<br>[ 3.0, 4.9] | 0.75 (0.04)<br>[ 0.63, 0.87] | 0.14 (0.01)<br>[ 0.10, 0.20] | 0.17 (0.01)<br>[ 0.12, 0.23] |
| SCC | 8.5 (1.1)<br>[ 5.5, 12.7] | 0.70 (0.02)<br>[ 0.57, 0.78] | 7.0 (0.2)<br>[ 6.3, 7.7] | 14.1 (0.4)<br>[12.6, 15.6] | 3.5 (0.2)<br>[ 2.8, 4.5] | 0.82 (0.04)<br>[ 0.68, 0.93] | 0.13 (0.01)<br>[ 0.09, 0.20] | 0.18 (0.01)<br>[ 0.12, 0.23] |
| TAP L | 0.2 (0.1)<br>[ 0.0, 0.4] | 0.61 (0.05)<br>[ 0.37, 0.76] | 8.1 (0.6)<br>[ 6.7, 11.7] | 14.7 (0.9)<br>[11.9, 18.5] | 4.8 (0.6)<br>[ 3.1, 9.0] | 0.59 (0.06)<br>[ 0.34, 0.80] | 0.14 (0.04)<br>[ 0.04, 0.39] | 0.20 (0.04)<br>[ 0.08, 0.50] |
| TAP R | 0.2 (0.1)<br>[ 0.0, 0.4] | 0.62 (0.05)<br>[ 0.44, 0.75] | 8.0 (0.4)<br>[ 6.9, 11.2] | 14.6 (0.8)<br>[12.6, 18.2] | 4.7 (0.5)<br>[ 3.2, 7.8] | 0.58 (0.05)<br>[ 0.37, 0.82] | 0.12 (0.03)<br>[ 0.05, 0.38] | 0.18 (0.03)<br>[ 0.08, 0.47] |

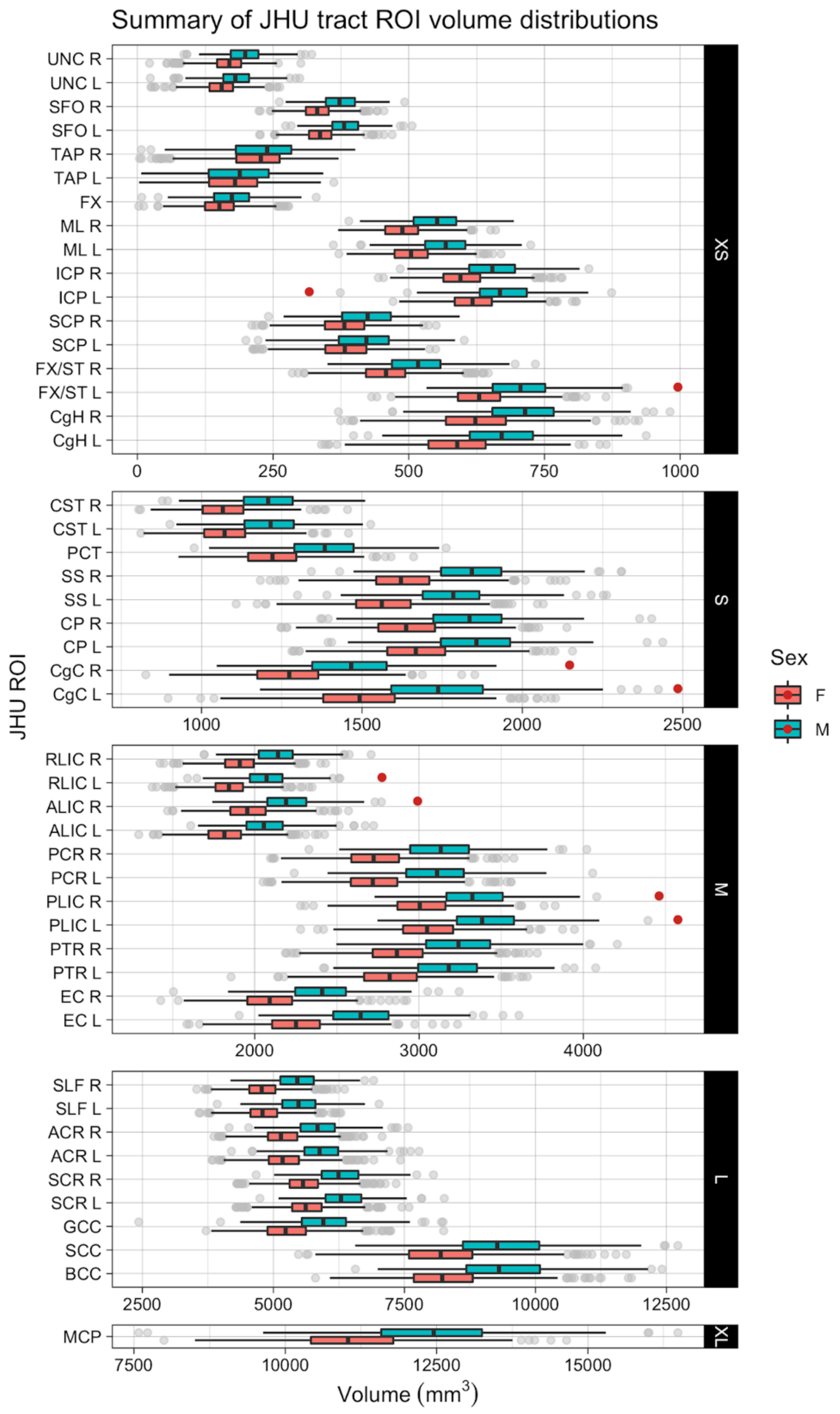

**Supplemental Figure 2. Distribution of WM volumes in JHU ROIs.**

Boxplots of regional white matter volumes for each sex in each JHU ROI. See Table 2 of the main text for the full names of the abbreviated ROIs. L: left, R: right. F: female, M: male. Red dots indicate outliers.

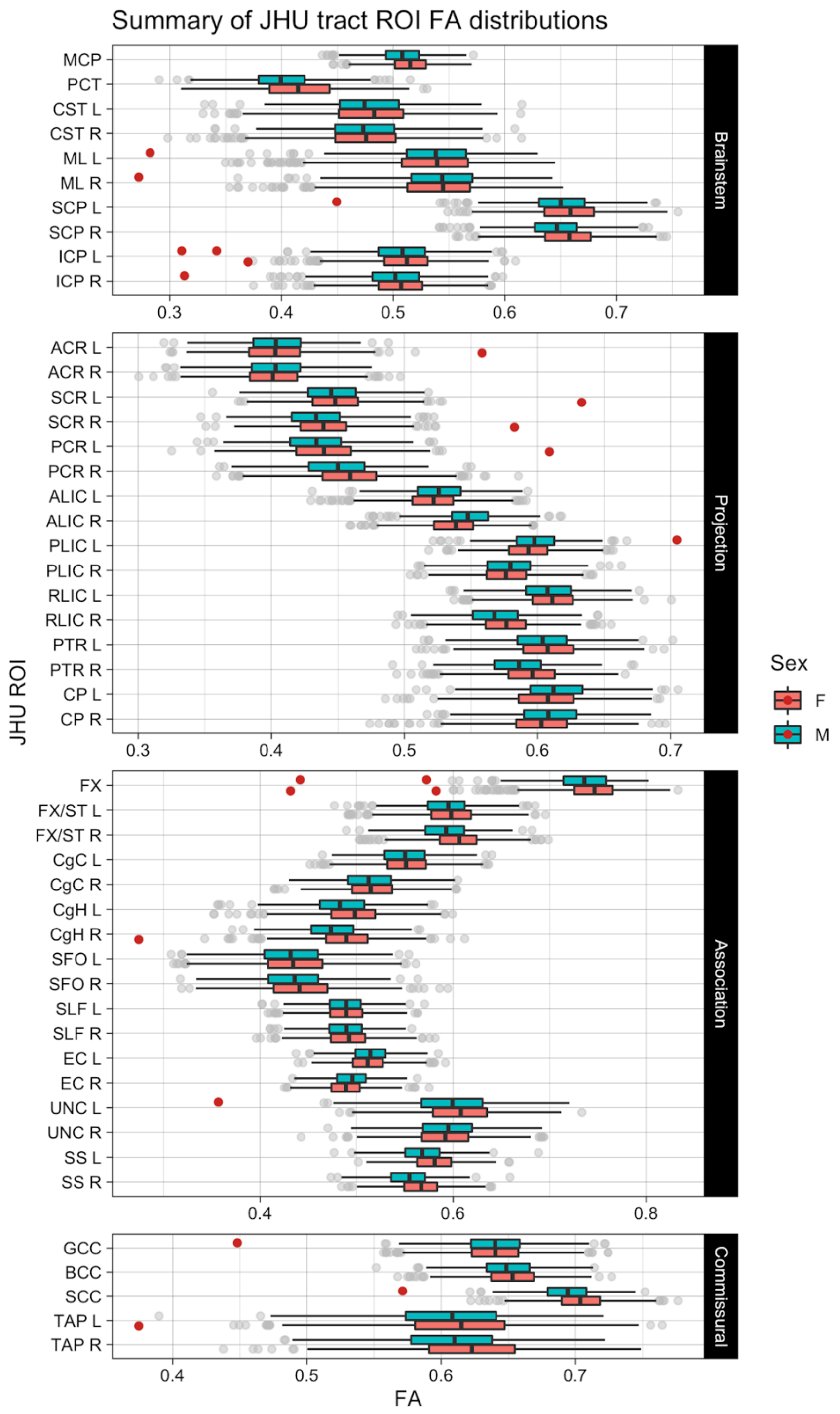

**Supplemental Figure 3. Distribution of WM FA in JHU ROIs.**

Boxplots of regional white matter FA values for each sex in each JHU ROI. See Table 2 of the main text for the full names of the abbreviated ROIs. L: left, R: right. F: female, M: male. Red dots indicate outliers.

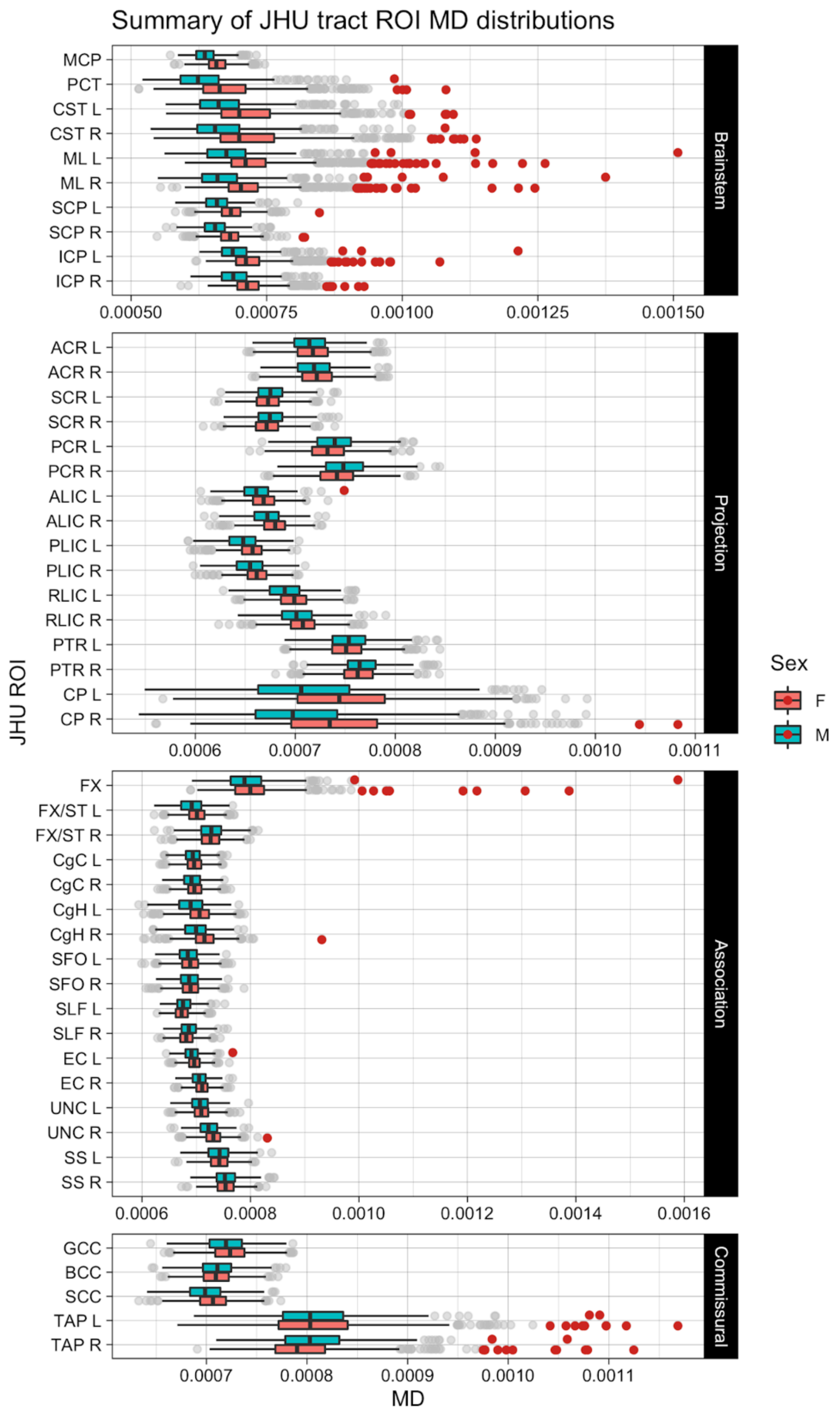

**Supplemental Figure 4. Distribution of WM MD in JHU ROIs.**

Boxplots of regional white matter MD values in $mm^2$/sec for each sex in each JHU ROI. See Table 2 of the main text for the full names of the abbreviated ROIs. L: left, R: right. F: female, M: male. Red dots indicate outliers.

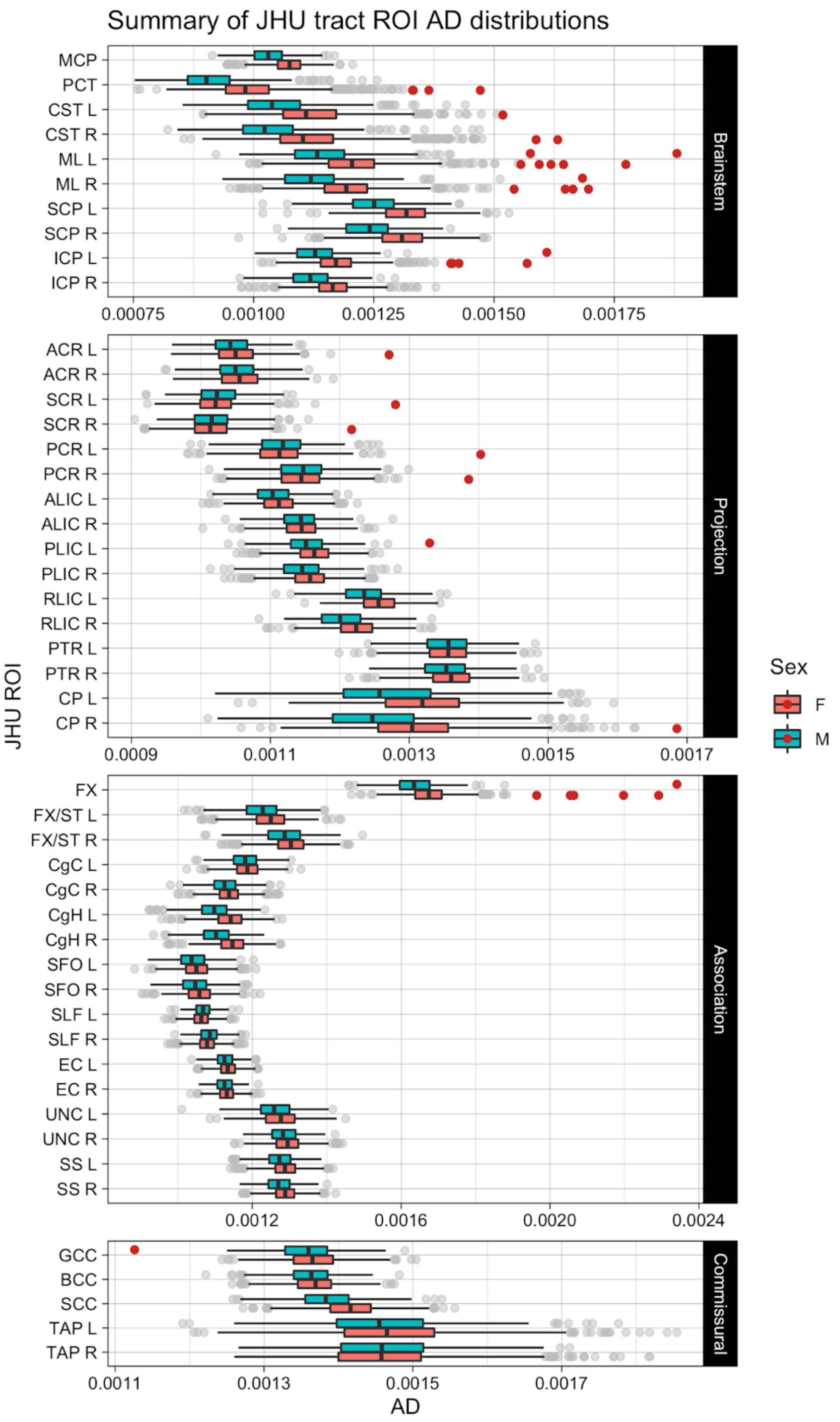

**Supplemental Figure 5. Distribution of WM AD in JHU ROIs.**

Boxplots of regional white matter AD values in $mm^2/sec$ for each sex in each JHU ROI. See Table 2 of the main text for the full names of the abbreviated ROIs. L: left, R: right. F: female, M: male. Red dots indicate outliers.

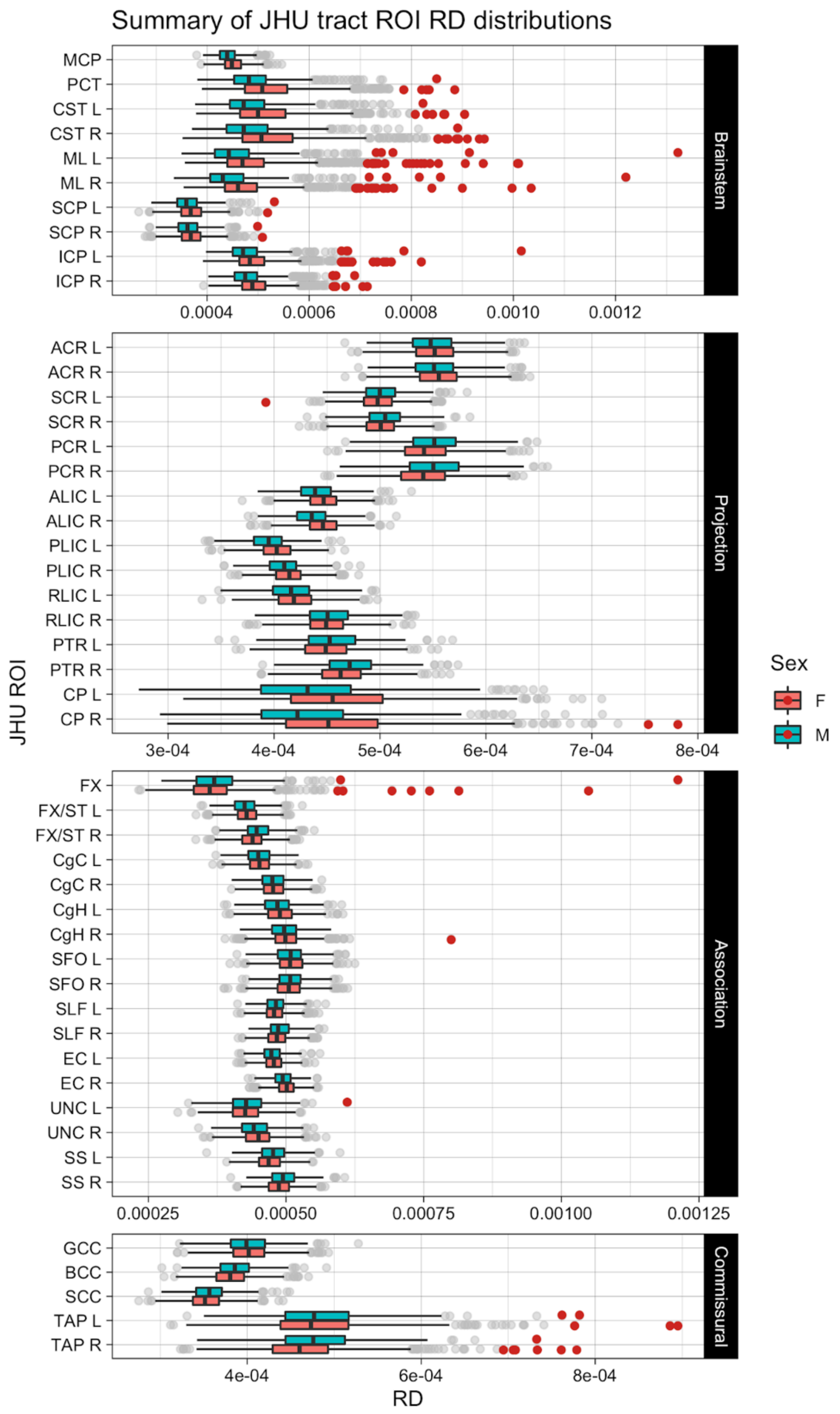

**Supplemental Figure 6. Distribution of WM RD in JHU ROIs.**

Boxplots of regional white matter RD values in $mm^2$/sec for each sex in each JHU ROI. See Table 2 of the main text for the full names of the abbreviated ROIs. L: left, R: right. F: female, M: male. Red dots indicate outliers.

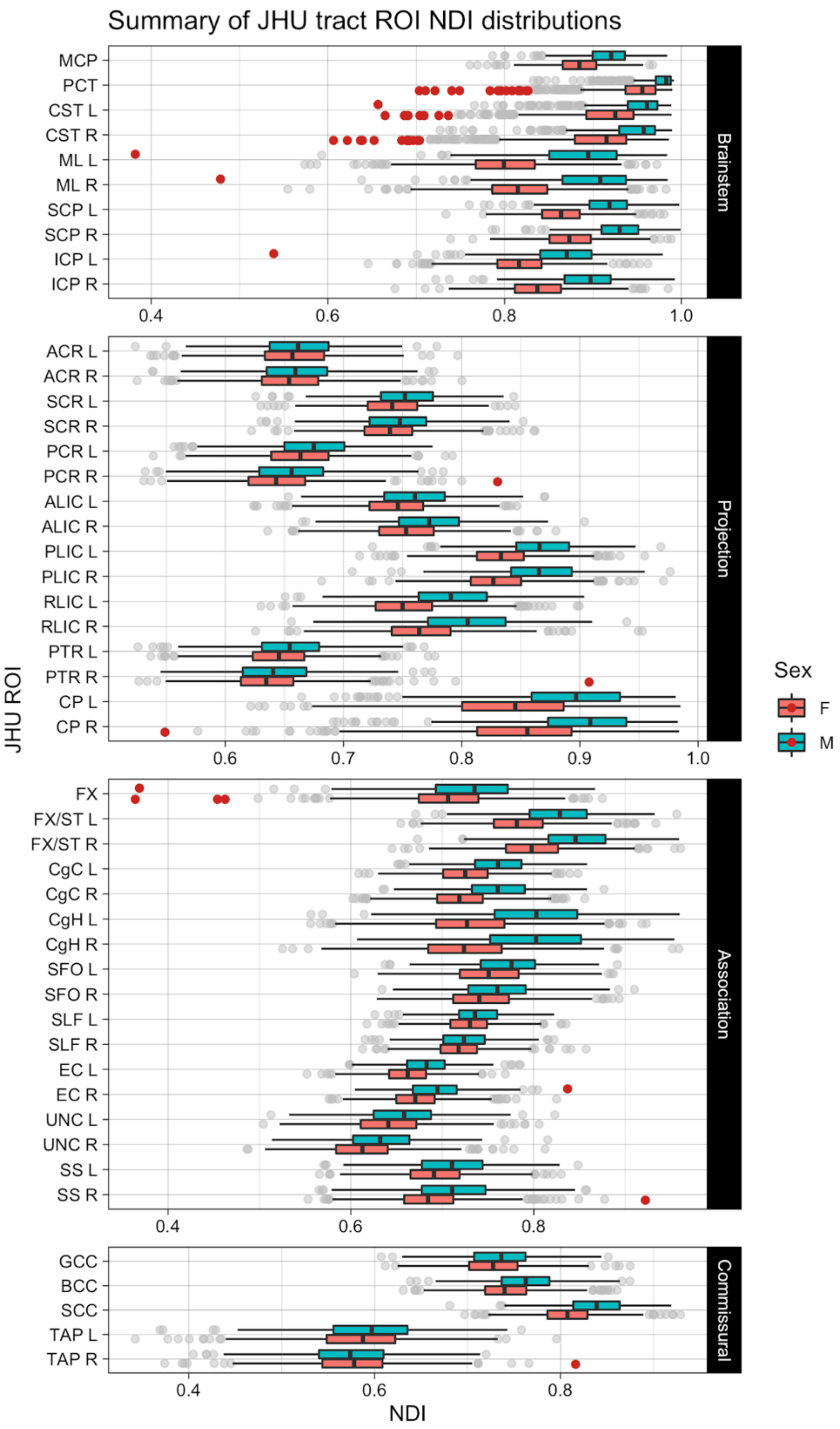

**Supplemental Figure 7. Distribution of WM NDI in JHU ROIs.**

Boxplots of regional white matter NDI values for each sex in each JHU ROI. See Table 2 of the main text for the full names of the abbreviated ROIs. L: left, R: right. F: female, M: male. Red dots indicate outliers.

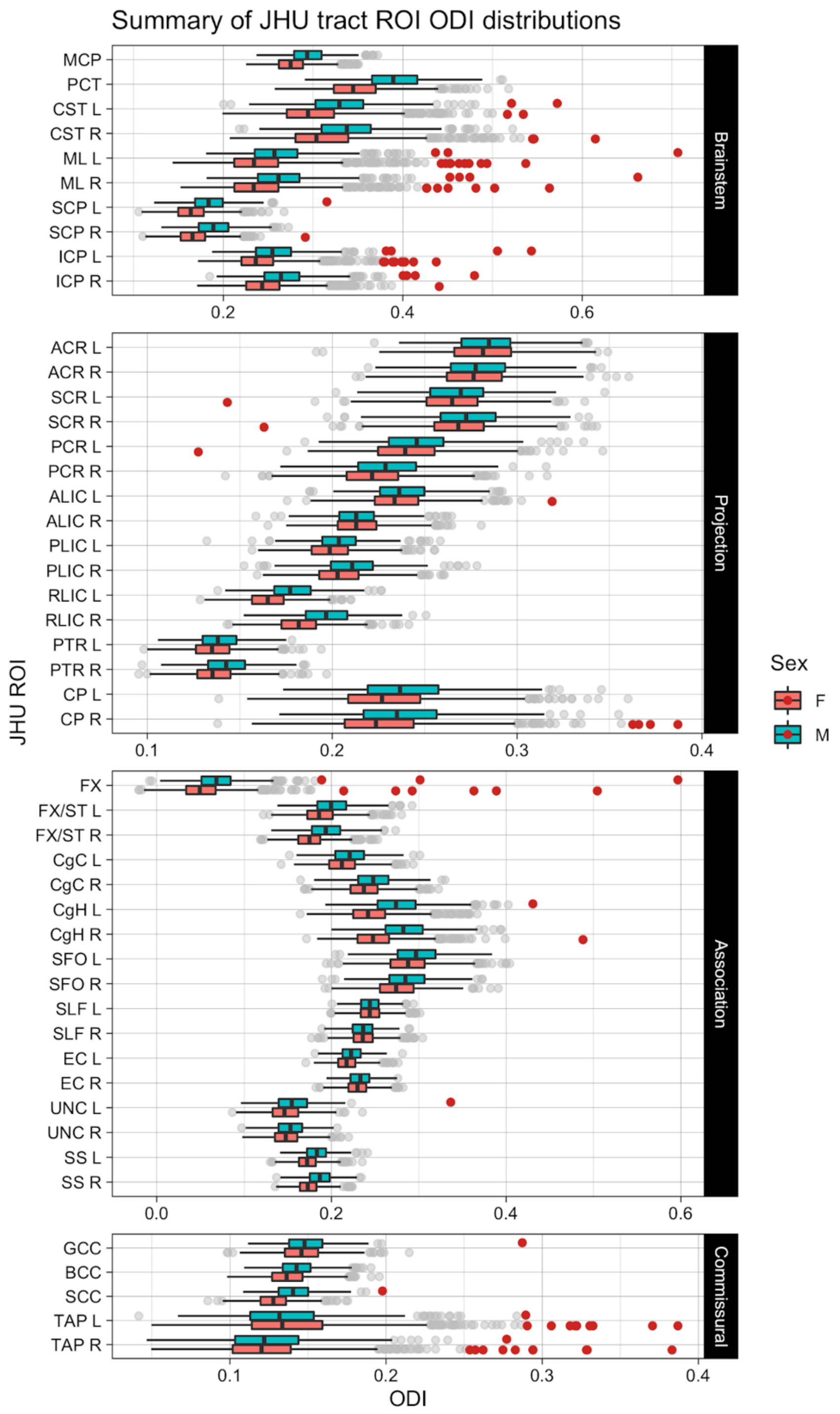

**Supplemental Figure 8. Distribution of WM ODI in JHU ROIs.**

Boxplots of regional white matter ODI values for each sex in each JHU ROI. See Table 2 of the main text for the full names of the abbreviated ROIs. L: left, R: right. F: female, M: male. Red dots indicate outliers.

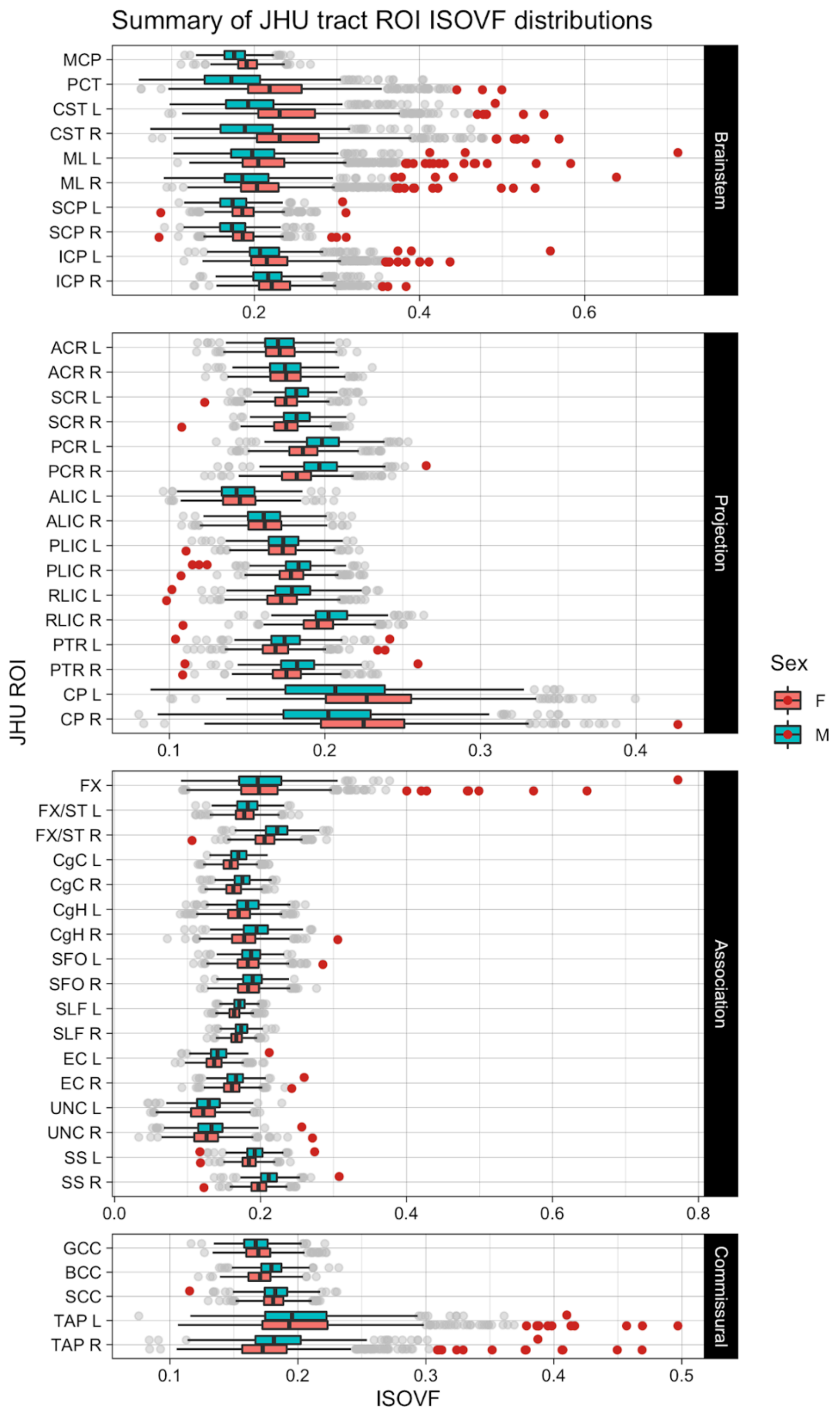

**Supplemental Figure 9. Distribution of WM IsoVF in JHU ROIs.**

Boxplots of regional white matter ISOVF values for each sex in each JHU ROI. See Table 2 of the main text for the full names of the abbreviated ROIs. L: left, R: right. F: female, M: male. Red dots indicate outliers.

### Complete tables of model results for each metric

Due to limited space, we provided tables of raw parameter estimates for selected effects (age, sex, and hemisphere asymmetry) in the main manuscript. Here we summarize parameter estimates ($\beta$), $p$ values, and generalized $\eta^2$ for these main effects as well as for *Age x Sex*, *Age x Hemisphere*, and *Sex x Hemisphere* interaction terms, and also provide adjusted $R^2$ for linear model and marginal $R^2$ for the linear mixed-effects model results.

##### Supplemental Table 2 Model results for WM JHU ROI volumes.

The raw parameter estimate ($\beta$) values and their 95 % confidence intervals (in square brackets) are in $mm^3$/year for Age, Age X Sex and Age X Hemisphere effects, and in $mm^3$ for Sex, Hemisphere and Sex by Hemisphere effects. The first columns give the abbreviated JHU ROIs (see Table 2 of the main text for their full names). Statistical significance symbols (uncorrected for multiple comparisons) *: 0.05 < $p$ <0.001, **: 0.001 < $p$ < 0.0001, ***: $p$ < 0.0001. Bold symbols indicate Bonferroni-corrected significant p-values.

| | Age | | | Sex | | | Age X Sex | | | |
|---|---|---|---|---|---|---|---|---|---|---|
| | *β [95%CI]* | *p value* | $\eta^2_G$ | *β [95%CI]* | *p value* | $\eta^2_G$ | *β [95%CI]* | *p value* | $\eta^2_G$ | *adj. R2/ mar. R2* |
| Brainstem | | | | | | | | | | |
| MCP | -13.6 [ -37.8, 10.7] | >0.1 | <.001 | -70.0* [-125.3, -14.7] | 0.0131 | 0.002 | 7.1 [ -17.1, 31.4] | >0.1 | <.001 | 0.547 |
| PCT | -1.5 [ -4.0, 1.1] | >0.1 | <.001 | **-14.0*** [ -19.8, -8.1]** | **<.0001** | **0.007** | 1.3 [ -1.2, 3.9] | >0.1 | <.001 | 0.577 |
| CST | -0.7 [ -2.8, 1.5] | >0.1 | <.001 | **-10.8*** [ -15.6, -6.0]** | **<.0001** | **0.006** | 0.7 [ -1.4, 2.8] | >0.1 | <.001 | 0.608 |
| ML | -0.7 [ -1.7, 0.4] | >0.1 | <.001 | -2.8* [ -5.2, -0.4] | 0.0224 | 0.002 | 0.6 [ -0.5, 1.7] | >0.1 | <.001 | 0.567 |
| SCP | -1.8* [ -3.2, -0.4] | 0.0142 | 0.002 | **7.9*** [ 4.6, 11.1]** | **<.0001** | **0.009** | -0.5 [ -1.9, 0.9] | >0.1 | <.001 | 0.354 |
| ICP | 0.2 [ -1.1, 1.5] | >0.1 | <.001 | 1.9 [ -1.0, 4.8] | >0.1 | <.001 | 0.1 [ -1.2, 1.4] | >0.1 | <.001 | 0.474 |
| Projection | | | | | | | | | | |
| ACR | 10.2* [ 3.7, 16.6] | 0.002 | 0.001 | 11.0 [ -3.6, 25.7] | >0.1 | <.001 | -1.6 [ -8.0, 4.9] | >0.1 | <.001 | 0.838 |
| SCR | **12.6*** [ 6.7, 18.5]** | **<.0001** | **0.002** | 11.4 [ -2.1, 24.8] | 0.0977 | <.001 | -5.7 [ -11.6, 0.2] | 0.0588 | <.001 | 0.861 |
| PCR | 6.6** [ 3.0, 10.2] | 0.0003 | 0.002 | **-25.5*** [ -33.8, -17.3]** | **<.0001** | **0.006** | -3.4 [ -7.0, 0.2] | 0.0679 | <.001 | 0.811 |
| ALIC | 4.0* [ 1.1, 6.9] | 0.0073 | 0.001 | -6.4 [ -13.0, 0.2] | 0.0567 | <.001 | -2.6 [ -5.5, 0.3] | 0.0817 | <.001 | 0.743 |
| PLIC | 2.8 [ -1.0, 6.6] | >0.1 | <.001 | 3.4 [ -5.3, 12.1] | >0.1 | <.001 | -2.1 [ -5.9, 1.7] | >0.1 | <.001 | 0.773 |
| RLIC | 1.1 [ -0.8, 3.1] | >0.1 | <.001 | -7.9** [ -12.5, -3.4] | 0.0007 | 0.002 | -1.6 [ -3.6, 0.4] | >0.1 | <.001 | 0.835 |
| PTR | 2.1 [ -1.7, 5.9] | >0.1 | <.001 | -4.8 [ -13.4, 3.8] | >0.1 | <.001 | -2.4 [ -6.1, 1.4] | >0.1 | <.001 | 0.79 |

| | | | | | | | | | | |
|---|---|---|---|---|---|---|---|---|---|---|
| CP | -2.1 [ -4.7, 0.5] | >0.1 | <.001 | 1.4 [ -4.6, 7.3] | >0.1 | <.001 | -0.3 [ -2.9, 2.3] | >0.1 | <.001 | 0.704 |
| Association | | | | | | | | | | |
| FX | **2.8*** [ 1.5, 4.0]** | **<.0001** | **0.011** | -3.7* [ -6.4, -0.9] | 0.0094 | 0.004 | -0.2 [ -1.4, 1.0] | >0.1 | <.001 | 0.111 |
| FX/ST | 0.4 [ -0.9, 1.7] | >0.1 | <.001 | -0.7 [ -3.7, 2.3] | >0.1 | <.001 | -1.2 [ -2.5, 0.2] | 0.085 | <.001 | 0.816 |
| CgC | **9.2*** [ 6.3, 12.0]** | **<.0001** | **0.009** | -10.4* [ -16.9, -3.9] | 0.0018 | 0.002 | -2.7 [ -5.6, 0.2] | 0.0639 | <.001 | 0.585 |
| CgH | **6.1*** [ 4.2, 8.0]** | **<.0001** | **0.013** | 2.5 [ -1.9, 6.8] | >0.1 | <.001 | -1.8 [ -3.7, 0.1] | 0.0684 | 0.001 | 0.463 |
| SFO | 0.9** [ 0.4, 1.4] | 0.0006 | 0.002 | 2.0** [ 0.8, 3.2] | 0.0009 | 0.002 | -0.2 [ -0.8, 0.3] | >0.1 | <.001 | 0.754 |
| SLF | **12.4*** [ 6.7, 18.1]** | **<.0001** | **0.002** | **-29.8*** [ -42.8, -16.8]** | **<.0001** | **0.003** | -6.8* [ -12.5, -1.1] | 0.0197 | <.001 | 0.841 |
| EC | 6.3* [ 1.5, 11.1] | 0.0098 | 0.002 | **-48.0*** [ -58.9, -37.1]** | **<.0001** | **0.019** | 2.3 [ -2.4, 7.1] | >0.1 | <.001 | 0.658 |
| UNC | -0.2 [ -1.1, 0.8] | >0.1 | <.001 | **-6.9*** [ -9.0, -4.7]** | **<.0001** | **0.018** | 0.0 [ -0.9, 0.9] | >0.1 | <.001 | 0.191 |
| SS | 1.1 [ -1.0, 3.2] | >0.1 | <.001 | **-11.3*** [ -16.1, -6.5]** | **<.0001** | **0.003** | -1.7 [ -3.8, 0.4] | >0.1 | <.001 | 0.79 |
| Commissural | | | | | | | | | | |
| GCC | 12.5* [ 0.1, 24.9] | 0.0485 | 0.001 | 25.1 [ -3.2, 53.4] | 0.0822 | <.001 | -5.6 [ -18.1, 6.8] | >0.1 | <.001 | 0.621 |
| BCC | 32.7** [ 14.8, 50.7] | 0.0004 | 0.003 | 1.8 [ -39.1, 42.7] | >0.1 | <.001 | -24.0* [ -41.9, -6.0] | 0.0088 | 0.002 | 0.647 |
| SCC | **41.8** [ 20.1, 63.6]** | **0.0002** | **0.005** | -23.1 [ -72.8, 26.6] | >0.1 | <.001 | -5.5 [ -27.2, 16.3] | >0.1 | <.001 | 0.533 |
| TAP | -2.7* [ -4.5, -0.9] | 0.0038 | 0.004 | -4.0 [ -8.1, 0.2] | 0.0611 | 0.001 | -0.3 [ -2.1, 1.5] | >0.1 | <.001 | 0.128 |

| | Hemisphere | | | Age X Hemisphere | | | Sex X Hemisphere | | |
|---|---|---|---|---|---|---|---|---|---|
| | *β [95%CI]* | *p value* | $\eta^2_G$ | *β [95%CI]* | *p value* | $\eta^2_G$ | *β [95%CI]* | *p value* | $\eta^2_G$ |
| Brainstem | | | | | | | | | |
| CST | **-0.8*** [ -1.1, -0.5]** | **<.0001** | **<.001** | 0.0 [ -0.1, 0.2] | >0.1 | <.001 | 0.3* [ 0.0, 0.6] | 0.0491 | <.001 |
| ML | **5.2*** [ 4.9, 5.6]** | **<.0001** | **0.008** | -0.2* [ -0.4, -0.0] | 0.0135 | <.001 | 0.1 [ -0.2, 0.4] | >0.1 | <.001 |
| SCP | -0.1 [ -0.7, 0.4] | >0.1 | <.001 | -0.3 [ -0.6, 0.0] | 0.0902 | <.001 | **1.4*** [ 0.8, 1.9]** | **<.0001** | **<.001** |
| ICP | **9.2*** [ 8.5, 9.9]** | **<.0001** | **0.018** | -0.2 [ -0.6, 0.2] | >0.1 | <.001 | 1.0* [ 0.3, 1.6] | 0.0053 | <.001 |
| Projection | | | | | | | | | |
| ACR | **20.1*** [ 17.1, 23.0]** | **<.0001** | **<.001** | 1.0 [ -0.7, 2.6] | >0.1 | <.001 | **-10.2*** [ -13.1, -7.2]** | **<.0001** | **<.001** |
| SCR | **21.6*** [ 19.0, 24.3]** | **<.0001** | **0.001** | -0.8 [ -2.3, 0.7] | >0.1 | <.001 | -2.5 [ -5.1, 0.1] | 0.0641 | <.001 |
| PCR | -1.1 [ -2.8, 0.6] | >0.1 | <.001 | -0.8 [ -1.8, 0.1] | 0.0715 | <.001 | 2.7* [ 1.0, 4.3] | 0.0015 | <.001 |
| ALIC | **-30.9*** [ -32.6, -29.2]** | **<.0001** | **0.022** | -0.2 [ -1.1, 0.8] | >0.1 | <.001 | **-3.3** [ -5.0, -1.6]** | **0.0001** | **<.001** |

| | | | | | | | | | |
|---|---|---|---|---|---|---|---|---|---|
| PLIC | **26.5***<br>[ 25.2, 27.8]** | **<.0001** | **0.008** | -0.3<br>[ -1.0, 0.4] | >0.1 | <.001 | **-4.6***<br>[ -5.9, -3.3]** | **<.0001** | **<.001** |
| RLIC | **-12.2***<br>[ -13.4, -10.9]** | **<.0001** | **0.005** | 0.2<br>[ -0.5, 0.9] | >0.1 | <.001 | -2.0*<br>[ -3.3, -0.8] | 0.0016 | <.001 |
| PTR | **-30.9***<br>[ -33.2, -28.5]** | **<.0001** | **0.011** | -1.4*<br>[ -2.8, -0.1] | 0.0315 | <.001 | **6.6***<br>[ 4.2, 9.0]** | **<.0001** | **<.001** |
| CP | **13.6***<br>[ 12.7, 14.4]** | **<.0001** | **0.006** | -0.2<br>[ -0.6, 0.3] | >0.1 | <.001 | 0.1<br>[ -0.7, 0.9] | >0.1 | <.001 |
| Association | | | | | | | | | |
| FX/ST | **90.0***<br>[ 89.0, 90.9]** | **<.0001** | **0.597** | -0.1<br>[ -0.6, 0.4] | >0.1 | <.001 | **-4.5***<br>[ -5.4, -3.5]** | **<.0001** | **<.001** |
| CgC | 2.9<br>[ -0.4, 6.1] | 0.0822 | <.001 | 2.3*<br>[ 0.5, 4.1] | 0.0134 | <.001 | -2.8<br>[ -6.0, 0.4] | 0.0896 | <.001 |
| CgH | 2.4**<br>[ 1.0, 3.8] | 0.001 | <.001 | -0.3<br>[ -1.0, 0.5] | >0.1 | <.001 | 0.1<br>[ -1.3, 1.5] | >0.1 | <.001 |
| SFO | **3.4***<br>[ 3.0, 3.8]** | **<.0001** | **0.007** | -0.1<br>[ -0.3, 0.2] | >0.1 | <.001 | **-0.8***<br>[ -1.2, -0.4]** | **<.0001** | **<.001** |
| SLF | **14.0***<br>[ 10.7, 17.3]** | **<.0001** | **<.001** | -2.0*<br>[ -3.9, -0.2] | 0.0291 | <.001 | -1.9<br>[ -5.2, 1.4] | >0.1 | <.001 |
| EC | **105.8***<br>[ 103.1, 108.4]** | **<.0001** | **0.127** | 0.6<br>[ -0.8, 2.1] | >0.1 | <.001 | **-21.7***<br>[ -24.4, -19.1]** | **<.0001** | **0.002** |
| UNC | **-7.2***<br>[ -7.8, -6.6]** | **<.0001** | **0.030** | 0.4*<br>[ 0.0, 0.7] | 0.0239 | <.001 | 0.9*<br>[ 0.3, 1.5] | 0.0019 | <.001 |
| SS | **-30.9***<br>[ -32.4, -29.3]** | **<.0001** | **0.034** | 0.5<br>[ -0.3, 1.4] | >0.1 | <.001 | -1.1<br>[ -2.6, 0.5] | >0.1 | <.001 |
| Commissural | | | | | | | | | |
| TAP | **-23.4***<br>[ -24.4, -22.4]** | **<.0001** | **0.081** | -0.1<br>[ -0.6, 0.5] | >0.1 | <.001 | 0.2<br>[ -0.8, 1.3] | >0.1 | <.001 |

**Supplemental Table 3. Model results for mean FA in JHU ROIs.**

The raw parameter estimate ($\beta$) values and their 95 % confidence intervals (in square brackets) are x$10^{-3}$ differences in FA (/year change in FA for Age, Age X Sex and Age X Hemisphere effects). The first columns give the abbreviated JHU ROIs (see Table 2 of the main text for their full names). Statistical significance symbols (uncorrected for multiple comparisons) *: 0.05 < $p$ <0.001, **: 0.001 < $p$ < 0.0001, ***: $p$ < 0.0001. Bold symbols indicate Bonferroni-corrected significant p-values.

| | Age | | | Sex | | | Age X Sex | | | *adj. R2/ mar. R2* |
|---|---|---|---|---|---|---|---|---|---|---|
| | *β [95%CI]* | *p value* | $\eta^2_G$ | *β [95%CI]* | *p value* | $\eta^2_G$ | *β [95%CI]* | *p value* | $\eta^2_G$ | |
| Brainstem | | | | | | | | | | |
| MCP | -0.0<br>[ -0.7, 0.6] | >0.1 | <.001 | **2.8**<br>[ 1.4, 4.2]** | **0.0001** | **0.009** | -0.2<br>[ -0.8, 0.4] | >0.1 | <.001 | 0.029 |
| PCT | -0.7<br>[ -1.8, 0.3] | >0.1 | 0.001 | 3.3*<br>[ 0.8, 5.7] | 0.0087 | 0.004 | 0.0<br>[ -1.0, 1.1] | >0.1 | <.001 | 0.043 |
| CST | -1.3*<br>[ -2.4, -0.1] | 0.0311 | 0.002 | 0.3<br>[ -2.3, 2.9] | >0.1 | <.001 | 0.8<br>[ -0.4, 1.9] | >0.1 | <.001 | 0.007 |

| | | | | | | | | | | |
|---|---|---|---|---|---|---|---|---|---|---|
| ML | -1.0 [ -2.3, 0.3] | >0.1 | 0.001 | 0.9 [ -2.1, 3.9] | >0.1 | <.001 | 0.4 [ -0.9, 1.7] | >0.1 | <.001 | 0.006 |
| SCP | -0.2 [ -1.1, 0.7] | >0.1 | <.001 | **7.5*** [ 5.5, 9.5]** | **<.0001** | **0.028** | 1.0* [ 0.1, 1.8] | 0.031 | 0.002 | 0.032 |
| ICP | -0.3 [ -1.2, 0.6] | >0.1 | <.001 | 1.9 [ -0.2, 3.9] | 0.076 | 0.002 | 0.1 [ -0.8, 1.0] | >0.1 | <.001 | 0.012 |
| Projection | | | | | | | | | | |
| ACR | 1.4** [ 0.7, 2.2] | 0.0002 | 0.007 | 0.3 [ -1.4, 2.1] | >0.1 | <.001 | 0.0 [ -0.7, 0.8] | >0.1 | <.001 | 0.01 |
| SCR | -0.6 [ -1.4, 0.1] | 0.0971 | 0.001 | 2.2* [ 0.6, 3.9] | 0.009 | 0.003 | 0.0 [ -0.7, 0.8] | >0.1 | <.001 | 0.042 |
| PCR | 0.6 [ -0.3, 1.4] | >0.1 | <.001 | **4.8*** [ 2.8, 6.7]** | **<.0001** | **0.011** | 0.2 [ -0.7, 1.0] | >0.1 | <.001 | 0.088 |
| ALIC | 0.6 [ -0.1, 1.2] | 0.0809 | 0.001 | -2.7** [ -4.2, -1.2] | 0.0003 | 0.006 | 0.1 [ -0.5, 0.8] | >0.1 | <.001 | 0.149 |
| PLIC | -0.7* [ -1.3, -0.1] | 0.0213 | 0.002 | -1.0 [ -2.4, 0.4] | >0.1 | <.001 | 0.3 [ -0.4, 0.9] | >0.1 | <.001 | 0.14 |
| RLIC | 0.8* [ 0.2, 1.5] | 0.0094 | 0.002 | **4.3*** [ 2.9, 5.8]** | **<.0001** | **0.012** | 0.3 [ -0.4, 0.9] | >0.1 | <.001 | 0.371 |
| PTR | 0.3 [ -0.4, 1.1] | >0.1 | <.001 | 1.6 [ -0.1, 3.3] | 0.0654 | 0.002 | -0.2 [ -0.9, 0.6] | >0.1 | <.001 | 0.076 |
| CP | -1.2* [ -2.0, -0.3] | 0.0052 | 0.004 | -1.3 [ -3.1, 0.6] | >0.1 | <.001 | 0.7 [ -0.1, 1.5] | 0.0999 | 0.001 | 0.025 |
| Association | | | | | | | | | | |
| FX | 0.7 [ -0.4, 1.7] | >0.1 | <.001 | **4.8*** [ 2.5, 7.1]** | **0.0001** | **0.010** | 0.3 [ -0.7, 1.4] | >0.1 | <.001 | 0.019 |
| FX/ST | 1.0* [ 0.2, 1.8] | 0.0197 | 0.002 | **7.2*** [ 5.3, 9.0]** | **<.0001** | **0.026** | 0.1 [ -0.7, 0.9] | >0.1 | <.001 | 0.043 |
| CgC | **2.2*** [ 1.3, 3.0]** | **<.0001** | **0.011** | -0.4 [ -2.3, 1.6] | >0.1 | <.001 | -0.2 [ -1.0, 0.7] | >0.1 | <.001 | 0.272 |
| CgH | 1.2* [ 0.3, 2.1] | 0.0099 | 0.003 | **6.9*** [ 4.8, 9.0]** | **<.0001** | **0.019** | 0.4 [ -0.6, 1.3] | >0.1 | <.001 | 0.048 |
| SFO | 0.1 [ -1.0, 1.3] | >0.1 | <.001 | 4.0* [ 1.4, 6.5] | 0.0021 | 0.005 | 0.6 [ -0.5, 1.7] | >0.1 | <.001 | 0.01 |
| SLF | 1.0* [ 0.3, 1.7] | 0.0059 | 0.004 | -0.1 [ -1.7, 1.5] | >0.1 | <.001 | 0.2 [ -0.5, 0.9] | >0.1 | <.001 | 0.009 |
| EC | **1.3*** [ 0.7, 1.9]** | **<.0001** | **0.007** | 2.3* [ 0.9, 3.7] | 0.0012 | 0.004 | 0.0 [ -0.6, 0.7] | >0.1 | <.001 | 0.238 |
| UNC | 0.9 [ -0.1, 2.0] | 0.0783 | 0.001 | 3.4* [ 1.0, 5.8] | 0.0053 | 0.004 | 0.1 [ -0.9, 1.2] | >0.1 | <.001 | 0.03 |
| SS | **1.6*** [ 0.9, 2.3]** | **<.0001** | **0.010** | **5.6*** [ 4.0, 7.2]** | **<.0001** | **0.022** | -0.3 [ -1.0, 0.4] | >0.1 | <.001 | 0.114 |
| Commissural | | | | | | | | | | |
| GCC | **1.7*** [ 0.9, 2.5]** | **<.0001** | **0.010** | 0.5 [ -1.3, 2.4] | >0.1 | <.001 | -0.3 [ -1.1, 0.5] | >0.1 | <.001 | 0.009 |

| | | | | | | | | | | |
|---|---|---|---|---|---|---|---|---|---|---|
| BCC | 0.6 [ -0.1, 1.3] | 0.0721 | 0.002 | -0.1 [ -1.7, 1.5] | >0.1 | <.001 | -0.5 [ -1.2, 0.2] | >0.1 | <.001 | 0.012 |
| SCC | **1.3*** [ 0.7, 1.9]** | **<.0001** | **0.009** | 1.7* [ 0.3, 3.2] | 0.0167 | 0.003 | 0.2 [ -0.4, 0.8] | >0.1 | <.001 | 0.083 |
| TAP | 1.9* [ 0.6, 3.2] | 0.0051 | 0.004 | -0.4 [ -3.4, 2.6] | >0.1 | <.001 | 0.5 [ -0.8, 1.8] | >0.1 | <.001 | 0.04 |
| | Hemisphere | | | Age X Hemisphere | | | Sex X Hemisphere | | | |
| | *β [95%CI]* | *p value* | $\eta^2_G$ | *β [95%CI]* | *p value* | $\eta^2_G$ | *β [95%CI]* | *p value* | $\eta^2_G$ | |
| Brainstem | | | | | | | | | | |
| CST | **2.2*** [ 1.4, 2.9]** | **<.0001** | **0.002** | -0.4 [ -0.8, 0.0] | 0.0574 | <.001 | 0.9* [ 0.2, 1.7] | 0.0108 | <.001 | |
| ML | **-2.5*** [ -3.0, -2.0]** | **<.0001** | **0.002** | -0.2 [ -0.4, 0.1] | >0.1 | <.001 | 0.3 [ -0.2, 0.8] | >0.1 | <.001 | |
| SCP | **1.2*** [ 0.7, 1.6]** | **<.0001** | **0.001** | -0.0 [ -0.3, 0.3] | >0.1 | <.001 | -0.8* [ -1.2, -0.3] | 0.0024 | <.001 | |
| ICP | **2.9*** [ 2.4, 3.4]** | **<.0001** | **0.006** | -0.1 [ -0.4, 0.1] | >0.1 | <.001 | -0.3 [ -0.8, 0.1] | >0.1 | <.001 | |
| Projection | | | | | | | | | | |
| ACR | 0.4 [ -0.1, 0.9] | >0.1 | <.001 | 0.1 [ -0.1, 0.4] | >0.1 | <.001 | 0.2 [ -0.3, 0.7] | >0.1 | <.001 | |
| SCR | **5.4*** [ 5.0, 5.9]** | **<.0001** | **0.030** | 0.2 [ -0.1, 0.4] | >0.1 | <.001 | -0.7* [ -1.1, -0.2] | 0.0022 | <.001 | |
| PCR | **-8.2*** [ -8.8, -7.6]** | **<.0001** | **0.048** | -0.0 [ -0.3, 0.3] | >0.1 | <.001 | -1.0* [ -1.5, -0.4] | 0.0015 | <.001 | |
| ALIC | **-9.7*** [-10.1, -9.2]** | **<.0001** | **0.102** | 0.2 [ -0.0, 0.5] | 0.0538 | <.001 | **1.5*** [ 1.1, 2.0]** | **<.0001** | **0.001** | |
| PLIC | **9.0*** [ 8.6, 9.3]** | **<.0001** | **0.103** | 0.2* [ 0.0, 0.4] | 0.0469 | <.001 | **-0.7** [ -1.1, -0.3]** | **0.0002** | **<.001** | |
| RLIC | **18.5*** [ 17.9, 19.0]** | **<.0001** | **0.283** | -0.1 [ -0.4, 0.2] | >0.1 | <.001 | **-1.4*** [ -1.9, -0.8]** | **<.0001** | **<.001** | |
| PTR | **7.4*** [ 6.9, 7.8]** | **<.0001** | **0.050** | 0.1 [ -0.1, 0.4] | >0.1 | <.001 | **-1.5*** [ -2.0, -1.1]** | **<.0001** | **0.001** | |
| CP | **1.9*** [ 1.2, 2.5]** | **<.0001** | **0.002** | -0.1 [ -0.4, 0.2] | >0.1 | <.001 | -0.1 [ -0.7, 0.5] | >0.1 | <.001 | |
| Association | | | | | | | | | | |
| FX/ST | **-1.7*** [ -2.4, -1.0]** | **<.0001** | **0.002** | -0.1 [ -0.5, 0.3] | >0.1 | <.001 | **-2.7*** [ -3.4, -2.0]** | **<.0001** | **0.004** | |
| CgC | **18.0*** [ 17.5, 18.5]** | **<.0001** | **0.191** | -0.1 [ -0.3, 0.2] | >0.1 | <.001 | -0.3 [ -0.8, 0.1] | >0.1 | <.001 | |
| CgH | **4.2*** [ 3.5, 5.0]** | **<.0001** | **0.011** | -0.4 [ -0.8, 0.0] | 0.0765 | <.001 | -0.6 [ -1.3, 0.2] | >0.1 | <.001 | |
| SFO | **-2.2*** [ -3.0, -1.5]** | **<.0001** | **0.002** | 0.3 [ -0.1, 0.7] | 0.0915 | <.001 | -0.7* [ -1.5, -0.0] | 0.0471 | <.001 | |

| | | | | | | | | | |
|---|---|---|---|---|---|---|---|---|---|
| SLF | -0.4* [ -0.9, -0.0] | 0.0439 | <.001 | -0.0 [ -0.3, 0.2] | >0.1 | <.001 | -0.3 [ -0.8, 0.1] | >0.1 | <.001 |
| EC | **10.5*** [ 10.1, 10.9]** | **<.0001** | **0.135** | -0.0 [ -0.2, 0.2] | >0.1 | <.001 | **0.8*** [ 0.5, 1.2]** | **<.0001** | **<.001** |
| UNC | **4.5*** [ 3.6, 5.4]** | **<.0001** | **0.011** | -0.4 [ -0.9, 0.1] | >0.1 | <.001 | **2.5*** [ 1.6, 3.4]** | **<.0001** | **<.001** |
| SS | **6.8*** [ 6.3, 7.3]** | **<.0001** | **0.047** | -0.1 [ -0.4, 0.1] | >0.1 | <.001 | -0.4 [ -0.9, 0.1] | >0.1 | <.001 |
| Commissural | | | | | | | | | |
| TAP | **-2.6*** [ -3.5, -1.8]** | **<.0001** | **0.002** | 0.2 [ -0.3, 0.7] | >0.1 | <.001 | **-2.0*** [ -2.9, -1.2]** | **<.0001** | **<.001** |

**Supplemental Table 4. Model results for mean MD in JHU ROIs.**

The raw parameter estimate ($\beta$) values and their 95 % confidence intervals (in square brackets) are in x$10^{-6}$ $mm^2$/sec/year for Age, Age X Sex and Age X Hemisphere, and in $mm^2$/sec for Sex, Hemisphere and Sex X Hemisphere effects. The first columns give the abbreviated JHU ROIs (see Table 2 of the main text for their full names). Statistical significance symbols (uncorrected for multiple comparisons) *: 0.05 < $p$ <0.001, **: 0.001 < $p$ < 0.0001, ***: $p$ < 0.0001. Bold symbols indicate Bonferroni-corrected significant p-values.

| | Age | | | Sex | | | Age X Sex | | | *adj. R2/ mar. R2* |
|---|---|---|---|---|---|---|---|---|---|---|
| | *β [95%CI]* | *p value* | $\eta^2_G$ | *β [95%CI]* | *p value* | $\eta^2_G$ | *β [95%CI]* | *p value* | $\eta^2_G$ | |
| Brainstem | | | | | | | | | | |
| MCP | -0.9* [ -1.5, -0.2] | 0.0151 | 0.003 | **10.9*** [ 9.4, 12.5]** | **<.0001** | **0.098** | 0.5 [ -0.2, 1.2] | >0.1 | 0.001 | 0.145 |
| PCT | -1.8 [ -3.8, 0.1] | 0.0665 | 0.002 | **24.6*** [ 20.1, 29.1]** | **<.0001** | **0.063** | 1.7 [ -0.3, 3.6] | >0.1 | 0.001 | 0.094 |
| CST | -2.6* [ -4.7, -0.5] | 0.0147 | 0.003 | **23.4*** [ 18.7, 28.2]** | **<.0001** | **0.045** | 1.2 [ -0.9, 3.3] | >0.1 | <.001 | 0.079 |
| ML | -2.5* [ -4.5, -0.6] | 0.0107 | 0.003 | **15.7*** [ 11.3, 20.1]** | **<.0001** | **0.026** | 2.0* [ 0.0, 3.9] | 0.0483 | 0.002 | 0.087 |
| SCP | **-2.1*** [ -2.8, -1.3]** | **<.0001** | **0.014** | **9.0*** [ 7.3, 10.7]** | **<.0001** | **0.051** | 0.8* [ 0.1, 1.6] | 0.03 | 0.002 | 0.203 |
| ICP | -2.1** [ -3.2, -0.9] | 0.0004 | 0.006 | **11.2*** [ 8.6, 13.8]** | **<.0001** | **0.036** | 0.6 [ -0.6, 1.7] | >0.1 | <.001 | 0.075 |
| Projection | | | | | | | | | | |
| ACR | **-1.9*** [ -2.6, -1.3]** | **<.0001** | **0.018** | **4.9*** [ 3.4, 6.4]** | **<.0001** | **0.022** | 0.3 [ -0.3, 1.0] | >0.1 | <.001 | 0.059 |
| SCR | **-1.2*** [ -1.7, -0.7]** | **<.0001** | **0.013** | 0.5 [ -0.6, 1.6] | >0.1 | <.001 | 0.3 [ -0.2, 0.8] | >0.1 | <.001 | 0.044 |
| PCR | -1.2** [ -2.0, -0.5] | 0.0008 | 0.006 | -1.9* [ -3.5, -0.2] | 0.0261 | 0.003 | 0.6 [ -0.1, 1.3] | 0.0939 | 0.002 | 0.064 |
| ALIC | **-2.0*** [ -2.4, -1.5]** | **<.0001** | **0.034** | **4.2*** [ 3.1, 5.2]** | **<.0001** | **0.028** | 0.3 [ -0.1, 0.8] | >0.1 | <.001 | 0.161 |

| | | | | | | | | | | |
|---|---|---|---|---|---|---|---|---|---|---|
| PLIC | -1.4*** [ -1.9, -1.0] | <.0001 | 0.019 | 4.6*** [ 3.6, 5.6] | <.0001 | 0.039 | 0.5* [ 0.1, 1.0] | 0.0256 | 0.002 | 0.101 |
| RLIC | -1.6*** [ -2.1, -1.0] | <.0001 | 0.015 | 4.5*** [ 3.2, 5.8] | <.0001 | 0.023 | 0.5 [ -0.0, 1.1] | 0.0558 | 0.002 | 0.101 |
| PTR | -1.4*** [ -2.0, -0.7] | <.0001 | 0.009 | 1.7* [ 0.3, 3.2] | 0.0206 | 0.003 | 0.6 [ -0.0, 1.3] | 0.0621 | 0.002 | 0.098 |
| CP | -2.3* [ -4.1, -0.4] | 0.0158 | 0.003 | 13.9*** [ 9.6, 18.1] | <.0001 | 0.021 | 0.2 [ -1.6, 2.1] | >0.1 | <.001 | 0.068 |
| Association | | | | | | | | | | |
| FX | 0.3 [ -1.3, 1.9] | >0.1 | <.001 | 3.5 [ -0.1, 7.1] | 0.0549 | 0.002 | 0.2 [ -1.4, 1.8] | >0.1 | <.001 | 0.002 |
| FX/ST | -2.3*** [ -2.9, -1.6] | <.0001 | 0.017 | 0.8 [ -0.7, 2.3] | >0.1 | <.001 | 0.8* [ 0.1, 1.4] | 0.0214 | 0.002 | 0.28 |
| CgC | -2.0*** [ -2.5, -1.5] | <.0001 | 0.029 | 1.6* [ 0.3, 2.8] | 0.0134 | 0.003 | 0.2 [ -0.3, 0.8] | >0.1 | <.001 | 0.042 |
| CgH | -3.1*** [ -3.8, -2.3] | <.0001 | 0.030 | 7.6*** [ 5.9, 9.3] | <.0001 | 0.036 | 0.8* [ 0.0, 1.5] | 0.0419 | 0.002 | 0.127 |
| SFO | -1.7*** [ -2.3, -1.1] | <.0001 | 0.015 | 2.9*** [ 1.6, 4.3] | <.0001 | 0.008 | 0.2 [ -0.4, 0.8] | >0.1 | <.001 | 0.029 |
| SLF | -1.2*** [ -1.7, -0.7] | <.0001 | 0.012 | -0.3 [ -1.5, 0.8] | >0.1 | <.001 | 0.3 [ -0.2, 0.8] | >0.1 | <.001 | 0.096 |
| EC | -1.7*** [ -2.1, -1.3] | <.0001 | 0.026 | 3.2*** [ 2.2, 4.2] | <.0001 | 0.018 | -0.0 [ -0.5, 0.4] | >0.1 | <.001 | 0.21 |
| UNC | -1.8*** [ -2.3, -1.3] | <.0001 | 0.018 | 1.8* [ 0.6, 3.0] | 0.0026 | 0.004 | 0.1 [ -0.4, 0.6] | >0.1 | <.001 | 0.235 |
| SS | -1.9*** [ -2.6, -1.2] | <.0001 | 0.016 | 2.5* [ 1.0, 4.0] | 0.0012 | 0.005 | 0.7* [ 0.1, 1.4] | 0.0346 | 0.002 | 0.092 |
| Commissural | | | | | | | | | | |
| GCC | -1.6*** [ -2.3, -1.0] | <.0001 | 0.013 | 4.5*** [ 3.1, 6.0] | <.0001 | 0.020 | 0.4 [ -0.2, 1.1] | >0.1 | <.001 | 0.036 |
| BCC | -1.3*** [ -1.9, -0.8] | <.0001 | 0.012 | 0.4 [ -0.9, 1.7] | >0.1 | <.001 | 0.7* [ 0.2, 1.3] | 0.0119 | 0.004 | 0.014 |
| SCC | -1.5*** [ -2.1, -0.9] | <.0001 | 0.013 | 3.7*** [ 2.4, 5.1] | <.0001 | 0.016 | 0.9* [ 0.3, 1.5] | 0.0052 | 0.004 | 0.038 |
| TAP | -1.3 [ -2.6, 0.1] | 0.0661 | 0.002 | 3.7* [ 0.6, 6.8] | 0.0207 | 0.002 | 1.5* [ 0.2, 2.9] | 0.0275 | 0.002 | 0.037 |
| | Hemisphere | | | Age X Hemisphere | | | Sex X Hemisphere | | | |
| | *β [95%CI]* | *p value* | $\eta^2_G$ | *β [95%CI]* | *p value* | $\eta^2_G$ | *β [95%CI]* | *p value* | $\eta^2_G$ | |
| Brainstem | | | | | | | | | | |
| CST | -0.8 [ -2.2, 0.7] | >0.1 | <.001 | 0.0 [ -0.8, 0.8] | >0.1 | <.001 | -2.6** [ -4.1, -1.2] | 0.0005 | <.001 | |
| ML | 6.5*** [ 5.6, 7.4] | <.0001 | 0.005 | 0.1 [ -0.4, 0.6] | >0.1 | <.001 | -0.7 [ -1.6, 0.2] | >0.1 | <.001 | |

| | | | | | | | | | |
|---|---|---|---|---|---|---|---|---|---|
| SCP | **1.5*** [ 1.0, 1.9]** | **<.0001** | **<.001** | 0.1 [ -0.1, 0.4] | >0.1 | <.001 | -0.1 [ -0.6, 0.3] | >0.1 | <.001 |
| ICP | 1.2* [ 0.5, 2.0] | 0.0017 | <.001 | 0.2 [ -0.3, 0.6] | >0.1 | <.001 | 0.0 [ -0.7, 0.8] | >0.1 | <.001 |
| Projection | | | | | | | | | |
| ACR | **-2.0*** [ -2.1, -1.8]** | **<.0001** | **0.005** | -0.0 [ -0.1, 0.1] | >0.1 | <.001 | -0.1 [ -0.2, 0.1] | >0.1 | <.001 |
| SCR | 0.2* [ 0.1, 0.4] | 0.0082 | <.001 | 0.0 [ -0.1, 0.1] | >0.1 | <.001 | 0.3* [ 0.1, 0.4] | 0.0015 | <.001 |
| PCR | **-4.8*** [ -5.0, -4.5]** | **<.0001** | **0.025** | -0.1 [ -0.2, 0.1] | >0.1 | <.001 | 0.4* [ 0.2, 0.6] | 0.001 | <.001 |
| ALIC | **-5.1*** [ -5.4, -4.9]** | **<.0001** | **0.064** | -0.2* [ -0.3, -0.0] | 0.0361 | <.001 | -0.3* [ -0.5, -0.0] | 0.0441 | <.001 |
| PLIC | **-3.0*** [ -3.3, -2.8]** | **<.0001** | **0.028** | -0.0 [ -0.2, 0.1] | >0.1 | <.001 | **0.6*** [ 0.3, 0.8]** | **<.0001** | **0.002** |
| RLIC | **-5.4*** [ -5.7, -5.1]** | **<.0001** | **0.045** | -0.0 [ -0.2, 0.1] | >0.1 | <.001 | **0.9*** [ 0.5, 1.2]** | **<.0001** | **<.001** |
| PTR | **-5.6*** [ -5.9, -5.4]** | **<.0001** | **0.043** | -0.2* [ -0.3, -0.0] | 0.0105 | <.001 | 0.1 [ -0.2, 0.3] | >0.1 | <.001 |
| CP | **2.6*** [ 1.5, 3.8]** | **<.0001** | **<.001** | 0.0 [ -0.6, 0.7] | >0.1 | <.001 | -0.5 [ -1.6, 0.7] | >0.1 | <.001 |
| Association | | | | | | | | | |
| FX/ST | **-15.5*** [-16.0, -15.0]** | **<.0001** | **0.213** | **0.6*** [ 0.3, 0.9]** | **<.0001** | **0.001** | **3.1*** [ 2.6, 3.6]** | **<.0001** | **0.004** |
| CgC | 0.3* [ 0.1, 0.5] | 0.0115 | <.001 | 0.1* [ 0.0, 0.2] | 0.0392 | <.001 | **-0.5*** [ -0.7, -0.3]** | **<.0001** | **<.001** |
| CgH | **-4.7*** [ -5.1, -4.2]** | **<.0001** | **0.019** | 0.2 [ -0.1, 0.4] | >0.1 | <.001 | 0.2 [ -0.3, 0.7] | >0.1 | <.001 |
| SFO | -0.6* [ -1.1, -0.2] | 0.0057 | 0.001 | -0.1 [ -0.4, 0.1] | >0.1 | <.001 | 0.3 [ -0.1, 0.8] | >0.1 | <.001 |
| SLF | **-4.5*** [ -4.7, -4.4]** | **<.0001** | **0.047** | -0.0 [ -0.1, 0.1] | >0.1 | <.001 | 0.3* [ 0.1, 0.5] | 0.0014 | <.001 |
| EC | **-6.8*** [ -7.0, -6.5]** | **<.0001** | **0.117** | 0.1 [ -0.1, 0.2] | >0.1 | <.001 | -0.2 [ -0.5, 0.0] | 0.0638 | <.001 |
| UNC | **-9.5*** [-10.0, -9.0]** | **<.0001** | **0.144** | 0.2 [ -0.0, 0.5] | 0.0988 | <.001 | **-1.3*** [ -1.8, -0.8]** | **<.0001** | **<.001** |
| SS | **-6.2*** [ -6.5, -5.8]** | **<.0001** | **0.045** | 0.0 [ -0.1, 0.2] | >0.1 | <.001 | 0.4* [ 0.0, 0.7] | 0.031 | <.001 |
| Commissural | | | | | | | | | |
| TAP | **3.7*** [ 2.7, 4.7]** | **<.0001** | **0.004** | -0.4 [ -0.9, 0.1] | >0.1 | <.001 | **3.5*** [ 2.5, 4.5]** | **<.0001** | **0.002** |

**Supplemental Table 5. Model results for mean AD in JHU ROIs.**

The raw parameter estimate ($\beta$) values and their 95 % confidence intervals (in square brackets) are in x$10^{-6}$ $mm^2$/sec/year for Age, Age X Sex and Age X Hemisphere, and in $mm^2$/sec for Sex,

Hemisphere and Sex X Hemisphere effects. The first columns give the abbreviated JHU ROIs (see Table 2 of the main text for their full names). Statistical significance symbols (uncorrected for multiple comparisons) *: 0.05 < *p* <0.001, **: 0.001 < *p* < 0.0001, ***: *p* < 0.0001. Bold symbols indicate Bonferroni-corrected significant p-values.

| | Age | | | Sex | | | Age X Sex | | | *adj. R2/ mar. R2* |
|---|---|---|---|---|---|---|---|---|---|---|
| | *β [95%CI]* | *p value* | *η2G* | *β [95%CI]* | *p value* | *η2G* | *β [95%CI]* | *p value* | *η2G* | |
| Brainstem | | | | | | | | | | |
| MCP | -1.5*<br>[ -2.6, -0.3] | 0.0115 | 0.003 | **20.1*****<br>**[ 17.5, 22.7]** | **<.0001** | **0.119** | 0.7<br>[ -0.5, 1.8] | >0.1 | <.001 | 0.209 |
| PCT | -3.3*<br>[ -5.6, -1.1] | 0.004 | 0.004 | **38.2*****<br>**[ 33.0, 43.4]** | **<.0001** | **0.108** | 2.2<br>[ -0.1, 4.5] | 0.0558 | 0.002 | 0.189 |
| CST | **-5.7*****<br>**[ -8.2, -3.2]** | **<.0001** | **0.009** | **35.9*****<br>**[ 30.3, 41.6]** | **<.0001** | **0.072** | 2.7*<br>[ 0.2, 5.3] | 0.0314 | 0.002 | 0.132 |
| ML | **-5.2*****<br>**[ -7.5, -2.9]** | **<.0001** | **0.010** | **29.1*****<br>**[ 23.9, 34.3]** | **<.0001** | **0.060** | 3.3*<br>[ 1.0, 5.6] | 0.0051 | 0.004 | 0.154 |
| SCP | **-4.2*****<br>**[ -5.9, -2.5]** | **<.0001** | **0.011** | **29.2*****<br>**[ 25.2, 33.1]** | **<.0001** | **0.099** | 2.9*<br>[ 1.2, 4.7] | 0.001 | 0.005 | 0.212 |
| ICP | **-3.6*****<br>**[ -5.0, -2.1]** | **<.0001** | **0.011** | **20.8*****<br>**[ 17.5, 24.0]** | **<.0001** | **0.073** | 1.2<br>[ -0.3, 2.6] | >0.1 | 0.001 | 0.153 |
| Projection | | | | | | | | | | |
| ACR | -1.3*<br>[ -2.3, -0.3] | 0.0095 | 0.003 | **8.0*****<br>**[ 5.8, 10.3]** | **<.0001** | **0.024** | 0.4<br>[ -0.6, 1.4] | >0.1 | <.001 | 0.047 |
| SCR | **-2.5*****<br>**[ -3.4, -1.5]** | **<.0001** | **0.013** | 3.3*<br>[ 1.1, 5.4] | 0.0031 | 0.004 | 0.5<br>[ -0.4, 1.5] | >0.1 | <.001 | 0.054 |
| PCR | -1.3*<br>[ -2.4, -0.1] | 0.0344 | 0.002 | 2.7*<br>[ 0.0, 5.4] | 0.0462 | 0.002 | 1.1<br>[ -0.0, 2.3] | 0.0569 | 0.002 | 0.132 |
| ALIC | **-2.8*****<br>**[ -3.7, -1.9]** | **<.0001** | **0.016** | 3.8**<br>[ 1.8, 5.8] | 0.0002 | 0.006 | 0.7<br>[ -0.2, 1.6] | >0.1 | 0.001 | 0.24 |
| PLIC | **-3.3*****<br>**[ -4.3, -2.4]** | **<.0001** | **0.026** | **7.0*****<br>**[ 4.9, 9.0]** | **<.0001** | **0.021** | 1.3*<br>[ 0.4, 2.2] | 0.0055 | 0.004 | 0.065 |
| RLIC | **-1.8****<br>**[ -2.7, -0.9]** | **0.0002** | **0.006** | **14.0*****<br>**[ 11.9, 16.1]** | **<.0001** | **0.066** | 1.3*<br>[ 0.4, 2.2] | 0.0055 | 0.003 | 0.247 |
| PTR | **-2.1****<br>**[ -3.2, -1.0]** | **0.0002** | **0.007** | **5.6*****<br>**[ 3.1, 8.1]** | **<.0001** | **0.010** | 0.8<br>[ -0.3, 1.9] | >0.1 | 0.001 | 0.027 |
| CP | **-5.9*****<br>**[ -8.2, -3.6]** | **<.0001** | **0.013** | **22.4*****<br>**[ 17.1, 27.7]** | **<.0001** | **0.035** | 1.8<br>[ -0.5, 4.1] | >0.1 | 0.001 | 0.102 |
| Association | | | | | | | | | | |
| FX | 1.1<br>[ -0.8, 3.0] | >0.1 | <.001 | **17.0*****<br>**[ 12.7, 21.4]** | **<.0001** | **0.034** | 1.4<br>[ -0.5, 3.3] | >0.1 | 0.001 | 0.072 |
| FX/ST | **-3.1*****<br>**[ -4.6, -1.6]** | **<.0001** | **0.006** | **11.0*****<br>**[ 7.6, 14.4]** | **<.0001** | **0.015** | 1.6*<br>[ 0.1, 3.1] | 0.0381 | 0.002 | 0.231 |
| CgC | -0.6<br>[ -1.7, 0.5] | >0.1 | <.001 | 2.0<br>[ -0.6, 4.5] | >0.1 | <.001 | 0.2<br>[ -0.9, 1.3] | >0.1 | <.001 | 0.292 |
| CgH | **-3.6*****<br>**[ -4.9, -2.3]** | **<.0001** | **0.013** | **20.3*****<br>**[ 17.4, 23.3]** | **<.0001** | **0.081** | 1.7*<br>[ 0.4, 3.0] | 0.0113 | 0.003 | 0.156 |
| SFO | **-2.7*****<br>**[ -3.9, -1.5]** | **<.0001** | **0.010** | **8.9*****<br>**[ 6.2, 11.6]** | **<.0001** | **0.020** | 1.0<br>[ -0.2, 2.1] | >0.1 | 0.001 | 0.039 |
| SLF | -0.7<br>[ -1.5, 0.1] | 0.0712 | 0.002 | -0.9<br>[ -2.7, 0.9] | >0.1 | <.001 | 0.6<br>[ -0.2, 1.4] | >0.1 | 0.001 | 0.092 |

| | | | | | | | | | | |
|---|---|---|---|---|---|---|---|---|---|---|
| EC | -1.3** [ -2.0, -0.5] | 0.001 | 0.005 | 8.8*** [ 7.1, 10.6] | <.0001 | 0.048 | -0.0 [ -0.8, 0.7] | >0.1 | <.001 | 0.073 |
| UNC | -2.2* [ -3.5, -0.8] | 0.0014 | 0.005 | 7.3*** [ 4.3, 10.3] | <.0001 | 0.010 | 0.5 [ -0.8, 1.8] | >0.1 | <.001 | 0.061 |
| SS | -1.1* [ -2.2, -0.1] | 0.0394 | 0.002 | 11.8*** [ 9.4, 14.2] | <.0001 | 0.043 | 0.8 [ -0.2, 1.9] | >0.1 | 0.001 | 0.047 |
| Commissural | | | | | | | | | | |
| GCC | -0.7 [ -1.8, 0.5] | >0.1 | <.001 | 10.0*** [ 7.3, 12.7] | <.0001 | 0.030 | 0.4 [ -0.8, 1.6] | >0.1 | <.001 | 0.032 |
| BCC | -1.6* [ -2.6, -0.6] | 0.0022 | 0.005 | 1.4 [ -0.9, 3.6] | >0.1 | <.001 | 0.8 [ -0.2, 1.8] | >0.1 | 0.001 | 0.013 |
| SCC | -1.1 [ -2.3, 0.2] | 0.0895 | 0.002 | 10.2*** [ 7.3, 13.0] | <.0001 | 0.027 | 2.1* [ 0.8, 3.3] | 0.0011 | 0.006 | 0.121 |
| TAP | 0.6 [ -1.8, 3.0] | >0.1 | <.001 | 5.5* [ 0.1, 11.0] | 0.048 | 0.002 | 3.3* [ 0.9, 5.7] | 0.0066 | 0.004 | 0.009 |
| | Hemisphere | | | Age X Hemisphere | | | Sex X Hemisphere | | | |
| | *β [95%CI]* | *p value* | *η2G* | *β [95%CI]* | *p value* | *η2G* | *β [95%CI]* | *p value* | *η2G* | |
| Brainstem | | | | | | | | | | |
| CST | 3.8*** [ 2.0, 5.5] | <.0001 | <.001 | -0.5 [ -1.5, 0.5] | >0.1 | <.001 | -2.5* [ -4.3, -0.8] | 0.0047 | <.001 | |
| ML | 8.0*** [ 7.0, 9.1] | <.0001 | 0.006 | 0.1 [ -0.5, 0.7] | >0.1 | <.001 | -1.2* [ -2.2, -0.1] | 0.0322 | <.001 | |
| SCP | 5.1*** [ 4.2, 6.0] | <.0001 | 0.003 | 0.3 [ -0.2, 0.7] | >0.1 | <.001 | -1.4* [ -2.3, -0.5] | 0.0019 | <.001 | |
| ICP | 4.9*** [ 3.9, 5.8] | <.0001 | 0.005 | 0.0 [ -0.5, 0.6] | >0.1 | <.001 | -0.7 [ -1.6, 0.3] | >0.1 | <.001 | |
| Projection | | | | | | | | | | |
| ACR | -3.1*** [ -3.7, -2.6] | <.0001 | 0.004 | 0.0 [ -0.3, 0.4] | >0.1 | <.001 | 0.2 [ -0.4, 0.7] | >0.1 | <.001 | |
| SCR | 4.2*** [ 3.6, 4.7] | <.0001 | 0.009 | 0.1 [ -0.2, 0.4] | >0.1 | <.001 | -0.2 [ -0.7, 0.3] | >0.1 | <.001 | |
| PCR | -15.1*** [-15.8, -14.3] | <.0001 | 0.083 | -0.1 [ -0.5, 0.3] | >0.1 | <.001 | -0.5 [ -1.2, 0.3] | >0.1 | <.001 | |
| ALIC | -17.7*** [-18.2, -17.1] | <.0001 | 0.174 | 0.1 [ -0.2, 0.4] | >0.1 | <.001 | 1.5*** [ 0.9, 2.0] | <.0001 | <.001 | |
| PLIC | 3.5*** [ 3.0, 4.1] | <.0001 | 0.006 | 0.2 [ -0.1, 0.5] | >0.1 | <.001 | 0.2 [ -0.3, 0.8] | >0.1 | <.001 | |
| RLIC | 16.2*** [ 15.5, 16.9] | <.0001 | 0.120 | -0.2 [ -0.6, 0.2] | >0.1 | <.001 | 0.2 [ -0.6, 0.9] | >0.1 | <.001 | |
| PTR | -0.8* [ -1.5, -0.0] | 0.0429 | <.001 | -0.2 [ -0.6, 0.2] | >0.1 | <.001 | -2.0*** [ -2.7, -1.2] | <.0001 | <.001 | |
| CP | 6.0*** [ 4.7, 7.4] | <.0001 | 0.003 | -0.0 [ -0.8, 0.7] | >0.1 | <.001 | -0.5 [ -1.8, 0.8] | >0.1 | <.001 | |
| Association | | | | | | | | | | |

| | | | | | | | | | |
|---|---|---|---|---|---|---|---|---|---|
| FX/ST | **-29.7*** [-31.0, -28.4]** | **<.0001** | **0.155** | 0.9* [ 0.1, 1.6] | 0.0183 | <.001 | 1.7* [ 0.4, 3.0] | 0.0125 | <.001 |
| CgC | **25.8*** [ 25.1, 26.5]** | **<.0001** | **0.215** | 0.0 [ -0.4, 0.4] | >0.1 | <.001 | -1.2** [ -1.9, -0.5] | 0.0008 | <.001 |
| CgH | **-2.7*** [ -3.7, -1.7]** | **<.0001** | **0.002** | -0.2 [ -0.7, 0.4] | >0.1 | <.001 | -0.2 [ -1.2, 0.8] | >0.1 | <.001 |
| SFO | **-3.7*** [ -4.5, -2.8]** | **<.0001** | **0.006** | 0.1 [ -0.3, 0.6] | >0.1 | <.001 | -0.0 [ -0.9, 0.8] | >0.1 | <.001 |
| SLF | **-8.3*** [ -8.8, -7.7]** | **<.0001** | **0.054** | -0.1 [ -0.4, 0.2] | >0.1 | <.001 | 0.4 [ -0.1, 0.9] | >0.1 | <.001 |
| EC | **0.9*** [ 0.5, 1.3]** | **<.0001** | **<.001** | 0.0 [ -0.2, 0.3] | >0.1 | <.001 | 0.6* [ 0.1, 1.0] | 0.0109 | <.001 |
| UNC | **-11.0*** [-12.2, -9.8]** | **<.0001** | **0.032** | -0.1 [ -0.8, 0.5] | >0.1 | <.001 | 0.9 [ -0.3, 2.1] | >0.1 | <.001 |
| SS | -0.2 [ -1.1, 0.6] | >0.1 | <.001 | -0.1 [ -0.6, 0.4] | >0.1 | <.001 | 0.1 [ -0.7, 0.9] | >0.1 | <.001 |
| Commissural | | | | | | | | | |
| TAP | 1.6 [ -0.1, 3.4] | 0.0681 | <.001 | -0.4 [ -1.3, 0.6] | >0.1 | <.001 | 2.9* [ 1.1, 4.6] | 0.0012 | <.001 |

**Supplemental Table 6. Model results for mean RD in JHU ROIs.**

The raw parameter estimate (*β*) values and their 95 % confidence intervals (in square brackets) are in x$10^{-6}$ $mm^2$/sec/year for Age, Age X Sex and Age X Hemisphere, and in $mm^2$/sec for Sex, Hemisphere and Sex X Hemisphere effects. The first columns give the abbreviated JHU ROIs (see Table 2 of the main text for their full names). Statistical significance symbols (uncorrected for multiple comparisons) *: 0.05 < $p$ <0.001, **: 0.001 < $p$ < 0.0001, ***: $p$ < 0.0001. Bold symbols indicate Bonferroni-corrected significant p-values.

| | Age | | | Sex | | | Age X Sex | | | adj. R2/ mar. R2 |
|---|---|---|---|---|---|---|---|---|---|---|
| | *β [95%CI]* | *p value* | *η2G* | *β [95%CI]* | *p value* | *η2G* | *β [95%CI]* | *p value* | *η2G* | |
| Brainstem | | | | | | | | | | |
| MCP | -0.5 [ -1.2, 0.1] | >0.1 | 0.001 | **6.3*** [ 4.8, 7.9]** | **<.0001** | **0.036** | 0.5 [ -0.2, 1.1] | >0.1 | <.001 | 0.041 |
| PCT | -1.1 [ -3.1, 0.9] | >0.1 | <.001 | **17.9*** [ 13.4, 22.4]** | **<.0001** | **0.034** | 1.4 [ -0.6, 3.3] | >0.1 | 0.001 | 0.042 |
| CST | -1.1 [ -3.2, 1.0] | >0.1 | <.001 | **17.2*** [ 12.4, 22.0]** | **<.0001** | **0.025** | 0.4 [ -1.7, 2.5] | >0.1 | <.001 | 0.044 |
| ML | -1.2 [ -3.3, 0.9] | >0.1 | <.001 | **9.0** [ 4.3, 13.7]** | **0.0002** | **0.008** | 1.3 [ -0.8, 3.4] | >0.1 | <.001 | 0.04 |
| SCP | -1.0* [ -1.8, -0.2] | 0.0129 | 0.003 | -1.0 [ -2.9, 0.8] | >0.1 | <.001 | -0.2 [ -1.0, 0.6] | >0.1 | <.001 | 0.042 |
| ICP | -1.3* [ -2.6, -0.1] | 0.039 | 0.002 | **6.4*** [ 3.6, 9.3]** | **<.0001** | **0.010** | 0.3 [ -1.0, 1.6] | >0.1 | <.001 | 0.022 |
| Projection | | | | | | | | | | |

| | | | | | | | | | | |
|---|---|---|---|---|---|---|---|---|---|---|
| ACR | **-2.2*** [ -3.0, -1.5]** | **<.0001** | **0.018** | **3.3** [ 1.6, 5.0]** | **0.0002** | **0.007** | 0.3 [ -0.5, 1.0] | >0.1 | <.001 | 0.035 |
| SCR | -0.6 [ -1.2, 0.0] | 0.057 | 0.002 | -0.8 [ -2.2, 0.5] | >0.1 | <.001 | 0.2 [ -0.4, 0.8] | >0.1 | <.001 | 0.017 |
| PCR | -1.2* [ -2.1, -0.4] | 0.0044 | 0.004 | **-4.2*** [ -6.1, -2.2]** | **<.0001** | **0.010** | 0.4 [ -0.5, 1.2] | >0.1 | <.001 | 0.028 |
| ALIC | **-1.6*** [ -2.1, -1.0]** | **<.0001** | **0.016** | **4.3*** [ 3.1, 5.6]** | **<.0001** | **0.024** | 0.1 [ -0.4, 0.7] | >0.1 | <.001 | 0.061 |
| PLIC | -0.5 [ -1.0, 0.1] | 0.0899 | 0.001 | **3.4*** [ 2.2, 4.6]** | **<.0001** | **0.015** | 0.1 [ -0.4, 0.6] | >0.1 | <.001 | 0.118 |
| RLIC | **-1.5*** [ -2.1, -0.8]** | **<.0001** | **0.007** | -0.3 [ -1.8, 1.2] | >0.1 | <.001 | 0.2 [ -0.5, 0.8] | >0.1 | <.001 | 0.305 |
| PTR | -1.0* [ -1.8, -0.2] | 0.0159 | 0.003 | -0.2 [ -2.0, 1.7] | >0.1 | <.001 | 0.5 [ -0.3, 1.3] | >0.1 | <.001 | 0.093 |
| CP | -0.5 [ -2.3, 1.3] | >0.1 | <.001 | **9.6*** [ 5.5, 13.7]** | **<.0001** | **0.011** | -0.5 [ -2.3, 1.3] | >0.1 | <.001 | 0.042 |
| Association | | | | | | | | | | |
| FX | -0.1 [ -1.8, 1.7] | >0.1 | <.001 | -3.2 [ -7.2, 0.7] | >0.1 | 0.002 | -0.4 [ -2.1, 1.3] | >0.1 | <.001 | 0.003 |
| FX/ST | **-1.8*** [ -2.6, -1.1]** | **<.0001** | **0.011** | **-4.3*** [ -6.0, -2.7]** | **<.0001** | **0.012** | 0.4 [ -0.4, 1.1] | >0.1 | <.001 | 0.097 |
| CgC | **-2.7*** [ -3.4, -1.9]** | **<.0001** | **0.023** | 1.4 [ -0.3, 3.1] | >0.1 | 0.001 | 0.2 [ -0.5, 1.0] | >0.1 | <.001 | 0.211 |
| CgH | **-2.8*** [ -3.6, -1.9]** | **<.0001** | **0.019** | 1.2 [ -0.8, 3.2] | >0.1 | <.001 | 0.3 [ -0.5, 1.2] | >0.1 | <.001 | 0.054 |
| SFO | -1.3* [ -2.1, -0.4] | 0.003 | 0.005 | -0.0 [ -1.9, 1.9] | >0.1 | <.001 | -0.1 [ -1.0, 0.7] | >0.1 | <.001 | 0.008 |
| SLF | **-1.5*** [ -2.1, -0.8]** | **<.0001** | **0.011** | -0.1 [ -1.5, 1.4] | >0.1 | <.001 | 0.1 [ -0.5, 0.8] | >0.1 | <.001 | 0.034 |
| EC | **-1.9*** [ -2.5, -1.4]** | **<.0001** | **0.019** | 0.4 [ -0.9, 1.7] | >0.1 | <.001 | -0.0 [ -0.6, 0.5] | >0.1 | <.001 | 0.26 |
| UNC | -1.6** [ -2.5, -0.7] | 0.0005 | 0.005 | -0.9 [ -3.0, 1.2] | >0.1 | <.001 | -0.1 [ -1.0, 0.8] | >0.1 | <.001 | 0.091 |
| SS | **-2.3*** [ -3.1, -1.5]** | **<.0001** | **0.016** | -2.1* [ -3.9, -0.4] | 0.0178 | 0.003 | 0.7 [ -0.1, 1.4] | 0.0962 | 0.001 | 0.127 |
| Commissural | | | | | | | | | | |
| GCC | **-2.1*** [ -2.9, -1.2]** | **<.0001** | **0.014** | 1.8 [ -0.1, 3.7] | 0.0591 | 0.002 | 0.4 [ -0.4, 1.3] | >0.1 | <.001 | 0.016 |
| BCC | -1.2* [ -1.9, -0.5] | 0.0011 | 0.006 | -0.1 [ -1.8, 1.5] | >0.1 | <.001 | 0.7 [ -0.0, 1.4] | 0.062 | 0.002 | 0.022 |
| SCC | **-1.7*** [ -2.3, -1.0]** | **<.0001** | **0.013** | 0.5 [ -1.0, 2.0] | >0.1 | <.001 | 0.2 [ -0.4, 0.9] | >0.1 | <.001 | 0.041 |
| TAP | -2.2* [ -3.8, -0.7] | 0.005 | 0.004 | 2.8 [ -0.8, 6.3] | >0.1 | 0.001 | 0.6 [ -0.9, 2.2] | >0.1 | <.001 | 0.048 |

| | Hemisphere | | | Age X Hemisphere | | | Sex X Hemisphere | | |
|---|---|---|---|---|---|---|---|---|---|
| | *β [95%CI]* | *p value* | *η2G* | *β [95%CI]* | *p value* | *η2G* | *β [95%CI]* | *p value* | *η2G* |
| Brainstem | | | | | | | | | |
| CST | **-3.0*** [ -4.5, -1.6]** | **<.0001** | **0.001** | 0.3 [ -0.6, 1.1] | >0.1 | <.001 | -2.7** [ -4.1, -1.2] | 0.0003 | <.001 |
| ML | **5.8*** [ 4.9, 6.6]** | **<.0001** | **0.004** | 0.1 [ -0.3, 0.6] | >0.1 | <.001 | -0.5 [ -1.3, 0.4] | >0.1 | <.001 |
| SCP | -0.4 [ -0.9, 0.1] | >0.1 | <.001 | 0.1 [ -0.2, 0.3] | >0.1 | <.001 | 0.5 [ -0.0, 1.0] | 0.0656 | <.001 |
| ICP | -0.6 [ -1.4, 0.2] | >0.1 | <.001 | 0.2 [ -0.2, 0.6] | >0.1 | <.001 | 0.4 [ -0.4, 1.1] | >0.1 | <.001 |
| Projection | | | | | | | | | |
| ACR | **-1.4*** [ -1.7, -1.1]** | **<.0001** | **0.002** | -0.1 [ -0.3, 0.1] | >0.1 | <.001 | -0.2 [ -0.5, 0.1] | >0.1 | <.001 |
| SCR | **-1.7*** [ -2.0, -1.5]** | **<.0001** | **0.006** | -0.0 [ -0.2, 0.1] | >0.1 | <.001 | 0.5** [ 0.2, 0.8] | 0.0004 | <.001 |
| PCR | 0.4 [ -0.0, 0.8] | 0.0552 | <.001 | -0.1 [ -0.3, 0.2] | >0.1 | <.001 | **0.8*** [ 0.4, 1.2]** | **0.0001** | **<.001** |
| ALIC | **1.2*** [ 0.8, 1.5]** | **<.0001** | **0.002** | -0.3* [ -0.4, -0.1] | 0.0054 | <.001 | **-1.1*** [ -1.5, -0.8]** | **<.0001** | **<.001** |
| PLIC | **-6.3*** [ -6.6, -6.1]** | **<.0001** | **0.079** | -0.1* [ -0.3, -0.0] | 0.0498 | <.001 | **0.7*** [ 0.5, 1.0]** | **<.0001** | **0.001** |
| RLIC | **-16.2*** [-16.7, -15.7]** | **<.0001** | **0.233** | 0.0 [ -0.2, 0.3] | >0.1 | <.001 | **1.2*** [ 0.8, 1.7]** | **<.0001** | **<.001** |
| PTR | **-8.1*** [ -8.5, -7.7]** | **<.0001** | **0.054** | -0.2 [ -0.4, 0.0] | >0.1 | <.001 | **1.1*** [ 0.7, 1.5]** | **<.0001** | **<.001** |
| CP | 0.9 [ -0.3, 2.1] | >0.1 | <.001 | 0.1 [ -0.6, 0.7] | >0.1 | <.001 | -0.5 [ -1.7, 0.7] | >0.1 | <.001 |
| Association | | | | | | | | | |
| FX/ST | **-8.4*** [ -9.0, -7.8]** | **<.0001** | **0.062** | 0.5* [ 0.1, 0.8] | 0.0041 | <.001 | **3.8*** [ 3.2, 4.4]** | **<.0001** | **0.007** |
| CgC | **-12.5*** [-12.9, -12.2]** | **<.0001** | **0.134** | 0.2 [ -0.0, 0.4] | 0.089 | <.001 | -0.1 [ -0.5, 0.2] | >0.1 | <.001 |
| CgH | **-5.7*** [ -6.3, -5.0]** | **<.0001** | **0.023** | 0.4* [ 0.0, 0.7] | 0.0331 | <.001 | 0.4 [ -0.2, 1.0] | >0.1 | <.001 |
| SFO | 0.9* [ 0.3, 1.5] | 0.0019 | <.001 | -0.3 [ -0.6, 0.0] | 0.0903 | <.001 | 0.5 [ -0.1, 1.1] | 0.0748 | <.001 |
| SLF | **-2.7*** [ -3.0, -2.4]** | **<.0001** | **0.011** | -0.0 [ -0.2, 0.1] | >0.1 | <.001 | 0.2 [ -0.1, 0.5] | >0.1 | <.001 |
| EC | **-10.6*** [-10.9, -10.3]** | **<.0001** | **0.166** | 0.1 [ -0.1, 0.2] | >0.1 | <.001 | **-0.6** [ -0.9, -0.3]** | **0.0002** | **<.001** |

| | | | | | | | | | |
|---|---|---|---|---|---|---|---|---|---|
| UNC | **-8.8*** [ -9.6, -8.0]** | **<.0001** | **0.049** | 0.4 [ -0.0, 0.8] | 0.0672 | <.001 | **-2.4*** [ -3.1, -1.6]** | **<.0001** | **0.001** |
| SS | **-9.2*** [ -9.6, -8.7]** | **<.0001** | **0.070** | 0.1 [ -0.1, 0.4] | >0.1 | <.001 | 0.5* [ 0.1, 0.9] | 0.0205 | <.001 |
| Commissural | | | | | | | | | |
| TAP | **4.8*** [ 3.7, 5.9]** | **<.0001** | **0.005** | -0.4 [ -1.0, 0.2] | >0.1 | <.001 | **3.8*** [ 2.7, 4.9]** | **<.0001** | **0.002** |

**Supplemental Table 7. Model results for mean NDI in JHU ROIs.**

The raw parameter estimate (*β*) values and their 95 % confidence intervals (in square brackets) are x$10^{-3}$ difference in NDI for Sex, Hemisphere and Sex X Hemisphere effects, and /year change in NDI for Age, Age X Sex and Age X Hemisphere effects. The first columns give the abbreviated JHU ROIs (see Table 2 of the main text for their full names). Statistical significance symbols (uncorrected for multiple comparisons) *: 0.05 < *p* <0.001, **: 0.001 < *p* < 0.0001, ***: *p* < 0.0001. Bold symbols indicate Bonferroni-corrected significant p-values.

| | Age | | | Sex | | | Age X Sex | | | *adj. R2/ mar. R2* |
|---|---|---|---|---|---|---|---|---|---|---|
| | *β [95%CI]* | *p value* | *η2G* | *β [95%CI]* | *p value* | *η2G* | *β [95%CI]* | *p value* | *η2G* | |
| Brainstem | | | | | | | | | | |
| MCP | 1.1* [ 0.3, 1.9] | 0.0065 | 0.004 | **-12.9*** [-14.8, -11.1]** | **<.0001** | **0.097** | -0.2 [ -1.0, 0.6] | >0.1 | <.001 | 0.233 |
| PCT | 0.4 [ -0.6, 1.3] | >0.1 | <.001 | **-10.1*** [-12.3, -7.9]** | **<.0001** | **0.046** | -0.1 [ -1.0, 0.9] | >0.1 | <.001 | 0.125 |
| CST | 0.3 [ -1.0, 1.6] | >0.1 | <.001 | **-15.5*** [-18.5, -12.5]** | **<.0001** | **0.051** | -0.1 [ -1.4, 1.2] | >0.1 | <.001 | 0.126 |
| ML | 2.4* [ 1.0, 3.9] | 0.0011 | 0.005 | **-27.3*** [-30.7, -23.9]** | **<.0001** | **0.111** | -1.6* [ -3.1, -0.1] | 0.034 | 0.002 | 0.388 |
| SCP | **2.2*** [ 1.3, 3.1]** | **<.0001** | **0.010** | **-16.1*** [-18.1, -14.1]** | **<.0001** | **0.102** | -0.9* [ -1.8, -0.1] | 0.0365 | 0.002 | 0.414 |
| ICP | **2.5*** [ 1.3, 3.6]** | **<.0001** | **0.008** | **-20.6*** [-23.1, -18.0]** | **<.0001** | **0.107** | -1.2* [ -2.4, -0.1] | 0.035 | 0.002 | 0.308 |
| Projection | | | | | | | | | | |
| ACR | **2.9*** [ 1.8, 3.9]** | **<.0001** | **0.015** | **-5.9*** [ -8.3, -3.4]** | **<.0001** | **0.012** | -0.0 [ -1.1, 1.1] | >0.1 | <.001 | 0.034 |
| SCR | **2.2*** [ 1.3, 3.1]** | **<.0001** | **0.012** | **-5.7*** [ -7.8, -3.6]** | **<.0001** | **0.015** | 0.1 [ -0.9, 1.0] | >0.1 | <.001 | 0.036 |
| PCR | **2.8*** [ 1.7, 3.9]** | **<.0001** | **0.014** | **-5.1*** [ -7.5, -2.6]** | **<.0001** | **0.009** | -0.9 [ -1.9, 0.2] | >0.1 | 0.001 | 0.094 |
| ALIC | **3.2*** [ 2.2, 4.2]** | **<.0001** | **0.020** | -1.9 [ -4.1, 0.4] | 0.0991 | 0.001 | -0.0 [ -1.0, 1.0] | >0.1 | <.001 | 0.132 |
| PLIC | **2.0*** [ 1.1, 2.9]** | **<.0001** | **0.009** | **-9.0*** [-11.1, -6.9]** | **<.0001** | **0.035** | -0.2 [ -1.2, 0.7] | >0.1 | <.001 | 0.244 |
| RLIC | **3.5*** [ 2.4, 4.6]** | **<.0001** | **0.018** | **-14.0*** [-16.5, -11.5]** | **<.0001** | **0.056** | -1.1* [ -2.2, -0.0] | 0.0428 | 0.002 | 0.208 |
| PTR | **2.1*** [ 1.1, 3.1]** | **0.0001** | **0.008** | **-5.9*** [ -8.2, -3.6]** | **<.0001** | **0.013** | -1.2* [ -2.2, -0.2] | 0.0167 | 0.003 | 0.042 |
| CP | 1.7* [ 0.0, 3.3] | 0.0457 | 0.002 | **-11.6*** [-15.3, -7.9]** | **<.0001** | **0.018** | 0.1 [ -1.5, 1.7] | >0.1 | <.001 | 0.184 |

| | | | | | | | | | | |
|---|---|---|---|---|---|---|---|---|---|---|
| Association | | | | | | | | | | |
| FX | 3.7*** [ 2.1, 5.3] | <.0001 | 0.011 | -4.7* [ -8.3, -1.0] | 0.0135 | 0.003 | 0.5 [ -1.1, 2.2] | >0.1 | <.001 | 0.074 |
| FX/ST | 3.8*** [ 2.7, 5.0] | <.0001 | 0.019 | -15.5*** [-18.1, -12.9] | <.0001 | 0.060 | -0.9 [ -2.0, 0.3] | >0.1 | 0.001 | 0.243 |
| CgC | 3.9*** [ 2.8, 5.0] | <.0001 | 0.025 | -11.8*** [-14.3, -9.3] | <.0001 | 0.043 | -0.1 [ -1.2, 1.0] | >0.1 | <.001 | 0.209 |
| CgH | 6.9*** [ 5.2, 8.5] | <.0001 | 0.030 | -22.4*** [-26.1, -18.7] | <.0001 | 0.061 | -2.1* [ -3.7, -0.4] | 0.0129 | 0.003 | 0.277 |
| SFO | 3.5*** [ 2.3, 4.8] | <.0001 | 0.015 | -4.9** [ -7.7, -2.0] | 0.0008 | 0.005 | -0.1 [ -1.4, 1.1] | >0.1 | <.001 | 0.064 |
| SLF | 2.6*** [ 1.7, 3.5] | <.0001 | 0.017 | -4.3*** [ -6.3, -2.3] | <.0001 | 0.009 | -0.2 [ -1.1, 0.7] | >0.1 | <.001 | 0.064 |
| EC | 3.5*** [ 2.6, 4.4] | <.0001 | 0.029 | -3.4** [ -5.4, -1.4] | 0.0008 | 0.005 | -0.5 [ -1.3, 0.4] | >0.1 | <.001 | 0.173 |
| UNC | 4.1*** [ 3.0, 5.3] | <.0001 | 0.021 | -2.3 [ -4.9, 0.3] | 0.089 | 0.001 | -1.0 [ -2.1, 0.2] | 0.094 | 0.001 | 0.151 |
| SS | 3.8*** [ 2.6, 5.0] | <.0001 | 0.019 | -11.0*** [-13.8, -8.3] | <.0001 | 0.030 | -1.5* [ -2.7, -0.3] | 0.0134 | 0.003 | 0.066 |
| Commissural | | | | | | | | | | |
| GCC | 2.4** [ 1.2, 3.6] | 0.0001 | 0.009 | -5.4*** [ -8.1, -2.7] | 0.0001 | 0.009 | -0.1 [ -1.3, 1.1] | >0.1 | <.001 | 0.018 |
| BCC | 2.5*** [ 1.4, 3.5] | <.0001 | 0.012 | -9.0*** [-11.4, -6.6] | <.0001 | 0.030 | -0.3 [ -1.3, 0.8] | >0.1 | <.001 | 0.084 |
| SCC | 2.7*** [ 1.7, 3.7] | <.0001 | 0.015 | -10.0*** [-12.3, -7.7] | <.0001 | 0.040 | -1.0 [ -2.0, 0.0] | 0.0595 | 0.002 | 0.155 |
| TAP | 2.4* [ 0.9, 3.8] | 0.0016 | 0.005 | -9.0*** [-12.3, -5.6] | <.0001 | 0.013 | -1.2 [ -2.6, 0.3] | >0.1 | 0.001 | 0.045 |

| | Hemisphere | | | Age X Hemisphere | | | Sex X Hemisphere | | |
|---|---|---|---|---|---|---|---|---|---|
| | *β [95%CI]* | *p value* | *η2G* | *β [95%CI]* | *p value* | *η2G* | *β [95%CI]* | *p value* | *η2G* |
| Brainstem | | | | | | | | | |
| CST | 4.2*** [ 3.3, 5.1] | <.0001 | 0.005 | 0.1 [ -0.4, 0.6] | >0.1 | <.001 | 2.2*** [ 1.4, 3.1] | <.0001 | 0.001 |
| ML | -7.4*** [ -8.1, -6.7] | <.0001 | 0.012 | 0.1 [ -0.3, 0.4] | >0.1 | <.001 | -0.6 [ -1.3, 0.1] | 0.0939 | <.001 |
| SCP | -6.1*** [ -6.5, -5.6] | <.0001 | 0.022 | 0.1 [ -0.2, 0.3] | >0.1 | <.001 | 0.8** [ 0.3, 1.3] | 0.0008 | <.001 |
| ICP | -11.6*** [-12.2, -11.0] | <.0001 | 0.046 | 0.1 [ -0.3, 0.4] | >0.1 | <.001 | 0.9* [ 0.3, 1.5] | 0.005 | <.001 |
| Projection | | | | | | | | | |
| ACR | 1.1*** [ 0.7, 1.5] | <.0001 | <.001 | -0.0 [ -0.3, 0.2] | >0.1 | <.001 | 0.4* [ 0.0, 0.8] | 0.0347 | <.001 |
| SCR | 1.6*** [ 1.3, 2.0] | <.0001 | 0.002 | -0.1 [ -0.3, 0.1] | >0.1 | <.001 | -0.7** [ -1.0, -0.3] | 0.0005 | <.001 |
| PCR | 9.5*** [ 9.0, 9.9] | <.0001 | 0.044 | -0.2* [ -0.5, -0.0] | 0.0426 | <.001 | 0.1 [ -0.4, 0.5] | >0.1 | <.001 |
| ALIC | -5.5*** [ -6.1, -4.9] | <.0001 | 0.014 | 0.0 [ -0.3, 0.3] | >0.1 | <.001 | 1.1** [ 0.5, 1.7] | 0.0002 | <.001 |

| | | | | | | | | | |
|---|---|---|---|---|---|---|---|---|---|
| PLIC | **1.2*** [ 0.8, 1.7]** | **<.0001** | **<.001** | -0.2 [ -0.4, 0.1] | >0.1 | <.001 | **0.9*** [ 0.5, 1.3]** | **0.0001** | **<.001** |
| RLIC | **-7.0*** [ -7.7, -6.4]** | **<.0001** | **0.021** | -0.5* [ -0.9, -0.1] | 0.0069 | <.001 | -0.4 [ -1.1, 0.2] | >0.1 | <.001 |
| PTR | **5.1*** [ 4.5, 5.6]** | **<.0001** | **0.013** | -0.3* [ -0.6, -0.0] | 0.0333 | <.001 | -0.4 [ -0.9, 0.2] | >0.1 | <.001 |
| CP | **-4.4*** [ -5.3, -3.5]** | **<.0001** | **0.003** | 0.1 [ -0.4, 0.6] | >0.1 | <.001 | 0.6 [ -0.3, 1.5] | >0.1 | <.001 |
| Association | | | | | | | | | |
| FX/ST | **-8.7*** [ -9.5, -7.8]** | **<.0001** | **0.024** | -0.1 [ -0.6, 0.3] | >0.1 | <.001 | 1.2* [ 0.3, 2.0] | 0.0054 | <.001 |
| CgC | **1.7*** [ 1.3, 2.0]** | **<.0001** | **0.002** | -0.2 [ -0.4, 0.1] | >0.1 | <.001 | **1.2*** [ 0.9, 1.6]** | **<.0001** | **<.001** |
| CgH | 0.8 [ -0.3, 1.8] | >0.1 | <.001 | -0.3 [ -0.9, 0.3] | >0.1 | <.001 | 1.8* [ 0.7, 2.9] | 0.0015 | <.001 |
| SFO | **4.8*** [ 3.9, 5.7]** | **<.0001** | **0.009** | -0.2 [ -0.7, 0.3] | >0.1 | <.001 | -0.4 [ -1.3, 0.5] | >0.1 | <.001 |
| SLF | **6.1*** [ 5.7, 6.6]** | **<.0001** | **0.026** | -0.2 [ -0.4, 0.1] | >0.1 | <.001 | -0.7* [ -1.2, -0.3] | 0.0022 | <.001 |
| EC | **-5.0*** [ -5.6, -4.5]** | **<.0001** | **0.016** | -0.1 [ -0.4, 0.2] | >0.1 | <.001 | 0.4 [ -0.2, 0.9] | >0.1 | <.001 |
| UNC | **12.9*** [ 11.9, 13.9]** | **<.0001** | **0.058** | 0.1 [ -0.5, 0.6] | >0.1 | <.001 | 1.3* [ 0.3, 2.3] | 0.0145 | <.001 |
| SS | 0.8 [ -0.1, 1.6] | 0.0743 | <.001 | -0.6* [ -1.1, -0.1] | 0.0167 | <.001 | **2.1*** [ 1.2, 2.9]** | **<.0001** | **<.001** |
| Commissural | | | | | | | | | |
| TAP | **7.0*** [ 5.9, 8.0]** | **<.0001** | **0.012** | 0.7* [ 0.1, 1.3] | 0.0148 | <.001 | **-2.4*** [ -3.4, -1.4]** | **<.0001** | **0.001** |

**Supplemental Table 8. Model results for mean ODI in JHU ROIs.**

The raw parameter estimate ($\beta$) values and their 95 % confidence intervals (in square brackets) are x10$^{-3}$ difference in ODI for Sex, Hemisphere and Sex X Hemisphere effects, and /year change in NDI for Age, Age X Sex and Age X Hemisphere effects. The first columns give the abbreviated JHU ROIs (see Table 2 of the main text for their full names). Statistical significance symbols (uncorrected for multiple comparisons) *: 0.05 < $p$ <0.001, **: 0.001 < $p$ < 0.0001, ***: $p$ < 0.0001. Bold symbols indicate Bonferroni-corrected significant p-values.

| | Age | | | Sex | | | Age X Sex | | | adj. R2/ mar. R2 |
|---|---|---|---|---|---|---|---|---|---|---|
| | *β [95%CI]* | *p value* | $\eta^2_G$ | *β [95%CI]* | *p value* | $\eta^2_G$ | *β [95%CI]* | *p value* | $\eta^2_G$ | |
| Brainstem | | | | | | | | | | |
| MCP | 0.8* [ 0.1, 1.4] | 0.0155 | 0.003 | **-7.8*** [ -9.2, -6.4]** | **<.0001** | **0.065** | -0.3 [ -1.0, 0.3] | >0.1 | <.001 | 0.14 |
| PCT | 1.4* [ 0.3, 2.5] | 0.011 | 0.003 | **-17.7*** [-20.1, -15.2]** | **<.0001** | **0.101** | -0.6 [ -1.7, 0.5] | >0.1 | <.001 | 0.225 |

| | | | | | | | | | | |
|---|---|---|---|---|---|---|---|---|---|---|
| CST | 2.0*<br>[ 0.6, 3.3] | 0.0039 | 0.004 | **-11.9*****<br>**[-14.9, -8.9]** | **<.0001** | **0.030** | -1.6*<br>[ -2.9, -0.2] | 0.02 | 0.003 | 0.081 |
| ML | 1.3<br>[ -0.0, 2.6] | 0.0544 | 0.002 | **-10.7*****<br>**[-13.6, -7.7]** | **<.0001** | **0.026** | -0.4<br>[ -1.7, 0.9] | >0.1 | <.001 | 0.049 |
| SCP | 0.9*<br>[ 0.3, 1.6] | 0.0031 | 0.004 | **-9.2*****<br>**[-10.6, -7.8]** | **<.0001** | **0.077** | -1.1**<br>[ -1.7, -0.5] | 0.0006 | 0.006 | 0.172 |
| ICP | 1.2*<br>[ 0.2, 2.1] | 0.0151 | 0.003 | **-7.6*****<br>**[ -9.7, -5.4]** | **<.0001** | **0.025** | -0.8<br>[ -1.7, 0.2] | >0.1 | 0.001 | 0.076 |
| Projection | | | | | | | | | | |
| ACR | -0.3<br>[ -0.9, 0.3] | >0.1 | <.001 | **-3.4*****<br>**[ -4.8, -2.1]** | **<.0001** | **0.012** | -0.0<br>[ -0.6, 0.6] | >0.1 | <.001 | 0.033 |
| SCR | **1.2*****<br>**[ 0.6, 1.8]** | **0.0001** | **0.008** | **-3.1*****<br>**[ -4.4, -1.8]** | **<.0001** | **0.011** | -0.1<br>[ -0.7, 0.5] | >0.1 | <.001 | 0.03 |
| PCR | 0.4<br>[ -0.2, 1.0] | >0.1 | <.001 | **-4.4*****<br>**[ -5.8, -3.0]** | **<.0001** | **0.017** | -0.5<br>[ -1.1, 0.1] | >0.1 | 0.001 | 0.152 |
| ALIC | 0.7*<br>[ 0.2, 1.1] | 0.0068 | 0.003 | -0.8<br>[ -1.8, 0.3] | >0.1 | <.001 | -0.2<br>[ -0.7, 0.3] | >0.1 | <.001 | 0.301 |
| PLIC | **1.3*****<br>**[ 0.8, 1.7]** | **<.0001** | **0.016** | **-2.5*****<br>**[ -3.5, -1.5]** | **<.0001** | **0.012** | -0.5*<br>[ -0.9, -0.0] | 0.0356 | 0.002 | 0.073 |
| RLIC | 0.5*<br>[ 0.1, 0.9] | 0.0096 | 0.002 | **-7.0*****<br>**[ -7.8, -6.1]** | **<.0001** | **0.087** | -0.5*<br>[ -0.9, -0.2] | 0.0045 | 0.003 | 0.36 |
| PTR | 0.5*<br>[ 0.1, 0.8] | 0.0136 | 0.003 | **-2.9*****<br>**[ -3.7, -2.0]** | **<.0001** | **0.022** | -0.3<br>[ -0.6, 0.1] | >0.1 | <.001 | 0.039 |
| CP | **2.4*****<br>**[ 1.6, 3.2]** | **<.0001** | **0.016** | **-3.8*****<br>**[ -5.7, -2.0]** | **<.0001** | **0.008** | -1.2*<br>[ -2.0, -0.4] | 0.0034 | 0.004 | 0.037 |
| Association | | | | | | | | | | |
| FX | 1.5*<br>[ 0.4, 2.5] | 0.0063 | 0.004 | **-5.3*****<br>**[ -7.7, -2.9]** | **<.0001** | **0.011** | -0.8<br>[ -1.8, 0.3] | >0.1 | 0.001 | 0.065 |
| FX/ST | 0.8*<br>[ 0.2, 1.4] | 0.008 | 0.003 | **-8.1*****<br>**[ -9.4, -6.7]** | **<.0001** | **0.061** | -0.7*<br>[ -1.3, -0.1] | 0.0209 | 0.002 | 0.149 |
| CgC | -0.5<br>[ -1.1, 0.2] | >0.1 | <.001 | **-3.1*****<br>**[ -4.6, -1.6]** | **<.0001** | **0.007** | 0.1<br>[ -0.5, 0.8] | >0.1 | <.001 | 0.263 |
| CgH | **1.6****<br>**[ 0.8, 2.5]** | **0.0001** | **0.006** | **-13.5*****<br>**[-15.4, -11.6]** | **<.0001** | **0.085** | -1.2*<br>[ -2.1, -0.4] | 0.0035 | 0.004 | 0.21 |
| SFO | 0.6<br>[ -0.2, 1.5] | >0.1 | 0.001 | **-5.8*****<br>**[ -7.7, -3.9]** | **<.0001** | **0.018** | -0.6<br>[ -1.4, 0.3] | >0.1 | <.001 | 0.057 |
| SLF | -0.1<br>[ -0.6, 0.3] | >0.1 | <.001 | -0.0<br>[ -1.0, 1.0] | >0.1 | <.001 | -0.2<br>[ -0.6, 0.2] | >0.1 | <.001 | 0.058 |
| EC | 0.2<br>[ -0.2, 0.6] | >0.1 | <.001 | **-4.6*****<br>**[ -5.6, -3.7]** | **<.0001** | **0.040** | -0.1<br>[ -0.5, 0.3] | >0.1 | <.001 | 0.172 |
| UNC | 0.8*<br>[ 0.2, 1.3] | 0.0047 | 0.004 | **-3.4*****<br>**[ -4.7, -2.2]** | **<.0001** | **0.013** | -0.3<br>[ -0.8, 0.3] | >0.1 | <.001 | 0.03 |
| SS | -0.1<br>[ -0.5, 0.3] | >0.1 | <.001 | **-6.5*****<br>**[ -7.5, -5.6]** | **<.0001** | **0.084** | -0.3<br>[ -0.7, 0.1] | >0.1 | <.001 | 0.112 |
| Commissural | | | | | | | | | | |
| GCC | -0.5<br>[ -0.9, 0.0] | 0.0637 | 0.002 | **-3.4*****<br>**[ -4.5, -2.4]** | **<.0001** | **0.022** | 0.2<br>[ -0.3, 0.7] | >0.1 | <.001 | 0.025 |

| | | | | | | | | | | |
|---|---|---|---|---|---|---|---|---|---|---|
| BCC | 0.3 [ -0.1, 0.7] | >0.1 | 0.001 | -1.5* [ -2.5, -0.6] | 0.0015 | 0.006 | 0.1 [ -0.3, 0.5] | >0.1 | <.001 | 0.039 |
| SCC | 0.1 [ -0.3, 0.5] | >0.1 | <.001 | -3.6*** [ -4.4, -2.7] | <.0001 | 0.035 | -0.5* [ -0.9, -0.1] | 0.0135 | 0.003 | 0.21 |
| TAP | -0.3 [ -1.3, 0.7] | >0.1 | <.001 | 2.6* [ 0.4, 4.8] | 0.0195 | 0.003 | -0.5 [ -1.4, 0.5] | >0.1 | <.001 | 0.048 |
| | Hemisphere | | | Age X Hemisphere | | | Sex X Hemisphere | | | |
| | *β [95%CI]* | *p value* | $\eta^2_G$ | *β [95%CI]* | *p value* | $\eta^2_G$ | *β [95%CI]* | *p value* | $\eta^2_G$ | |
| Brainstem | | | | | | | | | | |
| CST | -5.1*** [ -5.9, -4.3] | <.0001 | 0.008 | 0.2 [ -0.3, 0.6] | >0.1 | <.001 | -1.1* [ -1.9, -0.3] | 0.0067 | <.001 | |
| ML | -0.2 [ -0.7, 0.3] | >0.1 | <.001 | 0.0 [ -0.3, 0.3] | >0.1 | <.001 | 0.7* [ 0.2, 1.3] | 0.0057 | <.001 | |
| SCP | -2.0*** [ -2.3, -1.6] | <.0001 | 0.004 | -0.1 [ -0.3, 0.1] | >0.1 | <.001 | 0.9*** [ 0.5, 1.2] | <.0001 | <.001 | |
| ICP | -4.0*** [ -4.5, -3.5] | <.0001 | 0.009 | -0.0 [ -0.3, 0.3] | >0.1 | <.001 | 1.3*** [ 0.8, 1.8] | <.0001 | <.001 | |
| Projection | | | | | | | | | | |
| ACR | 2.4*** [ 2.0, 2.9] | <.0001 | 0.008 | 0.0 [ -0.2, 0.2] | >0.1 | <.001 | 0.0 [ -0.4, 0.4] | >0.1 | <.001 | |
| SCR | -2.1*** [ -2.4, -1.7] | <.0001 | 0.006 | -0.1 [ -0.3, 0.1] | >0.1 | <.001 | 0.3 [ -0.1, 0.7] | 0.0899 | <.001 | |
| PCR | 8.6*** [ 8.2, 9.1] | <.0001 | 0.095 | -0.1 [ -0.3, 0.2] | >0.1 | <.001 | 0.5* [ 0.0, 0.9] | 0.0484 | <.001 | |
| ALIC | 11.4*** [ 11.1, 11.7] | <.0001 | 0.238 | -0.1 [ -0.3, 0.1] | >0.1 | <.001 | -0.8*** [ -1.1, -0.5] | <.0001 | <.001 | |
| PLIC | -3.0*** [ -3.3, -2.7] | <.0001 | 0.021 | -0.1 [ -0.3, 0.0] | >0.1 | <.001 | 0.5* [ 0.2, 0.8] | 0.0011 | <.001 | |
| RLIC | -8.8*** [ -9.1, -8.4] | <.0001 | 0.185 | -0.0 [ -0.2, 0.2] | >0.1 | <.001 | 0.4* [ 0.1, 0.8] | 0.0089 | <.001 | |
| PTR | -1.2*** [ -1.4, -0.9] | <.0001 | 0.005 | -0.1 [ -0.2, 0.1] | >0.1 | <.001 | 1.0*** [ 0.7, 1.3] | <.0001 | 0.002 | |
| CP | 0.9* [ 0.3, 1.5] | 0.002 | <.001 | -0.2 [ -0.5, 0.2] | >0.1 | <.001 | 0.2 [ -0.4, 0.8] | >0.1 | <.001 | |
| Association | | | | | | | | | | |
| FX/ST | 5.0*** [ 4.4, 5.5] | <.0001 | 0.032 | -0.2 [ -0.5, 0.1] | >0.1 | <.001 | 1.1*** [ 0.6, 1.6] | <.0001 | 0.001 | |
| CgC | -13.1*** [-13.5, -12.8] | <.0001 | 0.175 | 0.1 [ -0.1, 0.3] | >0.1 | <.001 | 0.7** [ 0.3, 1.0] | 0.0002 | <.001 | |
| CgH | -3.0*** [ -3.7, -2.4] | <.0001 | 0.007 | 0.2 [ -0.2, 0.5] | >0.1 | <.001 | 0.7* [ 0.0, 1.3] | 0.0411 | <.001 | |
| SFO | 5.4*** [ 4.9, 6.0] | <.0001 | 0.023 | -0.2 [ -0.5, 0.1] | >0.1 | <.001 | 0.4 [ -0.2, 1.0] | >0.1 | <.001 | |
| SLF | 4.0*** [ 3.7, 4.4] | <.0001 | 0.038 | -0.0 [ -0.2, 0.1] | >0.1 | <.001 | -0.0 [ -0.3, 0.3] | >0.1 | <.001 | |

| | | | | | | | | | |
|---|---|---|---|---|---|---|---|---|---|
| EC | **-5.5*** [ -5.8, -5.3]** | **<.0001** | **0.085** | -0.0 [ -0.2, 0.1] | >0.1 | <.001 | **-0.5*** [ -0.8, -0.3]** | **0.0001** | **<.001** |
| UNC | 0.7* [ 0.2, 1.2] | 0.0033 | <.001 | 0.2 [ -0.1, 0.5] | >0.1 | <.001 | **-1.1*** [ -1.5, -0.6]** | **<.0001** | **<.001** |
| SS | **-1.0*** [ -1.4, -0.6]** | **<.0001** | **0.003** | -0.0 [ -0.2, 0.2] | >0.1 | <.001 | 0.5* [ 0.1, 0.9] | 0.0061 | <.001 |
| Commissural | | | | | | | | | |
| TAP | **6.8*** [ 6.0, 7.6]** | **<.0001** | **0.024** | -0.1 [ -0.6, 0.4] | >0.1 | <.001 | 1.3* [ 0.5, 2.2] | 0.0015 | <.001 |

**Supplemental Table 9. Model results for mean IsoVF in JHU ROIs.**

The raw parameter estimate ($\beta$) values and their 95 % confidence intervals (in square brackets) are x$10^{-3}$ difference in IsoVF for Sex, Hemisphere and Sex X Hemisphere effects, and /year change in NDI for Age, Age X Sex and Age X Hemisphere effects. The first columns give the abbreviated JHU ROIs (see Table 2 of the main text for their full names). Statistical significance symbols (uncorrected for multiple comparisons) *: 0.05 < $p$ <0.001, **: 0.001 < $p$ < 0.0001, ***: $p$ < 0.0001. Bold symbols indicate Bonferroni-corrected significant p-values.

| | Age | | | Sex | | | Age X Sex | | | adj. R2/ mar. R2 |
|---|---|---|---|---|---|---|---|---|---|---|
| | *β [95%CI]* | *p value* | $\eta^2_G$ | *β [95%CI]* | *p value* | $\eta^2_G$ | *β [95%CI]* | *p value* | $\eta^2_G$ | |
| Brainstem | | | | | | | | | | |
| MCP | -0.5* [ -1.1, -0.0] | 0.0422 | 0.002 | **8.4*** [ 7.2, 9.6]** | **<.0001** | **0.099** | 0.6* [ 0.1, 1.1] | 0.0205 | 0.003 | 0.131 |
| PCT | -1.9* [ -3.5, -0.3] | 0.0233 | 0.003 | **25.9*** [ 22.2, 29.6]** | **<.0001** | **0.098** | 1.8* [ 0.2, 3.4] | 0.0306 | 0.002 | 0.151 |
| CST | -2.0* [ -3.6, -0.4] | 0.0165 | 0.003 | **22.6*** [ 18.9, 26.2]** | **<.0001** | **0.068** | 1.3 [ -0.3, 2.9] | >0.1 | 0.001 | 0.116 |
| ML | -1.5* [ -2.9, -0.1] | 0.0422 | 0.002 | **7.3*** [ 4.1, 10.5]** | **<.0001** | **0.011** | 1.3 [ -0.1, 2.7] | 0.0724 | 0.002 | 0.026 |
| SCP | **-1.3*** [ -1.9, -0.7]** | **<.0001** | **0.008** | **3.8*** [ 2.5, 5.2]** | **<.0001** | **0.014** | 0.6 [ -0.0, 1.2] | 0.0534 | 0.002 | 0.084 |
| ICP | -1.2* [ -2.2, -0.2] | 0.0182 | 0.003 | 4.0** [ 1.8, 6.2] | 0.0004 | 0.006 | 0.3 [ -0.7, 1.2] | >0.1 | <.001 | 0.015 |
| Projection | | | | | | | | | | |
| ACR | -0.0 [ -0.4, 0.4] | >0.1 | <.001 | 1.4* [ 0.5, 2.3] | 0.0028 | 0.005 | 0.3 [ -0.1, 0.7] | 0.0941 | 0.001 | 0.026 |
| SCR | -0.0 [ -0.4, 0.3] | >0.1 | <.001 | **-2.2*** [ -2.9, -1.5]** | **<.0001** | **0.017** | 0.4* [ 0.1, 0.7] | 0.0128 | 0.003 | 0.087 |
| PCR | 0.5* [ 0.1, 1.0] | 0.0127 | 0.003 | **-4.6*** [ -5.6, -3.7]** | **<.0001** | **0.043** | 0.1 [ -0.4, 0.5] | >0.1 | <.001 | 0.174 |
| ALIC | -0.3 [ -0.7, 0.1] | >0.1 | <.001 | **3.9*** [ 3.0, 4.8]** | **<.0001** | **0.025** | 0.3 [ -0.1, 0.7] | 0.088 | 0.001 | 0.273 |
| PLIC | -0.6* [ -0.9, -0.2] | 0.0013 | 0.004 | **2.2*** [ 1.4, 3.0]** | **<.0001** | **0.012** | 0.4* [ 0.1, 0.8] | 0.0121 | 0.003 | 0.146 |

| | | | | | | | | | | |
|---|---|---|---|---|---|---|---|---|---|---|
| RLIC | -0.1 [ -0.5, 0.3] | >0.1 | <.001 | -1.5** [ -2.4, -0.6] | 0.0009 | 0.003 | 0.1 [ -0.3, 0.5] | >0.1 | <.001 | 0.411 |
| PTR | 0.1 [ -0.3, 0.5] | >0.1 | <.001 | **-1.7*** [ -2.6, -0.9]** | **0.0001** | **0.007** | -0.1 [ -0.5, 0.3] | >0.1 | <.001 | 0.123 |
| CP | -1.6* [ -2.8, -0.4] | 0.0096 | 0.003 | **9.5*** [ 6.8, 12.3]** | **<.0001** | **0.024** | 0.3 [ -0.9, 1.5] | >0.1 | <.001 | 0.055 |
| Association | | | | | | | | | | |
| FX | 2.1* [ 0.7, 3.5] | 0.0041 | 0.005 | 2.8 [ -0.5, 6.0] | 0.0936 | 0.002 | 0.1 [ -1.3, 1.5] | >0.1 | <.001 | 0.008 |
| FX/ST | -0.7* [ -1.2, -0.2] | 0.0107 | 0.002 | **-4.3*** [ -5.5, -3.1]** | **<.0001** | **0.015** | 0.4 [ -0.1, 0.9] | 0.0981 | <.001 | 0.407 |
| CgC | -0.3 [ -0.7, 0.1] | >0.1 | <.001 | **-3.0*** [ -4.0, -2.1]** | **<.0001** | **0.019** | 0.3 [ -0.1, 0.7] | >0.1 | <.001 | 0.161 |
| CgH | -0.1 [ -0.7, 0.5] | >0.1 | <.001 | -2.0* [ -3.4, -0.6] | 0.0046 | 0.004 | 0.1 [ -0.5, 0.7] | >0.1 | <.001 | 0.141 |
| SFO | -0.0 [ -0.5, 0.5] | >0.1 | <.001 | 0.5 [ -0.7, 1.6] | >0.1 | <.001 | 0.3 [ -0.2, 0.8] | >0.1 | <.001 | 0.033 |
| SLF | 0.1 [ -0.2, 0.4] | >0.1 | <.001 | **-2.2*** [ -2.8, -1.5]** | **<.0001** | **0.019** | 0.2 [ -0.1, 0.5] | >0.1 | <.001 | 0.107 |
| EC | 0.3 [ -0.1, 0.7] | >0.1 | <.001 | **1.8** [ 0.8, 2.7]** | **0.0002** | **0.004** | -0.2 [ -0.6, 0.2] | >0.1 | <.001 | 0.435 |
| UNC | 0.8* [ 0.1, 1.4] | 0.0155 | 0.002 | 0.3 [ -1.1, 1.7] | >0.1 | <.001 | -0.4 [ -1.0, 0.2] | >0.1 | <.001 | 0.04 |
| SS | 0.1 [ -0.3, 0.5] | >0.1 | <.001 | **-3.2*** [ -4.0, -2.3]** | **<.0001** | **0.017** | -0.1 [ -0.4, 0.3] | >0.1 | <.001 | 0.286 |
| Commissural | | | | | | | | | | |
| GCC | -0.3 [ -0.7, 0.2] | >0.1 | <.001 | **2.7*** [ 1.7, 3.6]** | **<.0001** | **0.017** | 0.4 [ -0.1, 0.8] | 0.0937 | 0.002 | 0.024 |
| BCC | -0.1 [ -0.4, 0.3] | >0.1 | <.001 | **-2.7*** [ -3.5, -1.8]** | **<.0001** | **0.022** | 0.6* [ 0.2, 1.0] | 0.0015 | 0.006 | 0.132 |
| SCC | -0.4* [ -0.8, -0.1] | 0.0196 | 0.003 | 1.1* [ 0.3, 1.9] | 0.0095 | 0.004 | 0.5* [ 0.1, 0.8] | 0.013 | 0.003 | 0.038 |
| TAP | 0.6 [ -0.5, 1.6] | >0.1 | <.001 | 0.5 [ -1.9, 2.9] | >0.1 | <.001 | 0.4 [ -0.7, 1.4] | >0.1 | <.001 | 0.083 |
| | Hemisphere | | | Age X Hemisphere | | | Sex X Hemisphere | | | |
| | *β [95%CI]* | *p value* | $\eta^2_G$ | *β [95%CI]* | *p value* | $\eta^2_G$ | *β [95%CI]* | *p value* | $\eta^2_G$ | |
| Brainstem | | | | | | | | | | |
| CST | 0.4 [ -0.8, 1.5] | >0.1 | <.001 | 0.0 [ -0.6, 0.6] | >0.1 | <.001 | -1.9** [ -3.1, -0.8] | 0.0008 | <.001 | |
| ML | **3.4*** [ 2.7, 4.1]** | **<.0001** | **0.002** | 0.1 [ -0.2, 0.5] | >0.1 | <.001 | **-1.3** [ -2.0, -0.6]** | **0.0002** | **<.001** | |
| SCP | 0.3 [ -0.2, 0.7] | >0.1 | <.001 | 0.1 [ -0.1, 0.4] | >0.1 | <.001 | -0.1 [ -0.5, 0.3] | >0.1 | <.001 | |
| ICP | **-1.9*** [ -2.6, -1.3]** | **<.0001** | **0.003** | 0.2 [ -0.1, 0.6] | >0.1 | <.001 | -0.1 [ -0.8, 0.5] | >0.1 | <.001 | |

| | | | | | | | | | |
|---|---|---|---|---|---|---|---|---|---|
| Projection | | | | | | | | | |
| ACR | **-1.9***<br>[ -2.1, -1.7]** | **<.0001** | **0.011** | -0.1<br>[ -0.2, 0.0] | >0.1 | <.001 | 0.1<br>[ -0.1, 0.3] | >0.1 | <.001 |
| SCR | 0.0<br>[ -0.2, 0.3] | >0.1 | <.001 | -0.0<br>[ -0.1, 0.1] | >0.1 | <.001 | -0.0<br>[ -0.3, 0.2] | >0.1 | <.001 |
| PCR | **1.4***<br>[ 1.1, 1.6]** | **<.0001** | **0.006** | **-0.3**<br>[ -0.4, -0.1]** | **0.0002** | **<.001** | **0.7***<br>[ 0.4, 0.9]** | **<.0001** | **<.001** |
| ALIC | **-8.3***<br>[ -8.6, -7.9]** | **<.0001** | **0.161** | -0.1<br>[ -0.3, 0.1] | >0.1 | <.001 | 0.2<br>[ -0.1, 0.5] | >0.1 | <.001 |
| PLIC | **-3.8***<br>[ -4.1, -3.6]** | **<.0001** | **0.061** | -0.1<br>[ -0.3, 0.0] | >0.1 | <.001 | **0.9***<br>[ 0.6, 1.2]** | **<.0001** | **0.005** |
| RLIC | **-12.0***<br>[-12.4, -11.6]** | **<.0001** | **0.300** | -0.1<br>[ -0.3, 0.1] | >0.1 | <.001 | 0.4<br>[ -0.0, 0.7] | 0.0545 | <.001 |
| PTR | **-3.9***<br>[ -4.2, -3.6]** | **<.0001** | **0.056** | **-0.4***<br>[ -0.6, -0.2]** | **<.0001** | **0.002** | 0.2<br>[ -0.1, 0.5] | >0.1 | 0.001 |
| CP | **1.6***<br>[ 0.8, 2.3]** | **0.0001** | **<.001** | 0.2<br>[ -0.3, 0.6] | >0.1 | <.001 | -0.3<br>[ -1.1, 0.4] | >0.1 | <.001 |
| Association | | | | | | | | | |
| FX/ST | **-17.1***<br>[-17.6, -16.5]** | **<.0001** | **0.317** | **0.6**<br>[ 0.3, 0.9]** | **0.0002** | **0.001** | **3.1***<br>[ 2.6, 3.6]** | **<.0001** | **0.004** |
| CgC | **-2.3***<br>[ -2.5, -2.0]** | **<.0001** | **0.014** | 0.0<br>[ -0.1, 0.2] | >0.1 | <.001 | 0.3*<br>[ 0.1, 0.6] | 0.008 | <.001 |
| CgH | **-4.6***<br>[ -5.1, -4.1]** | **<.0001** | **0.026** | 0.1<br>[ -0.2, 0.4] | >0.1 | <.001 | **1.0**<br>[ 0.5, 1.5]** | **0.0002** | **<.001** |
| SFO | -0.6<br>[ -1.2, 0.1] | 0.0761 | <.001 | -0.3<br>[ -0.6, 0.1] | >0.1 | <.001 | 0.4<br>[ -0.2, 1.0] | >0.1 | <.001 |
| SLF | **-1.4***<br>[ -1.7, -1.2]** | **<.0001** | **0.013** | -0.2*<br>[ -0.3, -0.0] | 0.0218 | <.001 | -0.1<br>[ -0.3, 0.2] | >0.1 | <.001 |
| EC | **-12.0***<br>[-12.3, -11.6]** | **<.0001** | **0.285** | 0.0<br>[ -0.2, 0.2] | >0.1 | <.001 | -0.2<br>[ -0.5, 0.2] | >0.1 | <.001 |
| UNC | **-2.0***<br>[ -2.7, -1.3]** | **<.0001** | **0.005** | 0.2<br>[ -0.2, 0.6] | >0.1 | <.001 | -0.3<br>[ -1.0, 0.4] | >0.1 | <.001 |
| SS | **-8.4***<br>[ -8.8, -8.0]** | **<.0001** | **0.164** | -0.2<br>[ -0.4, 0.0] | 0.0685 | <.001 | **1.5***<br>[ 1.1, 1.9]** | **<.0001** | **0.001** |
| Commissural | | | | | | | | | |
| TAP | **10.2***<br>[ 9.3, 11.1]** | **<.0001** | **0.047** | 0.0<br>[ -0.5, 0.5] | >0.1 | <.001 | **2.2***<br>[ 1.3, 3.1]** | **<.0001** | **0.001** |

### Summary figures for age, sex, and hemispheric asymmetry effects

Supplemental Figure 10 presents the heatmap of □$^{2}_{G}$ values for the main effects of age, sex, and hemispheric asymmetry for each metric and ROI to provide the visual summary of the effect sizes, similar to Figure 2, 5, and 6 in the main text. Supplemental Figures 11 - 18 show the age scatter plots with regression lines for each sex, and for 21 pairs of ROI in each hemisphere, each hemisphere, for each metric and ROI (individual plots from the selected metric/ROI are shown in Figure 3 and 6 of the main text), at mean eTIV. Additionally, Supplemental Figures 19 - 26 show the line plots of hemispheric asymmetry in each sex for each metric and ROI. Values for the mean values in each sex/hemisphere represent predicted values for female or male subjects with the group mean eTIV and age to allow comparisons of values after eTIV difference is taken into account.

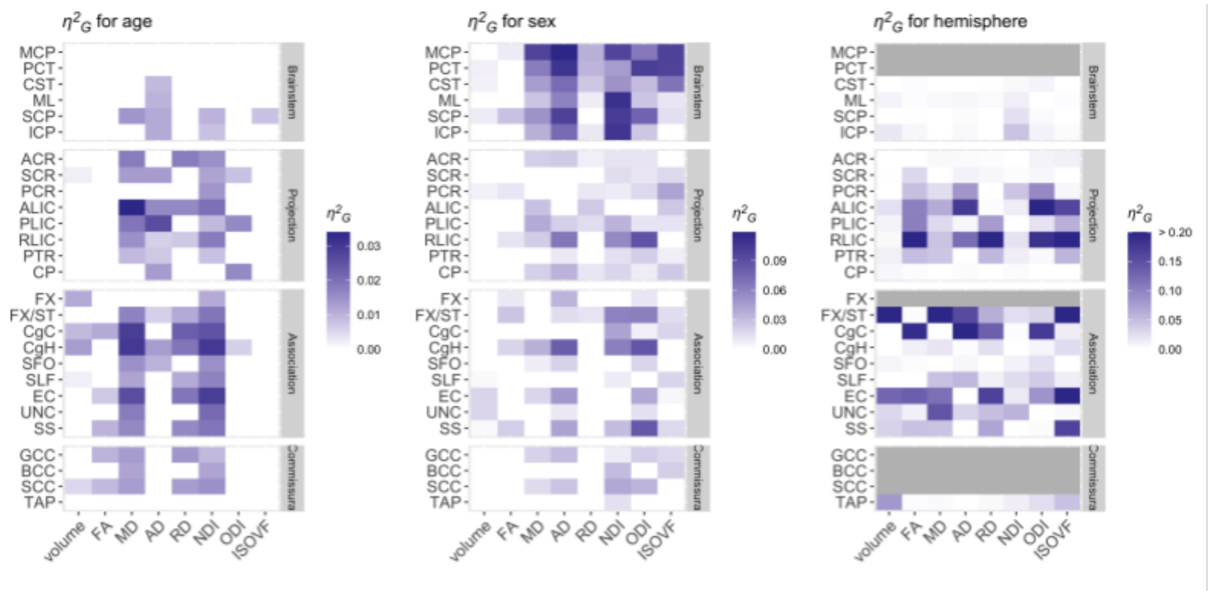

**Supplemental Figure 10. Effect size heatmaps for age, sex, and hemispheric asymmetry.** The heatmap of □$^{2}_{G}$ across metrics and ROIs is shown for the main effects of (A) age, (B) sex, and (C) hemisphere. Those that did not survive Bonferroni corrections for multiple comparisons were filtered out (set to 0) to facilitate comparisons within significant results. See Table 2 in the main text for the full names of the abbreviated ROIs.

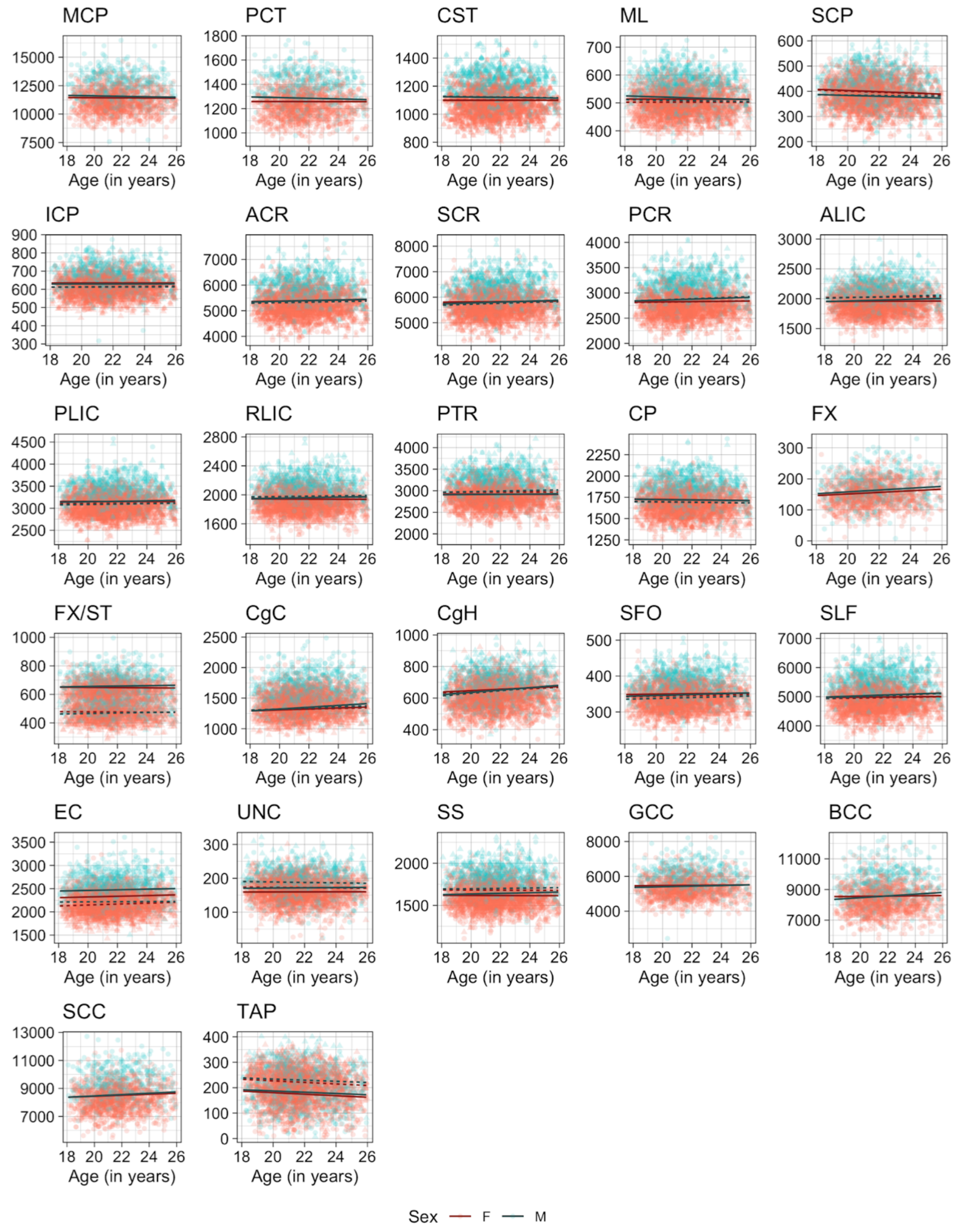

**Supplemental Figure 11. Scatter plots of individual age effects on WM volumes (in mm$^3$) in each ROI.**

Predicted linear regression lines are superimposed for each sex (dark red: females, dark cyan: males) and hemisphere (solid: left, dahed: right) at the mean eTIV value. See Table 2 in the main text for the full names of the abbreviated ROIs.

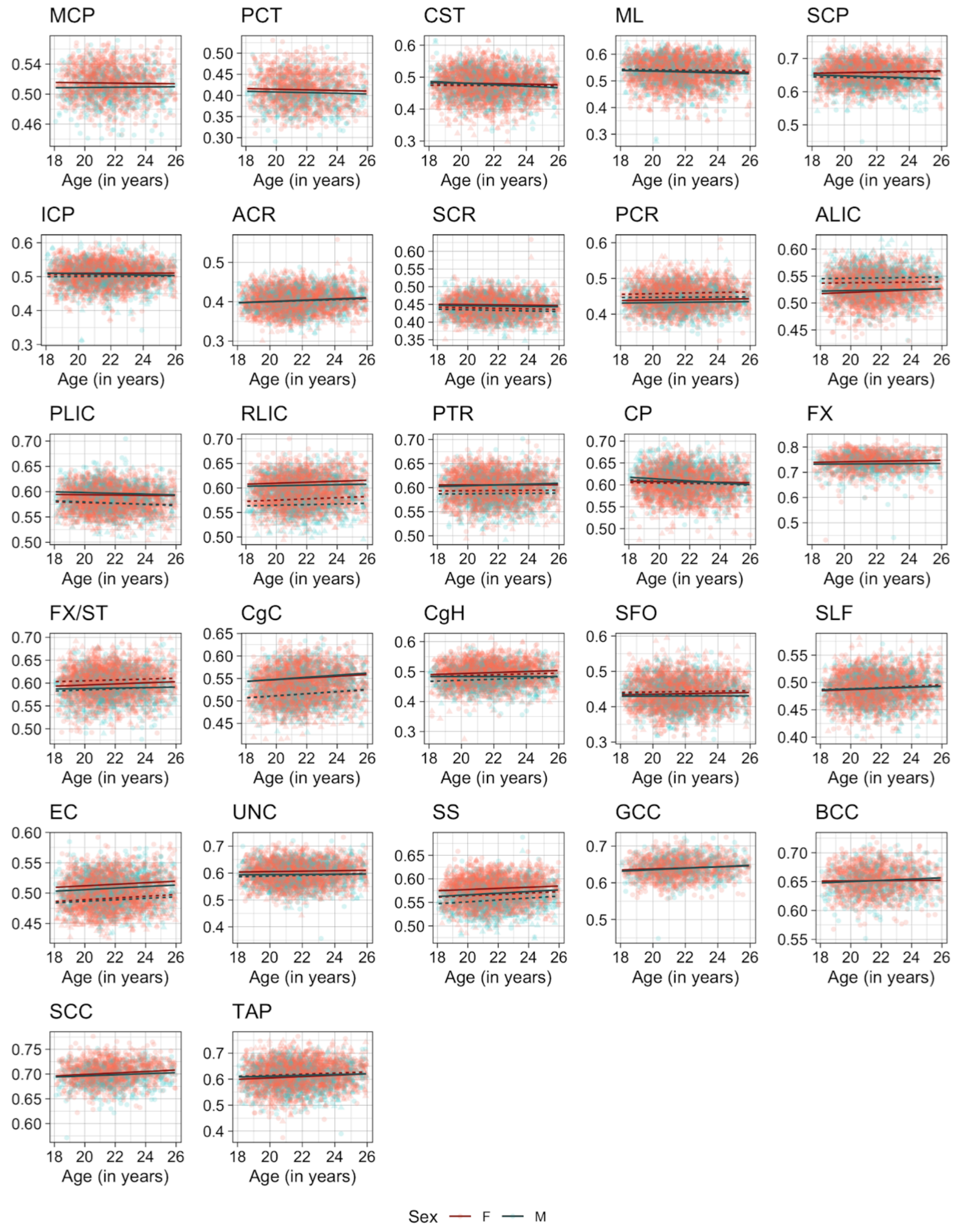

**Supplemental Figure 12. Scatter plots of individual age effects on WM FA values in each ROI.**

Predicted linear regression lines are superimposed for each sex (dark red: females, dark cyan: males) and hemisphere (solid: left, dahed: right) at the mean eTIV value. See Table 2 in the main text for the full names of the abbreviated ROIs.

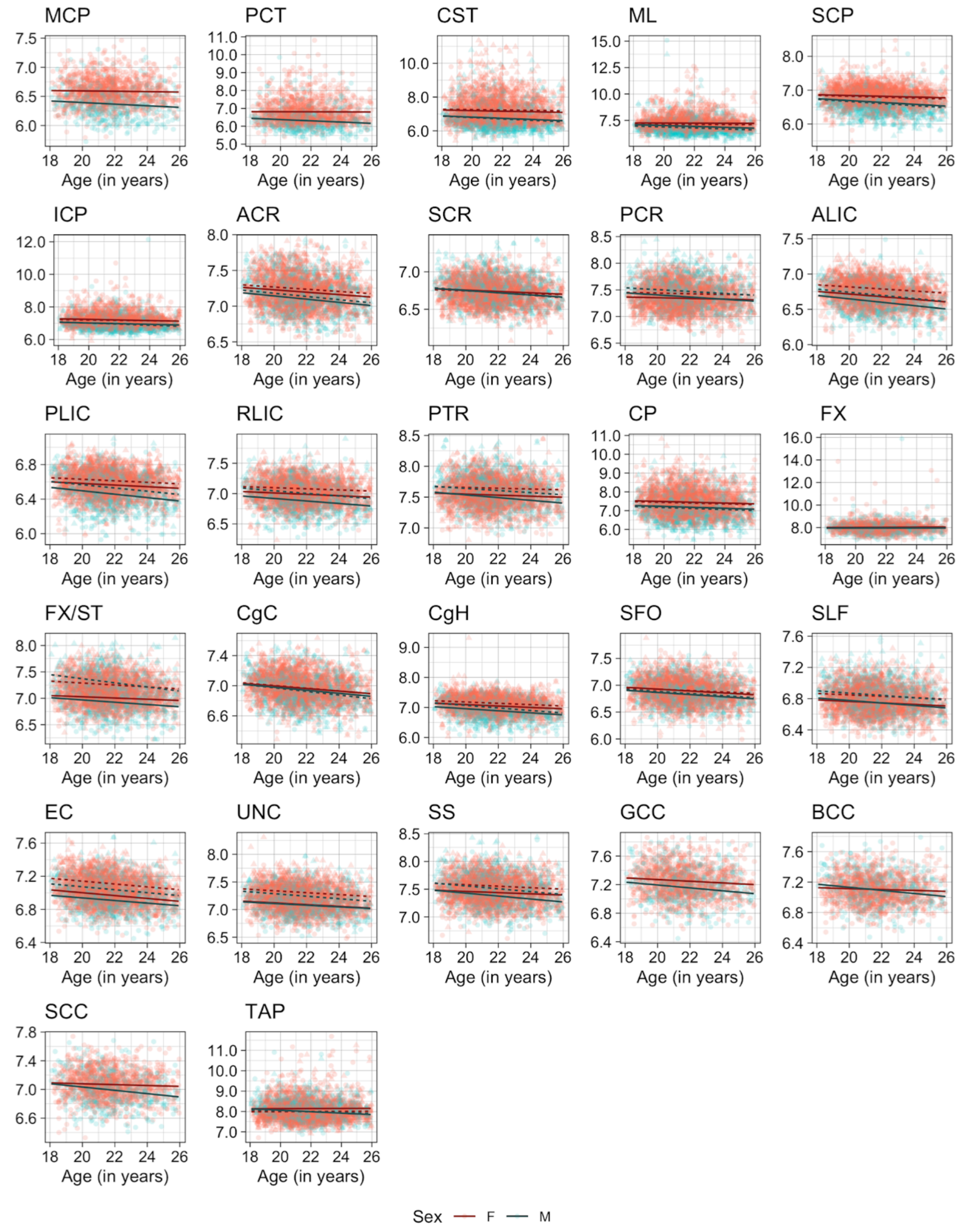

**Supplemental Figure 13. Scatter plots of individual age effects on WM MD values (x$10^{-4}$ $mm^2$/sec) in each ROI.**

Predicted linear regression lines are superimposed for each sex (dark red: females, dark cyan: males) and hemisphere (solid: left, dahed: right) at the mean eTIV value. See Table 2 in the main text for the full names of the abbreviated ROIs.

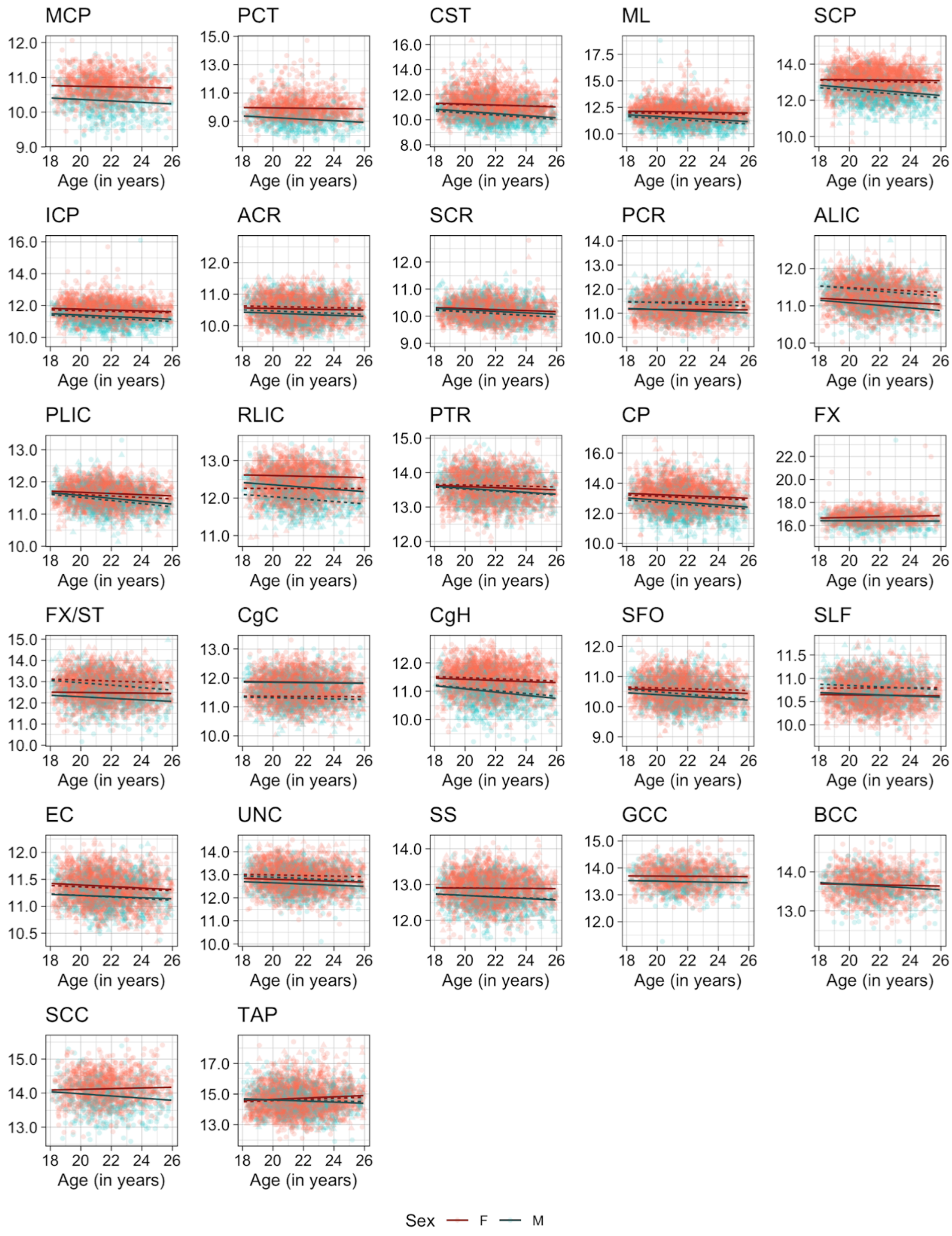

**Supplemental Figure 14. Scatter plots of individual age effects on WM AD values (x$10^{-4}$ $mm^2$/sec) in each ROI.**

Predicted linear regression lines are superimposed for each sex (dark red: females, dark cyan: males) and hemisphere (solid: left, dahed: right) at the mean eTIV value. See Table 2 in the main text for the full names of the abbreviated ROIs.

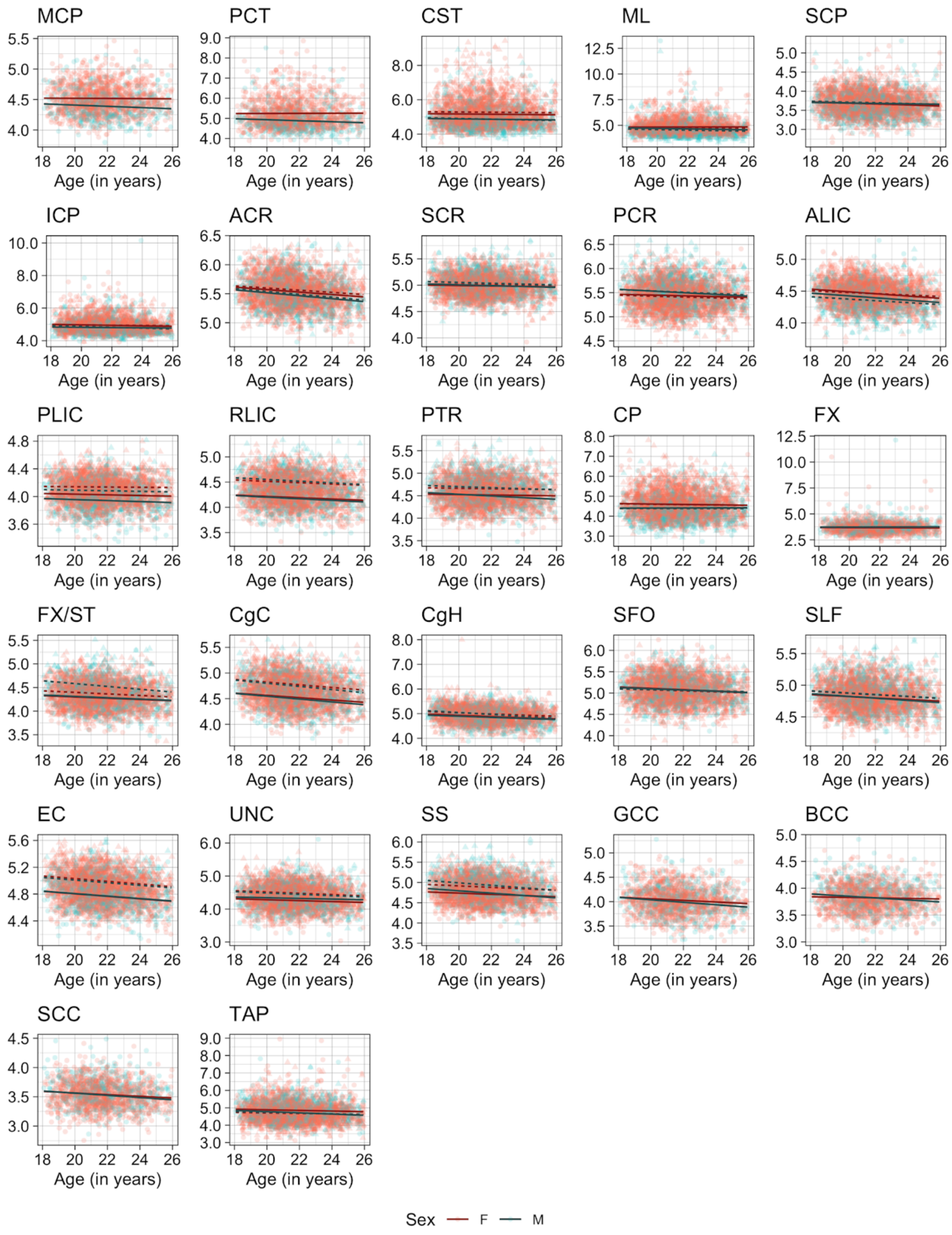

**Supplemental Figure 15. Scatter plots of individual age effects on WM RD values (x10$^{-4}$ mm$^2$/sec) in each ROI.**

Predicted linear regression lines are superimposed for each sex (dark red: females, dark cyan: males) and hemisphere (solid: left, dahed: right) at the mean eTIV value. See Table 2 in the main text for the full names of the abbreviated ROIs.

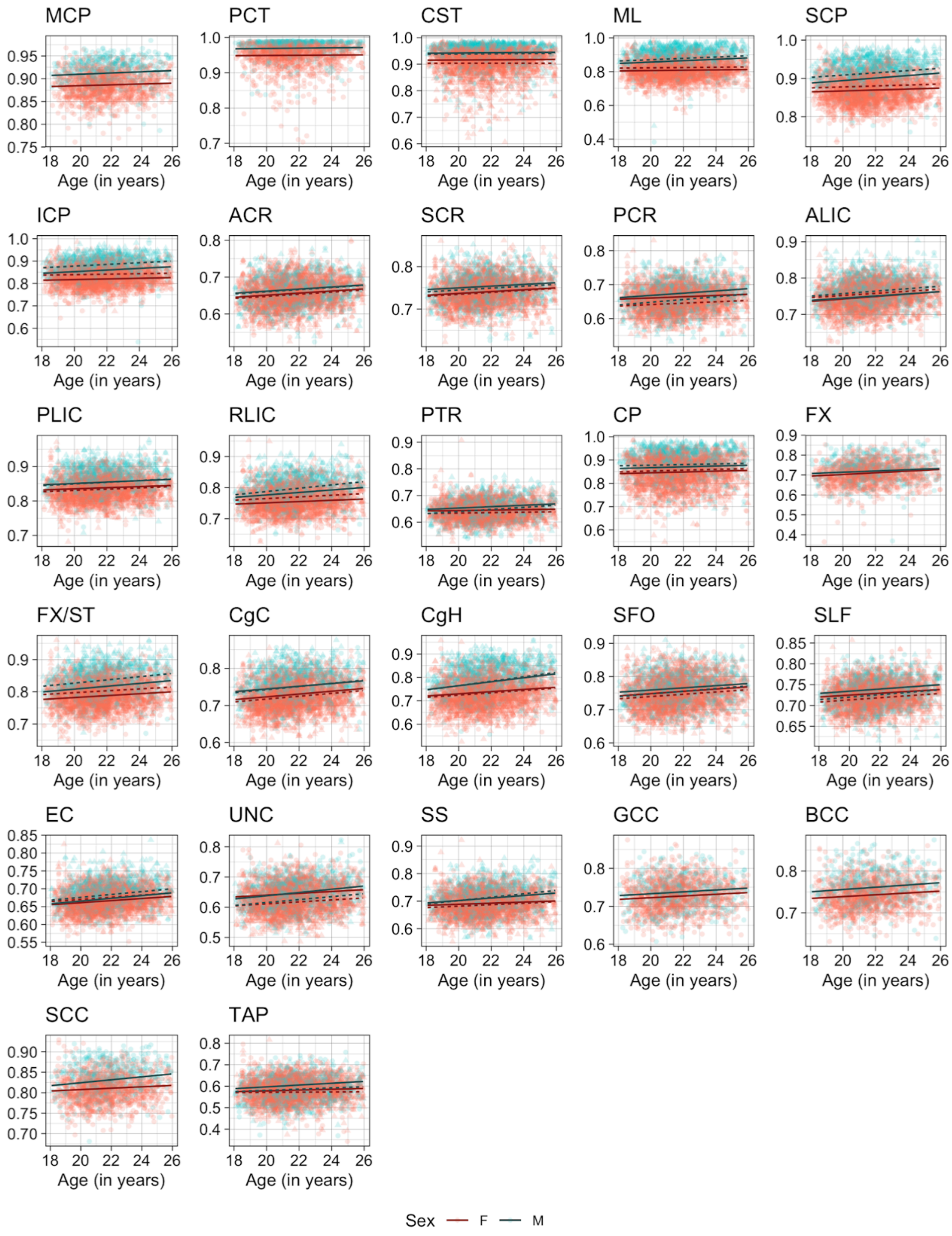

**Supplemental Figure 16. Scatter plots of individual age effects on WM NDI values in each ROI.**

Predicted linear regression lines are superimposed for each sex (dark red: females, dark cyan: males) and hemisphere (solid: left, dahed: right) at the mean eTIV value. See Table 2 in the main text for the full names of the abbreviated ROIs.

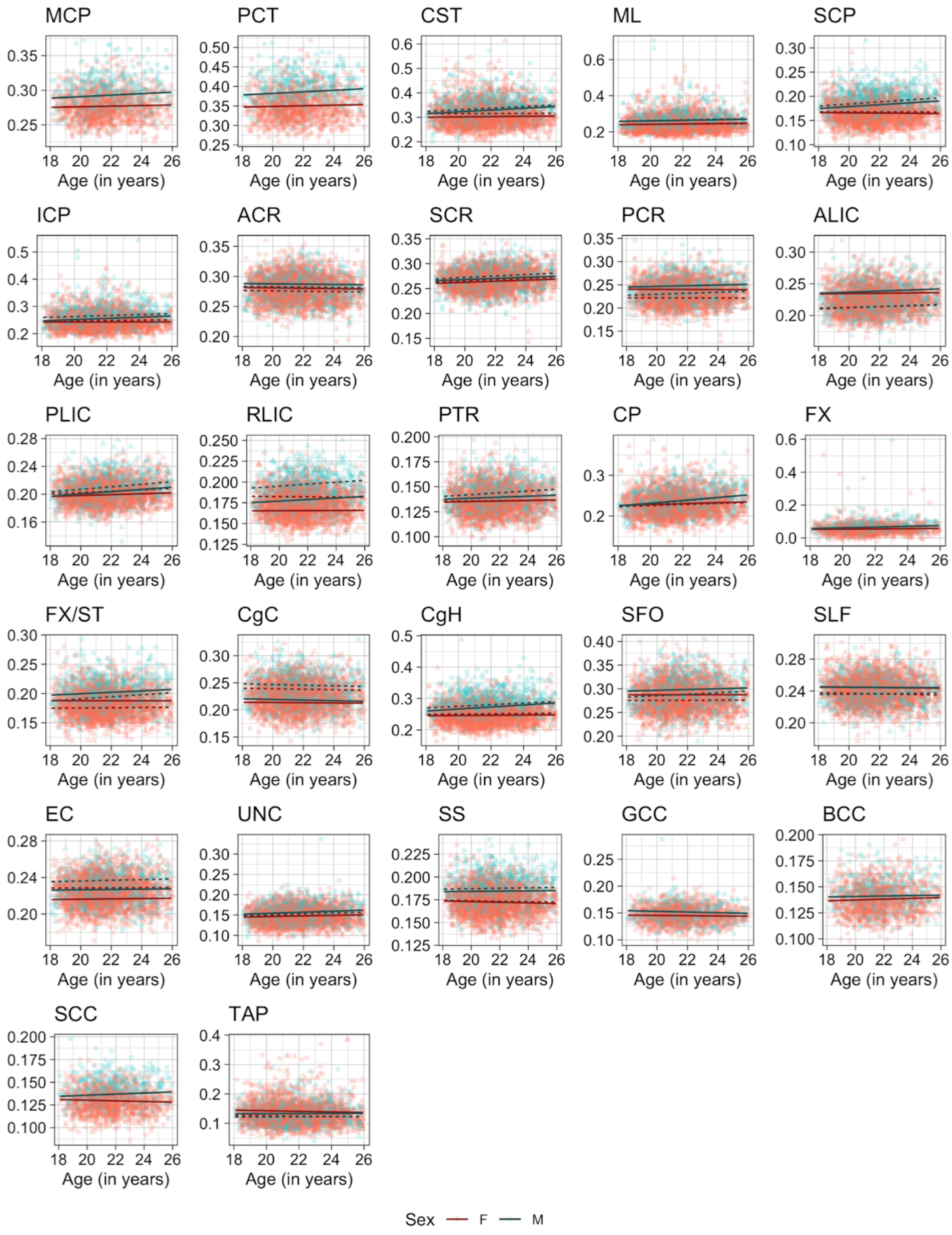

**Supplemental Figure 17. Scatter plots of individual age effects on WM ODI values in each ROI.**

Predicted linear regression lines are superimposed for each sex (dark red: females, dark cyan: males) and hemisphere (solid: left, dahed: right) at the mean eTIV value. See Table 2 in the main text for the full names of the abbreviated ROIs.

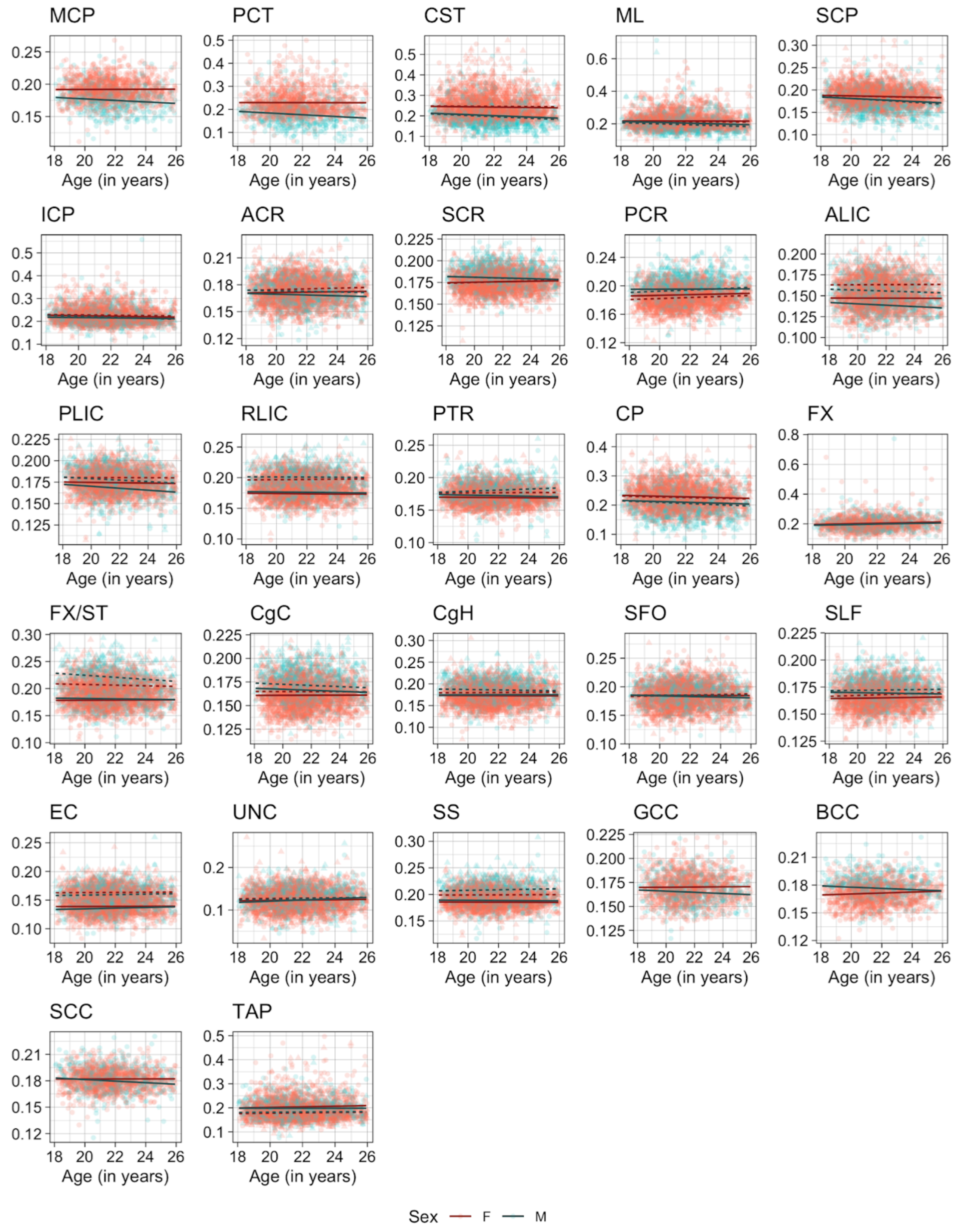

**Supplemental Figure 18. Scatter plots of individual age effects on WM IsoVF values in each ROI.**

Predicted linear regression lines are superimposed for each sex (dark red: females, dark cyan: males) and hemisphere (solid: left, dahed: right) at the mean eTIV value. See Table 2 in the main text for the full names of the abbreviated ROIs.

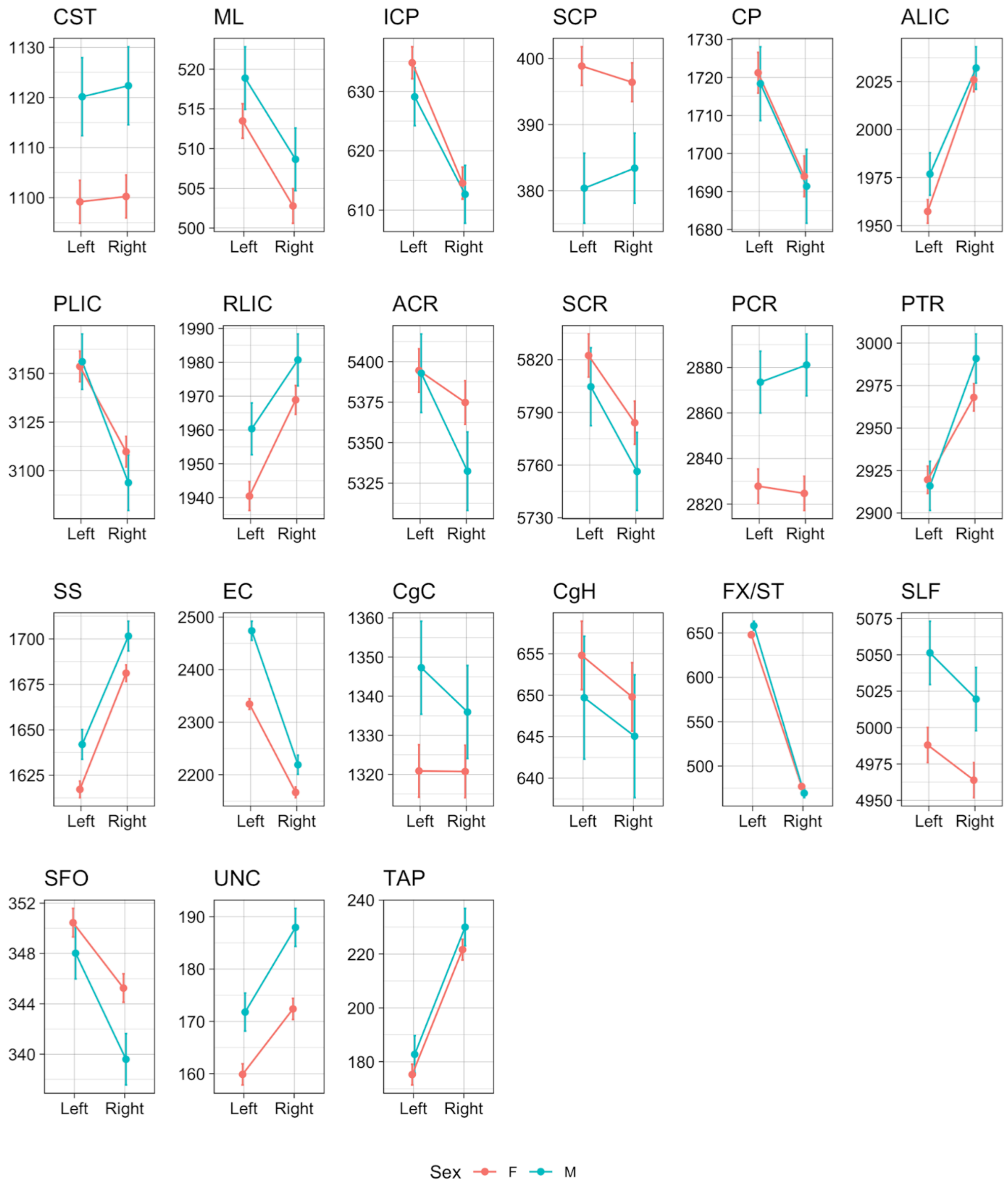

**Supplemental Figure 19. Line plots of individual hemisphere effects on WM volumes (in mm$^3$) in each ROI.**

Predicted mean values for each hemisphere at mean age and eTIV were computed for each sex (pink: females, blue: males). Error bars indicate 95% confidence intervals for predicted values. See Table 2 in the main text for the full names of the abbreviated ROIs.

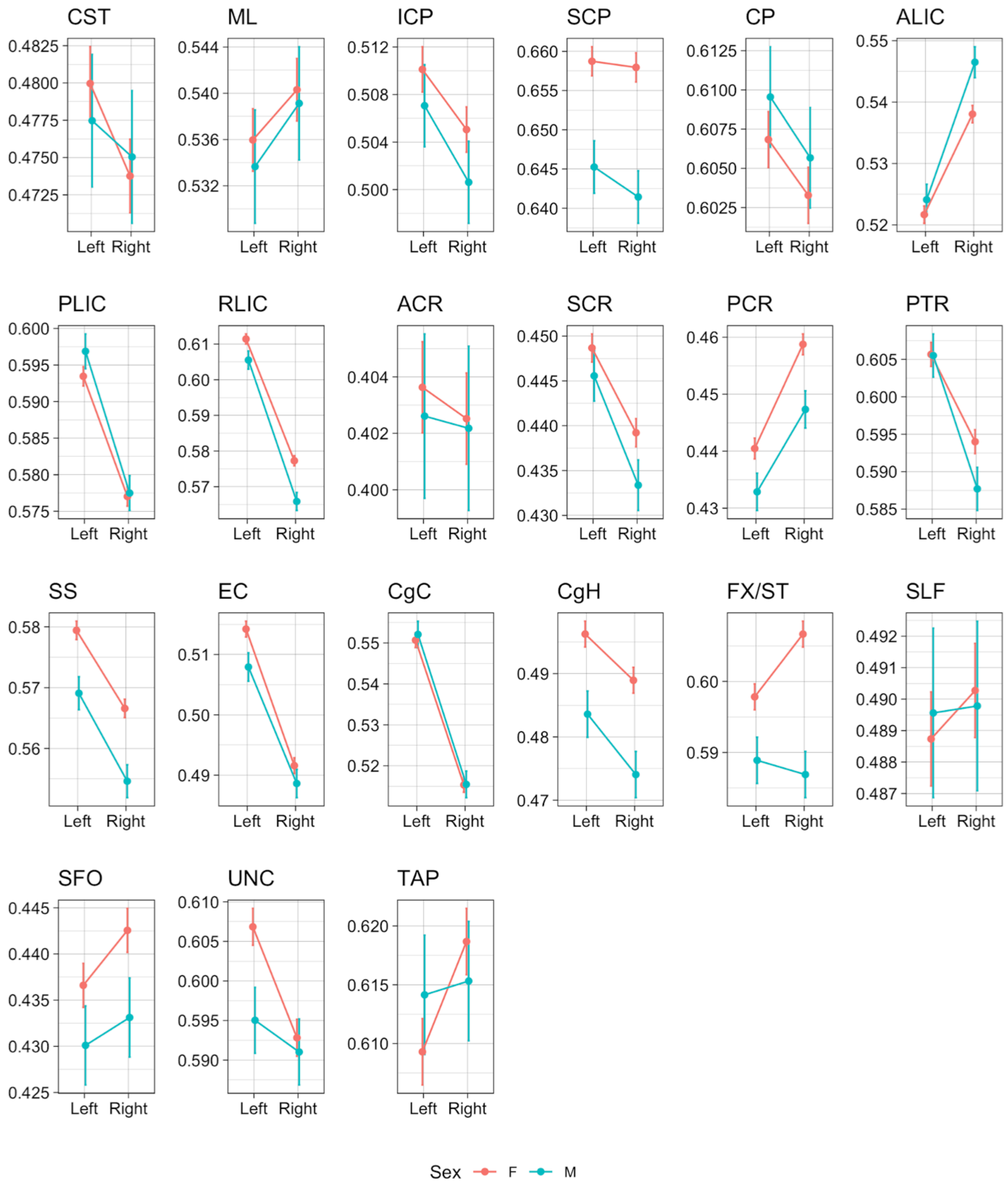

**Supplemental Figure 20. Line plots of individual hemisphere effects on WM FA values in each ROI.**

Predicted mean values for each hemisphere at mean age and eTIV were computed for each sex (pink: females, blue: males). Error bars indicate 95% confidence intervals for predicted values. See Table 2 in the main text for the full names of the abbreviated ROIs.

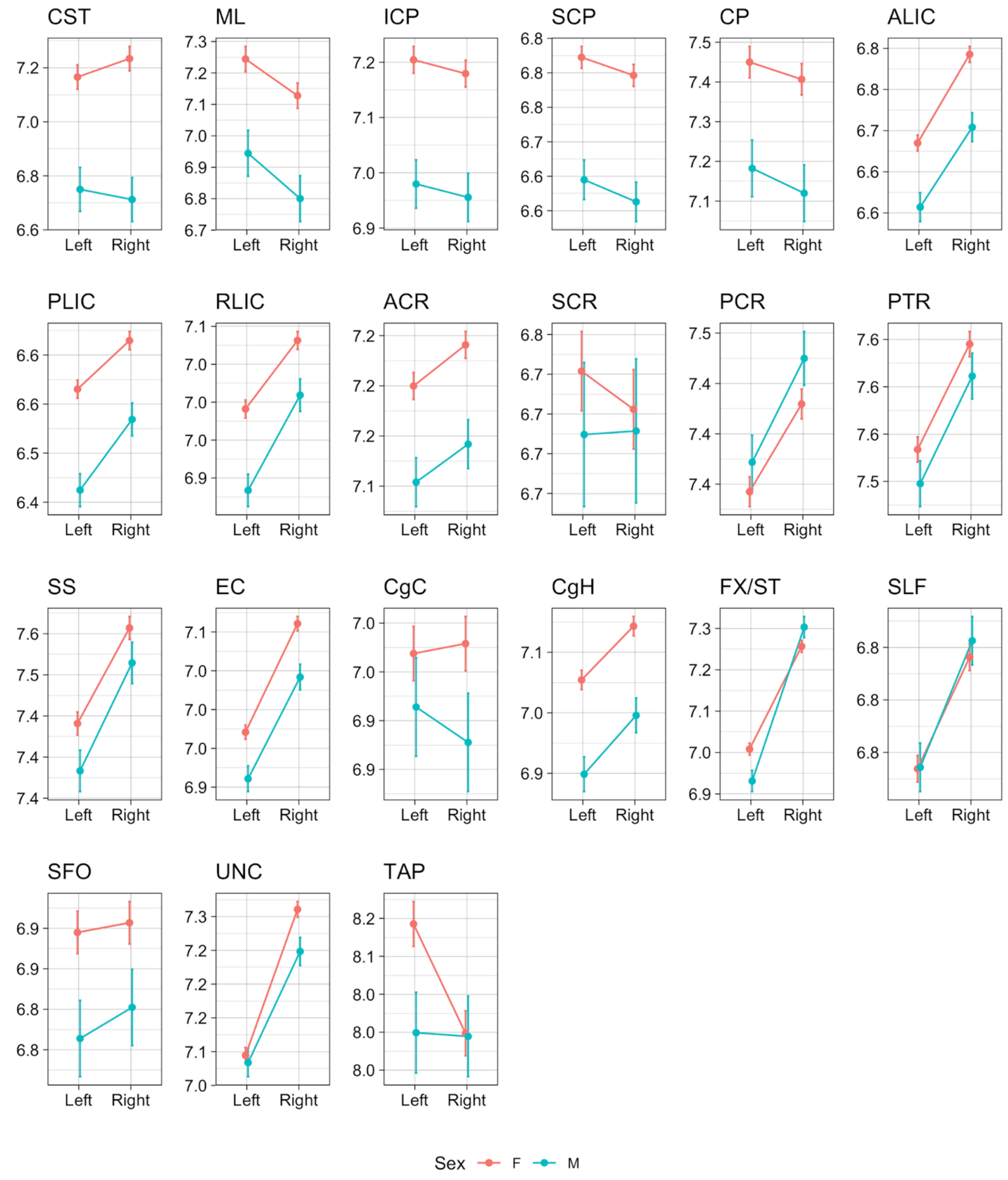

**Supplemental Figure 21. Line plots of individual hemisphere effects on WM MD values ($x10^{-4}$ $mm^2/sec$) in each ROI.**

Predicted mean values for each hemisphere at mean age and eTIV were computed for each sex (pink: females, blue: males). Error bars indicate 95% confidence intervals for predicted values. See Table 2 in the main text for the full names of the abbreviated ROIs.

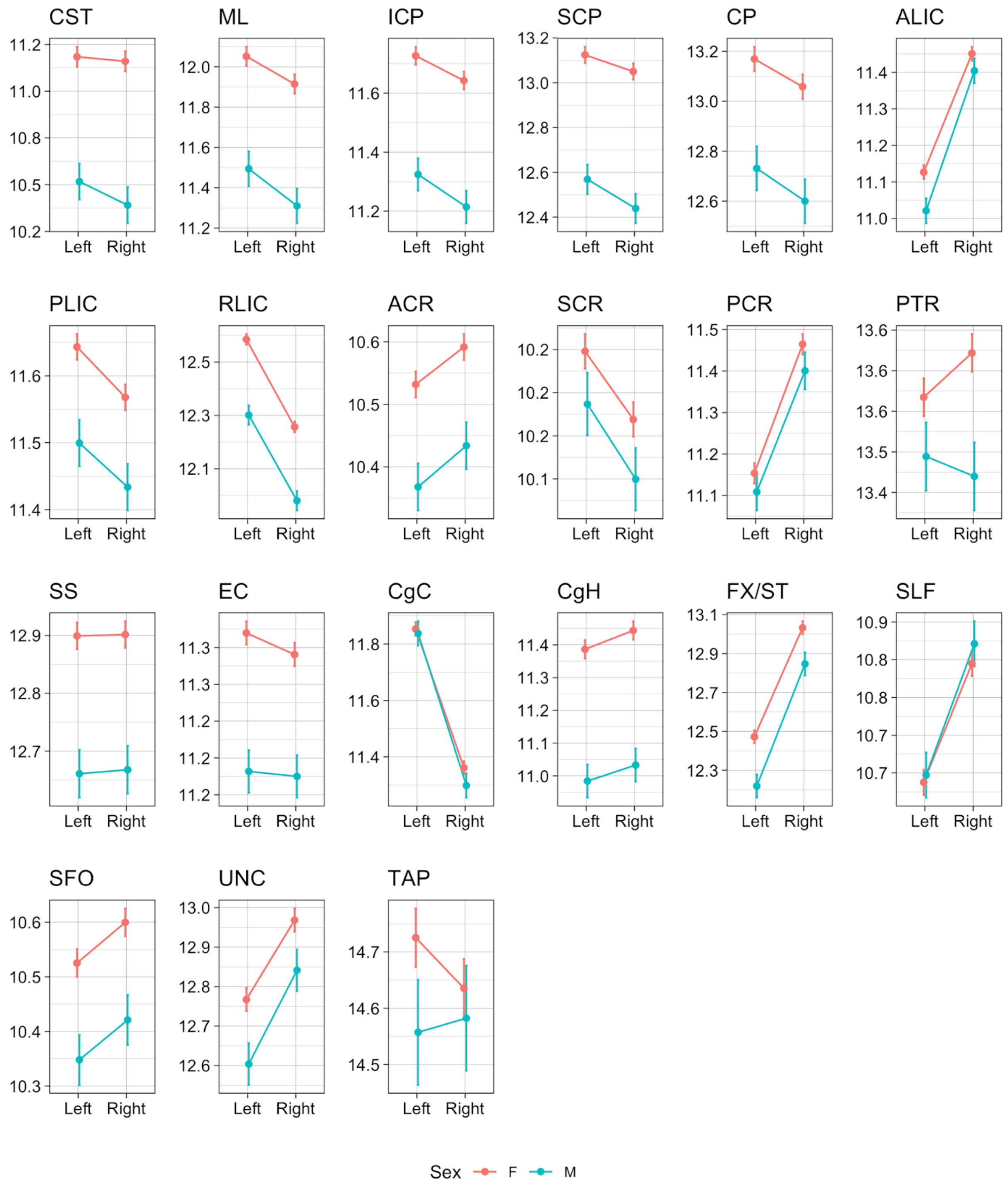

**Supplemental Figure 22. Line plots of individual hemisphere effects on WM AD values ($x10^{-4}$ $mm^2$/sec) in each ROI.**

Predicted mean values for each hemisphere at mean age and eTIV were computed for each sex (pink: females, blue: males). Error bars indicate 95% confidence intervals for predicted values. See Table 2 in the main text for the full names of the abbreviated ROIs.

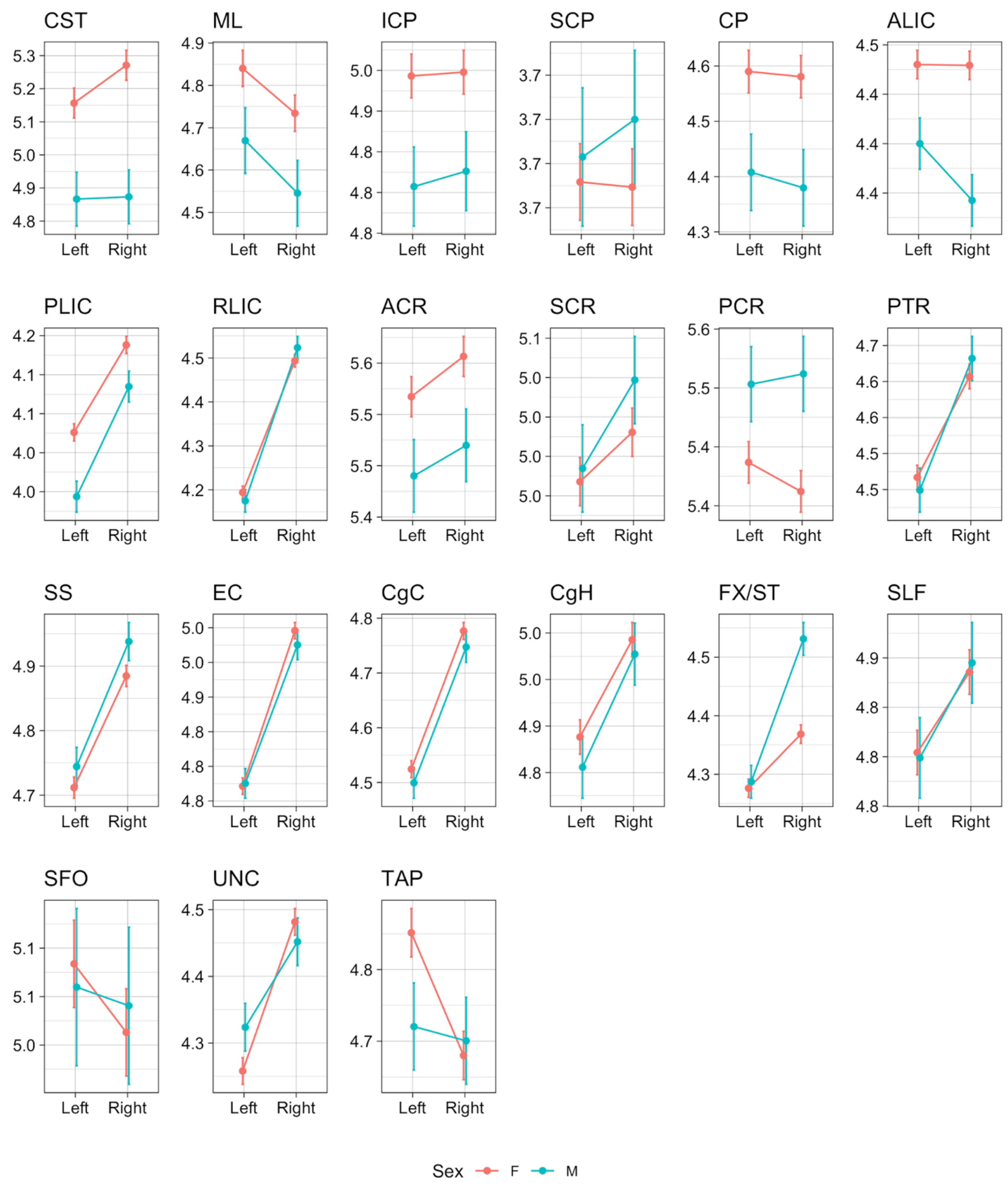

**Supplemental Figure 23. Line plots of individual hemisphere effects on WM RD values ($x10^{-4}$ $mm^2$/sec) in each ROI.**

Predicted mean values for each hemisphere at mean age and eTIV were computed for each sex (pink: females, blue: males). Error bars indicate 95% confidence intervals for predicted values. See Table 2 in the main text for the full names of the abbreviated ROIs.

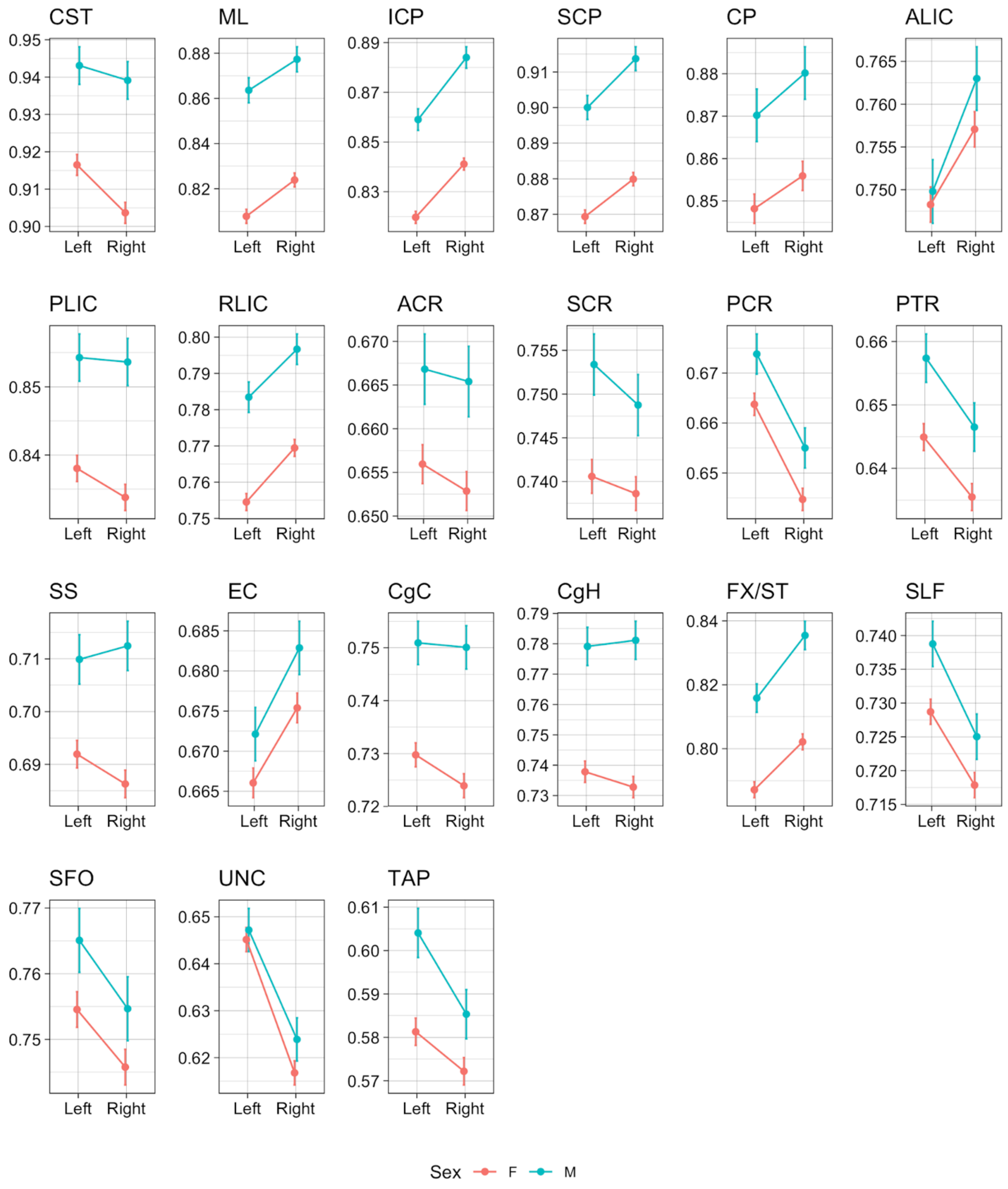

**Supplemental Figure 24. Line plots of individual hemisphere effects on WM NDI values in each ROI.**

Predicted mean values for each hemisphere at mean age and eTIV were computed for each sex (pink: females, blue: males). Error bars indicate 95% confidence intervals for predicted values. See Table 2 in the main text for the full names of the abbreviated ROIs.

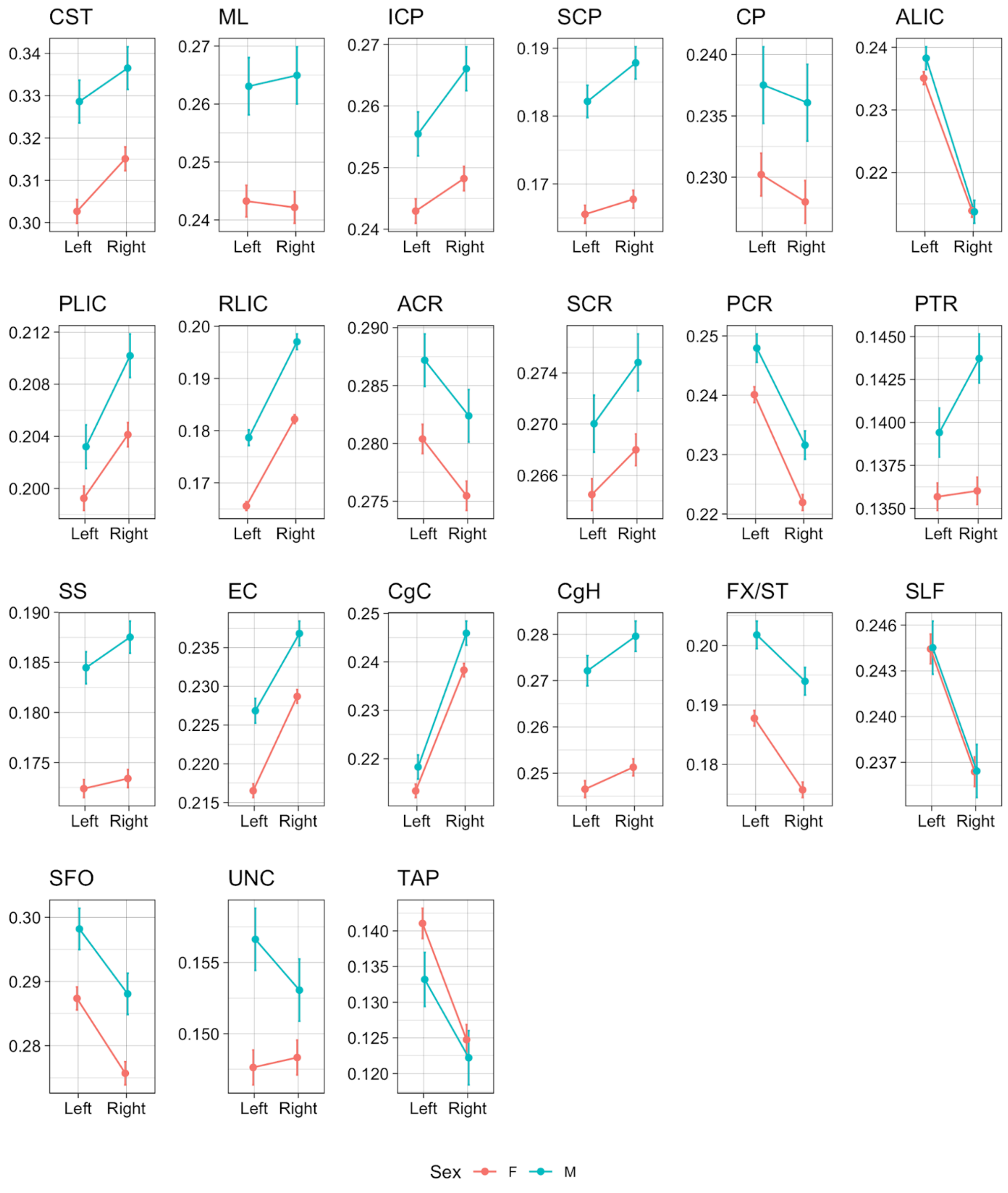

**Supplemental Figure 25. Line plots of individual hemisphere effects on WM ODI values in each ROI.**

Predicted mean values for each hemisphere at mean age and eTIV were computed for each sex (pink: females, blue: males). Error bars indicate 95% confidence intervals for predicted values. See Table 2 in the main text for the full names of the abbreviated ROIs.

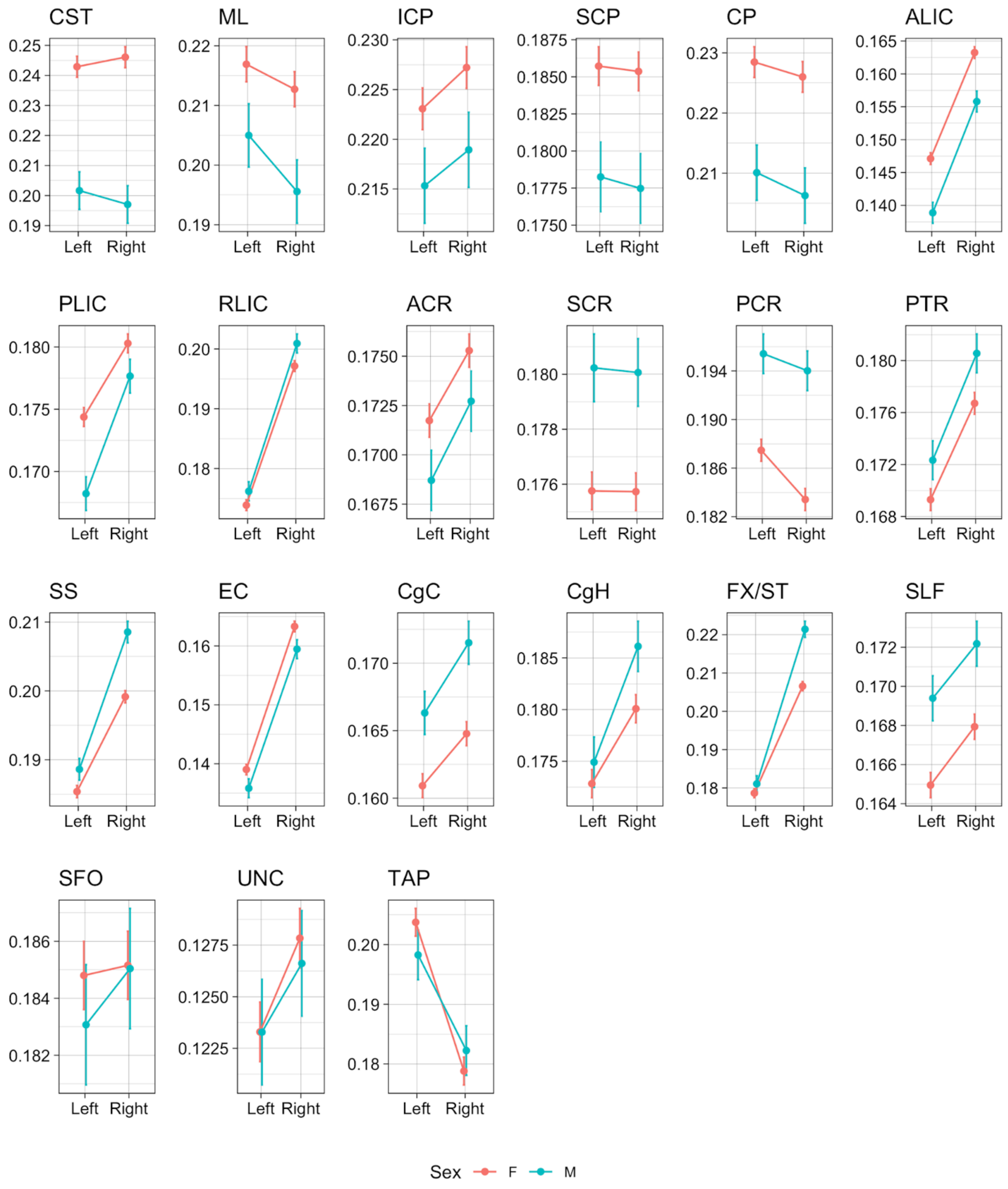

**Supplemental Figure 26. Line plots of individual hemisphere effects on WM IsoVF values in each ROI.**

Predicted mean values for each hemisphere at mean age and eTIV were computed for each sex (pink: females, blue: males). Error bars indicate 95% confidence intervals for predicted values. See Table 2 in the main text for the full names of the abbreviated ROIs.

### Effects of outliers on the estimates of age, sex, and hemispheric asymmetry

As we showed above, mean DTI/NODDI metrics for the very small ROIs and brainstem ROIs tended to be noisy and susceptible to small cross-modality misalignments in the images, resulting in skewed distributions and extreme outliers in these ROIs. Such noise in the data likely reduce the power to detect effects of interest, such as age and sex. Given the large sample size of our data, however, we believed such effects to be relatively small, and thus reported the results without any outlier removal in the main manuscript. Here, we demonstrate how the data 'cleaning' by removing extreme outliers can modify the estimates of age, sex, and hemispheric asymmetry.

Supplemental figure 27 shows the comparisons of standardized parameter estimates ($\beta^*$) for age, sex, and hemispheric asymmetry effects for the model with and without the removal of extreme outliers, defined as those below or above 3*IQR from the first or third quartile, respectively (i.e. *red* data points in Figures S2 - S9 above). As can been seen, the outlier removal had very little effects on the estimated $\beta^*$ overall, only changing the magnitudes of $\beta^*$ very slightly or affecting the significance inference in some (e.g. age effect for MD in ICP, sex effect for IsoVF in ICP).

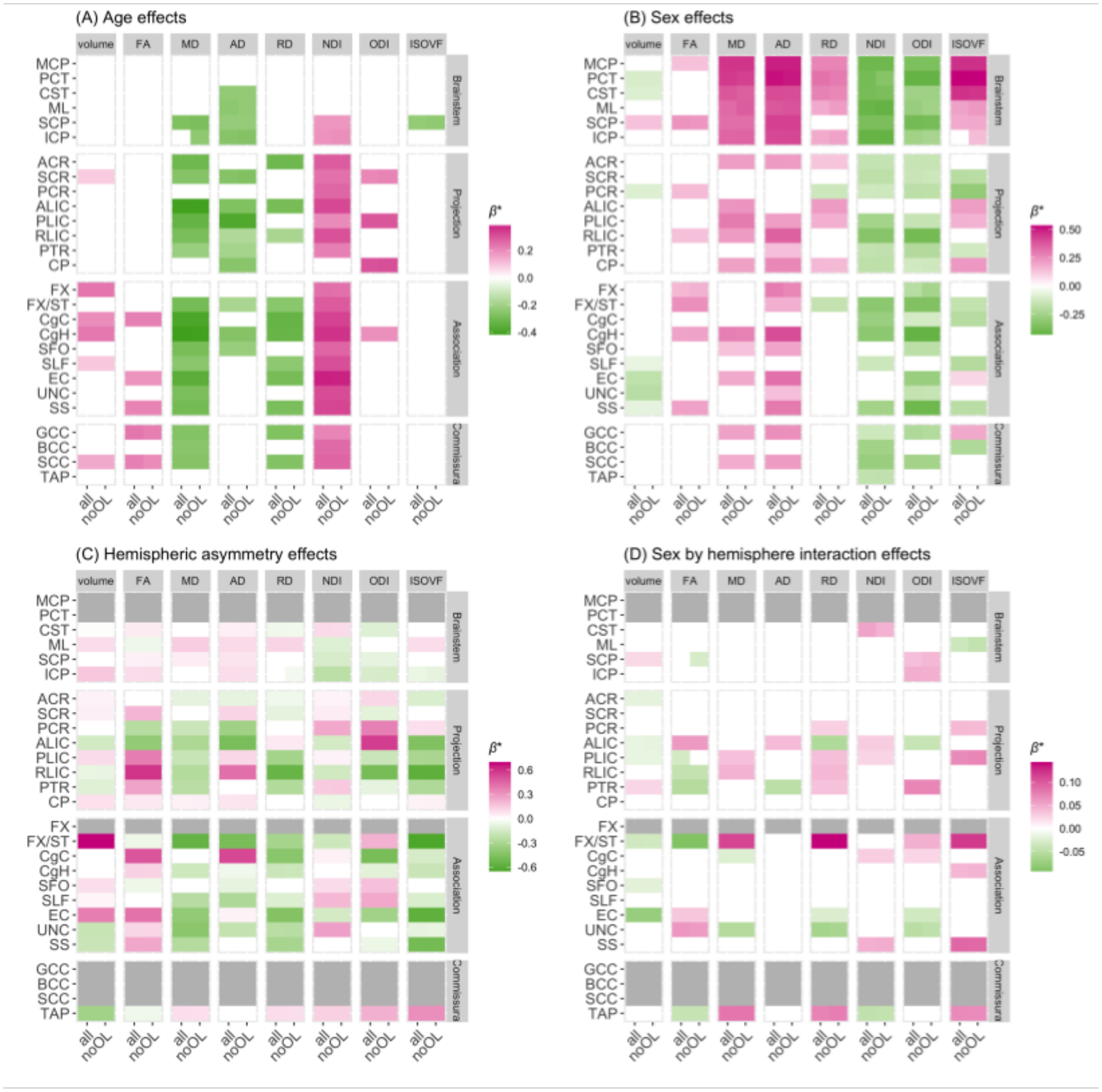

**Supplemental Figure 27. Impact of outlier removal on the standardized estimates of age, sex, and hemispheric asymmetry effects on the different WM IDPs in the JHU ROIs.**

The standardized estimates ($\beta^*$) of (A) age, (B) sex, (C) hemispheric asymmetry, and (D) sex by hemisphere interaction effects are shown, filtering out those that did not survive Bonferonni corrections for multiple comparisons to facilitate the comparisons of significant effects. The $\beta^*$ values for the model without outlier removal (“all”) and with extreme outlier removal (“noOL”) are compared side by side. See Table 2 in the main text for the full names of the abbreviated ROIs.

### Effects of motion parameters during the DWI scan

Pines et al (2020) recently demonstrated that DTI and NODDI metrics are sensitive to in-scanner motion, and that after accounting for age and sex effects, these metrics correlated with the mean relative root mean square (RMS) of the displacement, a measure of the in-scanner head motion (Pines et al., 2020). We quantified the association of WM property metrics in each tract with this parameter in our dataset, and also investigated how the addition of RMS as a covariate in the model influenced the estimates of age, sex, and hemispheric asymmetry effects.

Supplemental Figure 28 shows the comparison of model quality between the models with and without RMS as a covariate for the different DTI/NODDI IDPs and JHU ROIs, as judged by the BIC and total variance explained (adjusted $R^2$ or marginal $R^2$). It shows that in our dataset, the addition of the RMS did not improve the model fit or noticeably increased the total variance explained. Supplemental Figure 29 shows the $\beta$* values of the RMS itself in the model with RMS, as well as any changes in the $\beta$* values for age and sex effects with and without the RMS in the model. While the mean DTI/NODDI values in a handful of ROIs were significantly related to the RMS positively or negatively, the effect sizes were very small, accounting for less than 1 % of variance in all cases (maximum $\eta^2_G$ = 0.008). More importantly, their impact on the $\beta$* values for age or sex was minimal, with no visible changes in the $\beta$* values. As the RMS values do not differ for the right and left hemisphere ROIs, the inclusion of RMS in the model was not expected to affect the estimates for hemispheric asymmetry (except through its influence on the amount of variance explained, which may affect the level of significance), and it did not (data not shown).

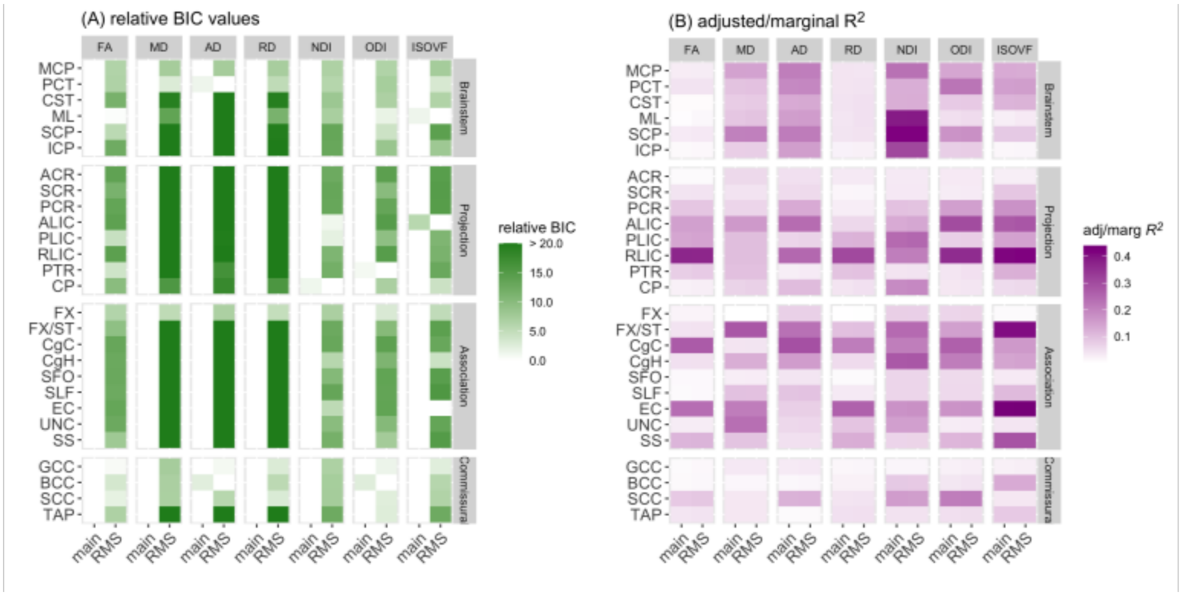

**Supplemental Figure 28. Comparison of model quality between models with and without RMS for the DTI/NODDI IDPs and JHU ROIs.**

(A) Relative Bayesian information criterion (BIC), defined as the difference between the BIC values between a given model and the most parsimonious model (with minimum BIC), comparing the model with (“RMS”) and without (“main”) RMS as a covariate. (B) Adjusted or marginal $R^2$, comparing the “RMS” and “main” models. See Table 2 in the main text for the full names of the abbreviated ROIs.

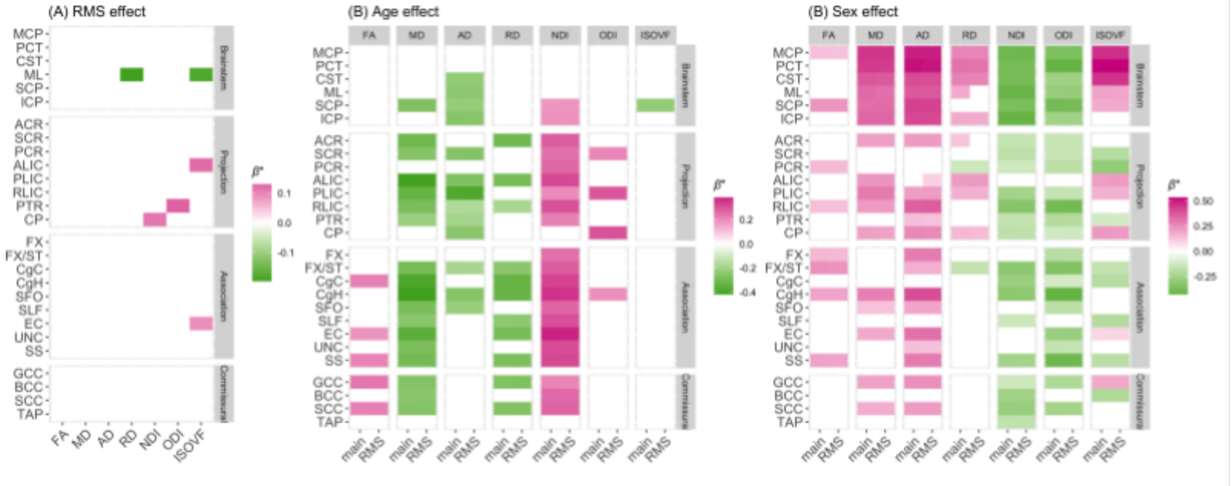

**Supplemental Figure 29. In-scanner motion effects and their impact on the estimates of age and sex effects for the different DTI/NODDI IDPs and JHU ROIs.**

The standardized estimates ($\beta^*$) of (A) RMS, (B) age, and (C) sex effects are shown for each DTI/NODDI metric in the JHU ROIs, filtering out those that did not survive Bonferonni corrections for multiple comparisons to facilitate the comparisons of significant effects. The $\beta^*$

values for age and sex for models with (“RMS”) or without (“main”) RMS as a covariate are compared side by side. See Table 2 in the main text for the full names of the abbreviated ROIs.

### Comparison of quadratic versus linear age effect models

While the developmental trajectory of WM properties is known to be non-linear (Lebel et al., 2019), maturational changes are likely to be much slower at the age range of our sample compared to earlier in the development. We expected that the age range is too narrow to capture any non-linearity during this period. In a preliminary analysis, we compared the model quality of linear versus quadratic age effect models.

Supplemental Figure 30 shows the comparison of model quality between the two models for all eight metrics and 27 ROIs, as judged by the BIC and total variance explained (adjusted $R^2$ or marginal $R^2$). The BIC values were almost always smaller for linear than quadratic age models across metrics and ROIs, and the total variance explained by the inclusion of quadratic age term did not change noticeably in any of the metric/ROI combinations.

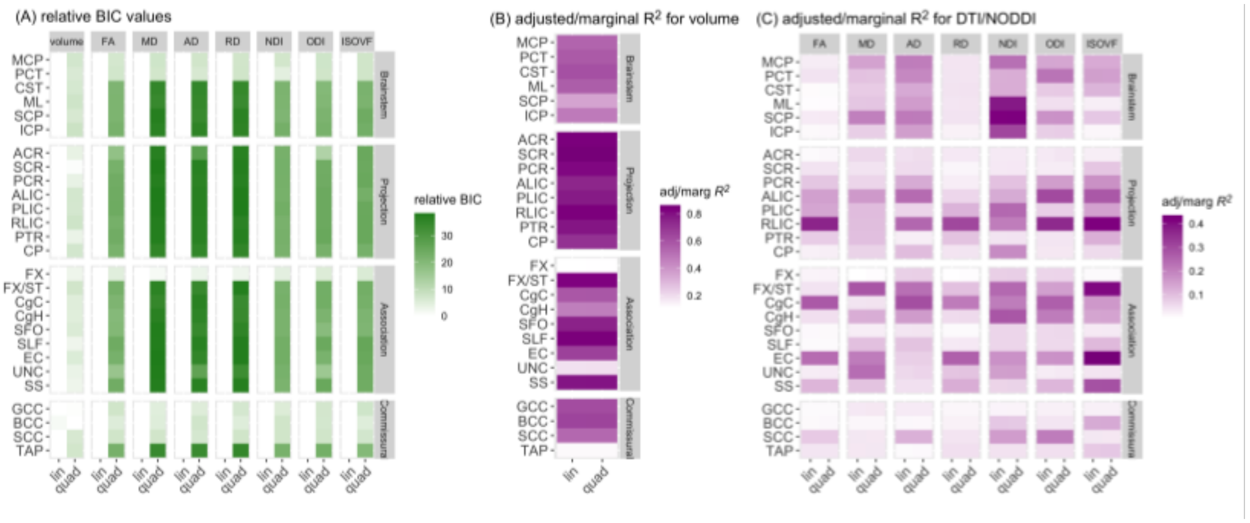

**Supplemental Figure 30. Comparison of linear and quadratic age model quality for the different WM IDPs and JHU ROIs.**

(A) Relative Bayesian information criterion (BIC), defined as the difference between the BIC values between a given model and the most parsimonious model (with minimum BIC), and adjusted or marginal $R^2$ values for the (B) WM volume and (C) mean DTI/NODDI in each JHU ROI, are compared side by side for the linear (“lin”) and quadratic (“quad”) age effect models. See Table 2 in the main text for the full names of the abbreviated ROIs.

### Effects of volume corrections

#### Effects of global volume correction on regional WM volumetry

The estimates of age and sex effects on the regional WM volumes partly depend on if and how global brain volume or head size is corrected. While we reported the age and sex effects on regional WM volumes in a model that included eTIV as a covariate in the main manuscript, there are inconsistencies in how global brain size is corrected (or not corrected) in the literature (e.g. Table 2 in a recent review by Vijayakumar et al., 2018). It is particularly important when quantifying the sex differences, since males typically have larger head size than females, but it can also affect the estimates for age-related changes if the overall brain volume itself vary with age. We have previously shown that the total WM volume (TWMV) significantly increased with age in this sample, only when controlling for overall head size by including eTIV in the model (Tsuchida et al., 2020). Here, we investigated how the estimates of age- and sex-related variations in the regional WM volumes were modulated by the inclusion of eTIV or TWMV in the model. We tested and compared the following variations of our primary models (1) and (2) described in the main manuscript:

1) "*noVol*" : model excluding the term $\beta_{eTIV} eTIV$ (i.e. no global volume correction)
2) "*eTIV*" : model that include the eTIV term (i.e. our primary model)
3) "*TWMV*": model that replaced the eTIV in the model with TWMV instead
4) "*eTIV + TWMV*": model that included both eTIV and TWMV

Supplemental Figure 31 compares the overall model quality for each of the four models, and also shows their impact on the $\beta^*$ values for age and sex. While the BIC values indicated that having both eTIV and TWMV almost always improved the overall model fit of the regional WM volumes across the ROIs, the total variances (adjusted or marginal $R^2$) explained by the model increased only marginally when including both volumes as covariates compared to having either eTIV or TWMV alone.

The comparison of $\beta^*$ values for age in the four models showed that the age-related increase in cingulum in the hippocampus (CgH) was significant regardless of how or if the global volume was controlled for in the model. The cingulum in the cingulate gyrus (CgC) also showed an age-related increase, but it did not survive multiple comparison correction when the global volume was not in the model (uncorrected $p$ = 0.00043). WM volume increases with age in superior corona radiata (SCR), superior longitudinal fasciculus (SLF), and splenium of corpus callosum (SCC) reached significance only when eTIV alone was in the model. In contrast, cerebral peduncle (CP) and superior cerebellar peduncle (SCP) showed a significant decrease with age when TWMV was taken into account.

For sex effects, males had significantly larger WM volumes than females across all the ROIs if the global volume is not taken into account. Many of these differences disappeared when eTIV or TWMV was included in the model, and in a few cases reversed, with females showing significantly larger relative volume than males, most notably in SCP. However, some ROIs exhibited attenuated but still significant sex differences, with males having larger volumes than females, after global volume corrections (e.g. external capsule (EC), uncinate fasciculus (UNC)).

Since the global volume values for the right and left ROIs are the same, the inclusion of the global volume should not affect the estimates of hemispheric asymmetry effects, although the degree of significance may be affected slightly through their impact on the explained variance. As expected, we did not observe any impact of global volume correction on the hemisphere asymmetry or sex by hemisphere interaction effects (not shown).

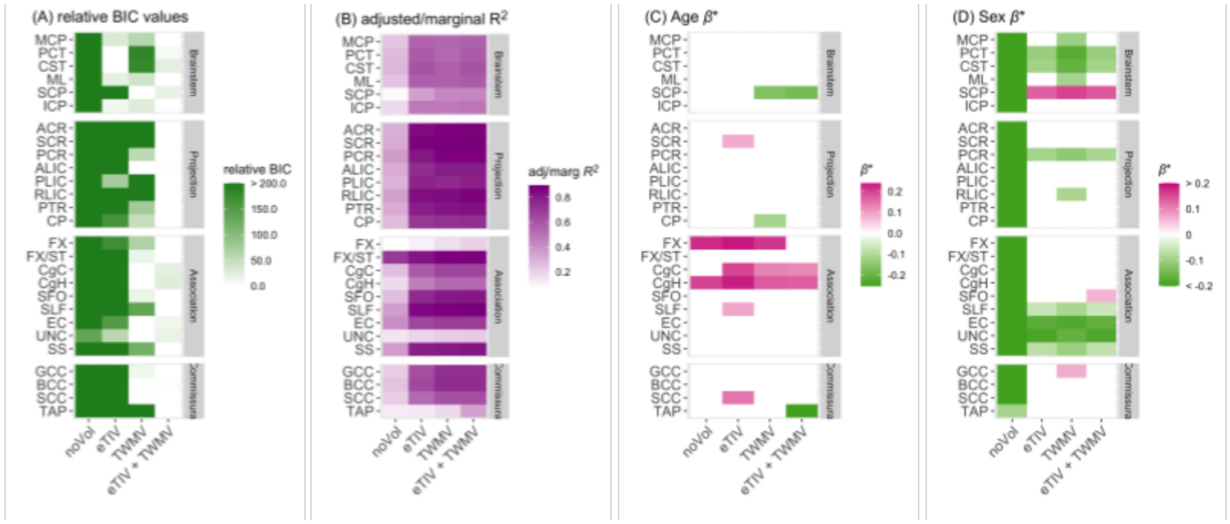

**Supplemental Figure 31. Effects of global volume correction on the model quality and the standardized estimates of age and sex effects in the WM JHU ROI volumes.**

Comparisons of (A) relative BIC values, (B) adjusted $R^2$, standardized parameter estimates *(β*)* for (C) age and (D) sex effects in four models with or without global volume covariates are shown for each WM JHU ROI volume. Those that did not survive Bonferroni corrections for multiple comparisons were filtered out (set to 0) to facilitate comparisons within significant results. See supplemental text for the description of the four models, and see Table 2 of the main text for the full names of abbreviated ROIs.

#### Effects of global and regional volume correction on DTI/NODDI metrics

Unlike for volumetry, the effects of global volume on mean DTI or NODDI values are not intuitive, and less well-understood; however, we have previously shown that eTIV significantly impacts mean DTI or NODDI metrics in WM skeleton or in WM mask (Beaudet et al., 2020; Tsuchida et al., 2020). For the regional mean values of DTI/NODDI metrics, a previous study has demonstrated that the mask volume for a given ROI could impact the mean DTI value inside the ROI, presumably due to the partial volume effect (Vos et al., 2011). Here we investigated the impact of the global volume (eTIV) or the local ROI volume (ROIV) on the mean DTI/NODDI metrics and the estimates of age, sex, and hemisphere asymmetry effects. We tested and compared the following models:

1) *noVol*: model excluding the term $\beta_{eTIV}eTIV$ (i.e. no global volume correction)
2) *eTIV*: model that includes the eTIV term (i.e. our primary model)
3) *ROIV*: model that replaced the eTIV with ROIV instead
4) *combined*: model that included eTIV, and ROIV

Supplemental Figure 32 compares the changes in model quality when the model (1) does not include eTIV or ROIV as a covariate (noVol), (2) only includes eTIV, or (3) ROIV, and (4) include both eTIV and ROIV. The BIC values indicated that the most parsimonious model varied across ROIs and DTI/NODDI metrics. Having global and/or local volumes as covariates tended to improve the model fit in particular for FA, NDI, and IsoVF values in a number of ROIs. The overall variance explained by the inclusion of these volumes did not visibly increase for the most part, except for NDI in many ROIs, where inclusion of eTIV visibly increased adjusted/marginal $R^2$.

Supplemental Figure 33 summarizes the contributions of eTIV and ROIV on DTI/NODDI metrics directly by visualizing the $\beta^*$ values of eTIV and ROIV terms in the models that include either eTIV or ROIV, respectively, and by showing the $\beta^*$ for age, sex, or hemisphere in the four models. Larger eTIV/ROIV was associated with higher NDI in many ROIs, although NDI in a few ROIs were negatively associated with eTIV (anterior corona radiata (ACR), TAP). They were also associated with higher IsoVF in most ROIs as well, except in the brainstem ROIs. For DTI metrics, the effects of eTIV and ROIV were more variable and mixed across different ROIs.

Despite the significant relationships between global and local volumes with the mean DTI/NODDI values in these ROIs, their impact on age effect estimates was minimal. Most of the age-related variations observed for DTI/NODDI metrics in the 27 ROIs remained significant across the models with or without eTIV and/or ROIV as covariates. The sex differences were modulated slightly by the inclusion of eTIV across many metrics and ROIs, typically attenuating the degree of significance as well as the estimated slope. Hemispheric asymmetry was not affected by the presence or absence of eTIV in the model, as would be expected since eTIV does not differ between the pair of right and left ROIs. Importantly, controlling for the volume differences in the right and left ROIs for the 21 pairs of ROIs also did not impact the observed asymmetry in the mean DTI/NODDI values in these ROIs.

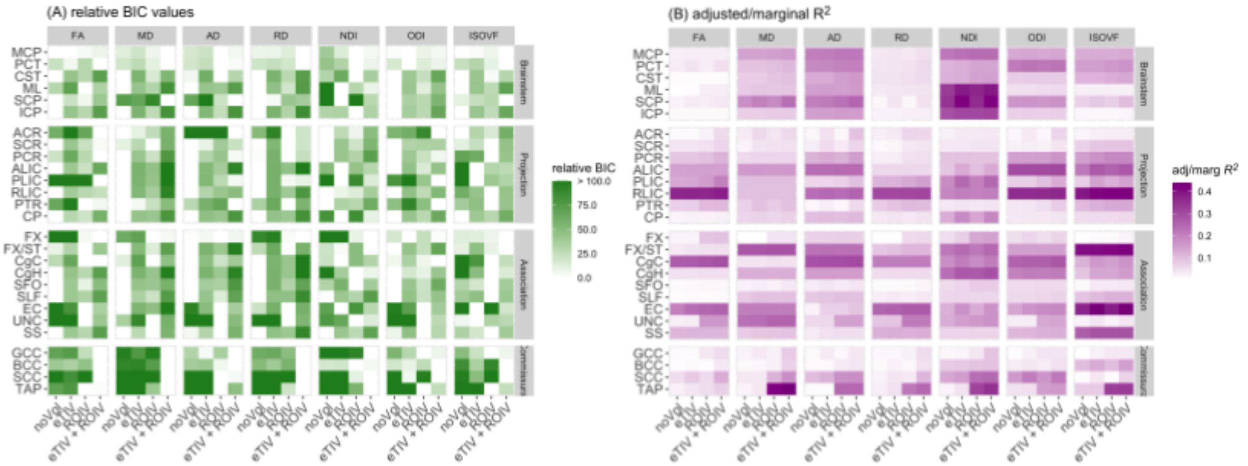

**Supplemental Figure 32. Comparisons of model quality for models with or without global and/or local volume correction for the mean DTI/NODDI values in JHU ROIs.**

Comparisons of (A) relative BIC values and (B) adjusted/marginal $R^2$ in four models with or without global and/or ROI volume as covariates are shown for each DTI/NODDI metric in the JHU ROIs. See supplemental text for the description of the four models, and see Table 2 of the main text for the full names of abbreviated ROIs.

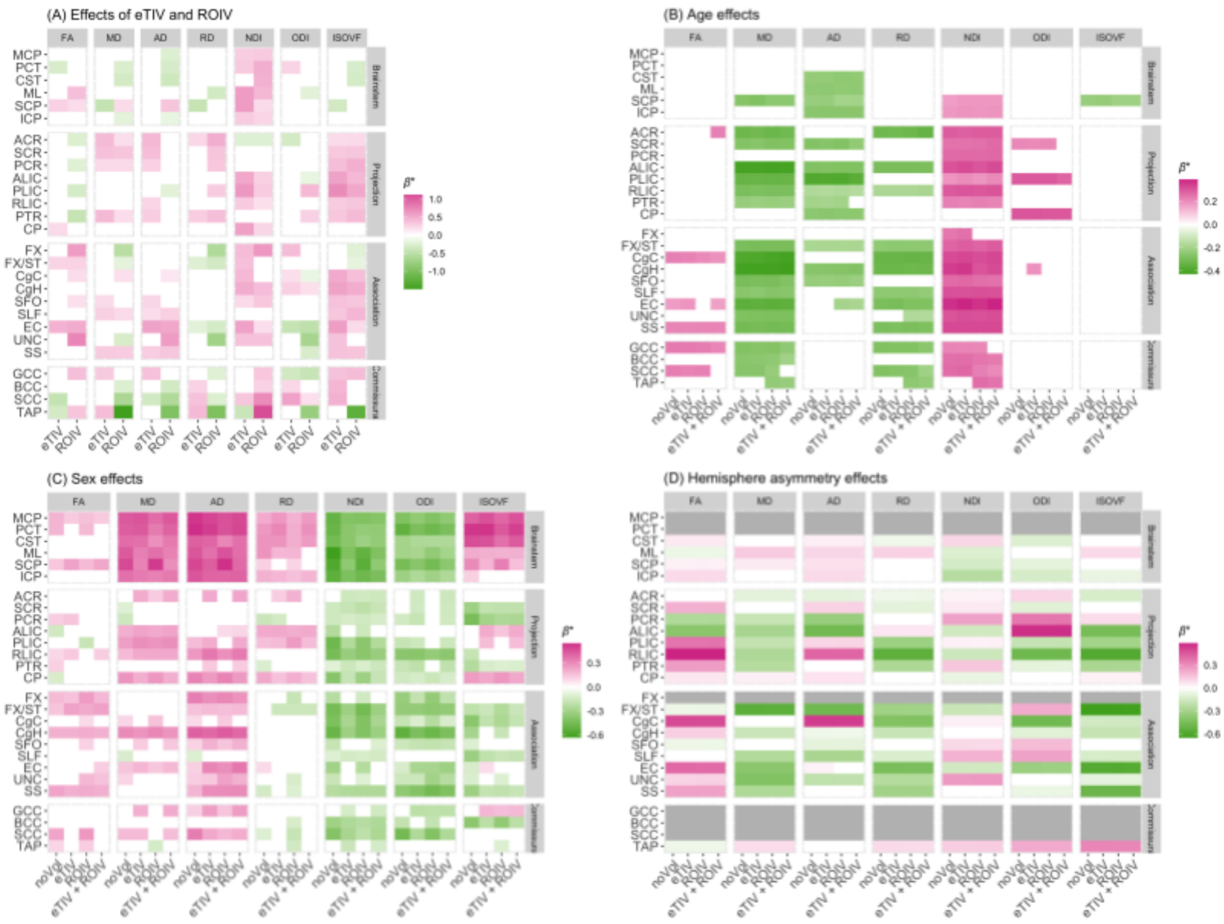

**Supplemental Figure 33. Effects of global or ROI volumes on the mean DTI/NODDI values and their impact on the estimated age, sex, and hemisphere effects.**

The standardized parameter estimates ($\beta^*$) for (A) eTIV or ROIV, (B) age, (C) sex, and (D) hemisphere effects are shown for each DTI/NODDI metric in the JHU ROIs. The eTIV/ROIV $\beta^*$ values were derived from models that included either eTIV or ROIV. For (B), (C) and (D), $\beta^*$ values are compared for each of the four models described in the supplemental text. Those that did not survive Bonferroni corrections for multiple comparisons were filtered out (set to 0) to facilitate comparisons within significant results. See Table 2 of the main text for the full names of abbreviated ROIs.
